# DNA stimulates SIRT6 to mono-ADP-ribosylate proteins within histidine repeats

**DOI:** 10.1101/2024.07.31.606047

**Authors:** Nicholas J. Pederson, Katharine L. Diehl

## Abstract

Sirtuins are the NAD^+^-dependent class III lysine deacylases (KDACs). Members of this family have been linked to longevity and a wide array of different diseases, motivating the pursuit of sirtuin modulator compounds. Sirtuin 6 (SIRT6) is a primarily nuclear KDAC that deacetylates histones to facilitate gene repression. In addition to this canonical post-translational modification (PTM) “eraser” function, SIRT6 can use NAD^+^ instead to “write” mono-ADP-ribosylation (mARylation) on target proteins. This enzymatic function has been primarily associated with SIRT6’s role in the DNA damage response. This modification has been challenging to study because it is not clear under what precise cellular contexts it occurs, only a few substrates are known, and potential interference from other ADP-ribosyltransferases in cells, among other reasons. In this work, we used commercially available ADP-ribosylation detection reagents to investigate the mARylation activity of SIRT6 in a reconstituted system. We observed that SIRT6 is activated in its mARylation activity by binding to dsDNA ends. We further identified a surprising target motif within biochemical substrates of SIRT6, polyhistidine (polyHis) repeat tracts, that are present in several previously identified SIRT6 mARylation substrates and binding partners. This work provides important context for SIRT6 mARylation activity, in contrast to its KDAC activity, and proposes that SIRT6 is a histidine mARyltransferase enzyme.

## Introduction

In mammals, there are seven sirtuin family members (SIRT1-7). Sirtuins are NAD^+^-dependent enzymes that possess deacylase and/or mono-ADP-ribosyltransferase activity.^1–3^ Sirtuins have been associated with longevity and implicated in numerous diseases, such as cancer and cardiovascular disease.^2,4–8^ For example, SIRT6 overexpression leads to lifespan extension in mice and drosophila,^9–11^ while SIRT6 deficiency leads to detrimental cellular and organismal effects.^6,8^ Interestingly, SIRT6 has been identified as a tumor suppressor in some cancers, and as tumorigenic in others^6,7,10,12–15^. Due to the roles of SIRT6 in numerous pathways and disease states, this enzyme has been studied extensively and this includes the pursuit of pharmacological inhibitors and activators.^16–18^

SIRT6 is perhaps best known as a histone deacetylase that acts at H3K9ac, H3K18ac, H3K27ac, and other sites to compact chromatin and aid in the silencing of target genes.^19–22^ SIRT6’s relatively low activity on acetylated peptides^21,23–25^ has led to an idea that SIRT6 alone possesses insufficient deacetylase activity.^26,27^ However, biochemical and structural data show that SIRT6 binds to the nucleosome with high affinity, rendering it active toward this substrate, ^19,20,1,28,24,22^ providing a rationale for its low activity on histone peptides compared to the context of a nucleosome substrate. In addition, SIRT6 has been shown to bind non-nucleosomal DNA, or so-called “free DNA,” and its ability to bind DNA has been linked to its role in the DNA damage response.^29–31^ Moreover, a recent paper reported that binding to free DNA increases SIRT6 activity on acylated peptides.^27^

In addition to its deacetylase function, SIRT6 can deacylate,^32^ defatty-acylate,^26,25^ or mono-ADP-ribosylate (“mARylate”)^33–38^ various substrates. Much like PARP1/2-catalyzed poly-ADP-ribosylation (pARylation), SIRT6-catalyzed mARylation is associated primarily with oxidative stress and DNA damage conditions.^33–36^ Its reported substrates to date are itself (specific site not identified), PARP1 (at K521),^33,34^ KAP1 (specific site not identified),^37^ KDM2A (at R1019),^35^ BAF170/SMARCC2 (at K312),^36^ and lamin A (specific site not identified).^38^ In the case of PARP1, SIRT6 was reported to mARylate K521, which is within the automodification loop of PARP1, to facilitate recruitment of PARP1 to DNA breaks and to activate its pARylation activity.^33,34^ SIRT6-catalyzed mARylation of the BAF chromatin remodeling complex subunit BAF170 was shown to recruit the remodeler complex to the enhancer region of an oxidative stress response gene to enable activation of this gene.^36^ Lamin A was first shown to activate SIRT6 deacetylation and mARylation activity in DNA repair,^39^ and then was later also shown to itself be mARylated by SIRT6 in cells to promote genome stability.^38^ Yet, relatively little is understood about what causes SIRT6 to MARylate substrates, what other substrates it might have, or what the function of this modification is. Here we demonstrate that SIRT6 binds to free DNA to induce mARylation of its known target BAF170^36^ as well as two novel substrates MeCP2 and NLK. We demonstrate that SIRT6 mARylates these proteins within histidine repeats. Taken together, our data provide mechanistic insights into how SIRT6 interacts with DNA to induce mARylation, identify SIRT6 as an enzyme that mARylates histidine, and provide a list of other potential SIRT6 mARylation substrates based on the poly-histidine (“polyHis”) motif.

### DNA induces SIRT6 mARylation of substrates containing histidine repeats

Since SIRT6 binds to free DNA ends and is recruited to damaged DNA,^29–31^ we first tested if the addition of DNA would affect SIRT6 mARylation activity. We started by measuring SIRT6 auto-activity^40^ with a biochemical ribosylation assay using a poly/mono-ADP-ribose antibody (Cell Signaling Technologies, CST) for detection. We prepared two full-length (2-355), human SIRT6 proteins, one with an N-Terminal 6xhistidine tag (“6xHis-SIRT6”) and another in which the 6xHis tag was proteolytically cleaved, rendering it identical to native human SIRT6 (“Cleaved-SIRT6,” residues S2-S355). For the DNA, we used 177-bp dsDNA derived from the Widom 601^41^ sequence (“ds601”). Strikingly, these enzymes respond differently to the presence of ds601 (**Figure 1A-B**). The addition of ds601 significantly increased 6xHis-SIRT6 auto-activity compared to the absence of ds601. For the Cleaved-SIRT6, the ds601 significantly decreased its auto-activity compared to the absence of ds601.This result was surprising since we initially reasoned the 6xHis tag might decrease SIRT6 activity given that the N-terminus of SIRT6 is near the active site and is necessary for catalytic activity.^21,42^

**Figure 1.**
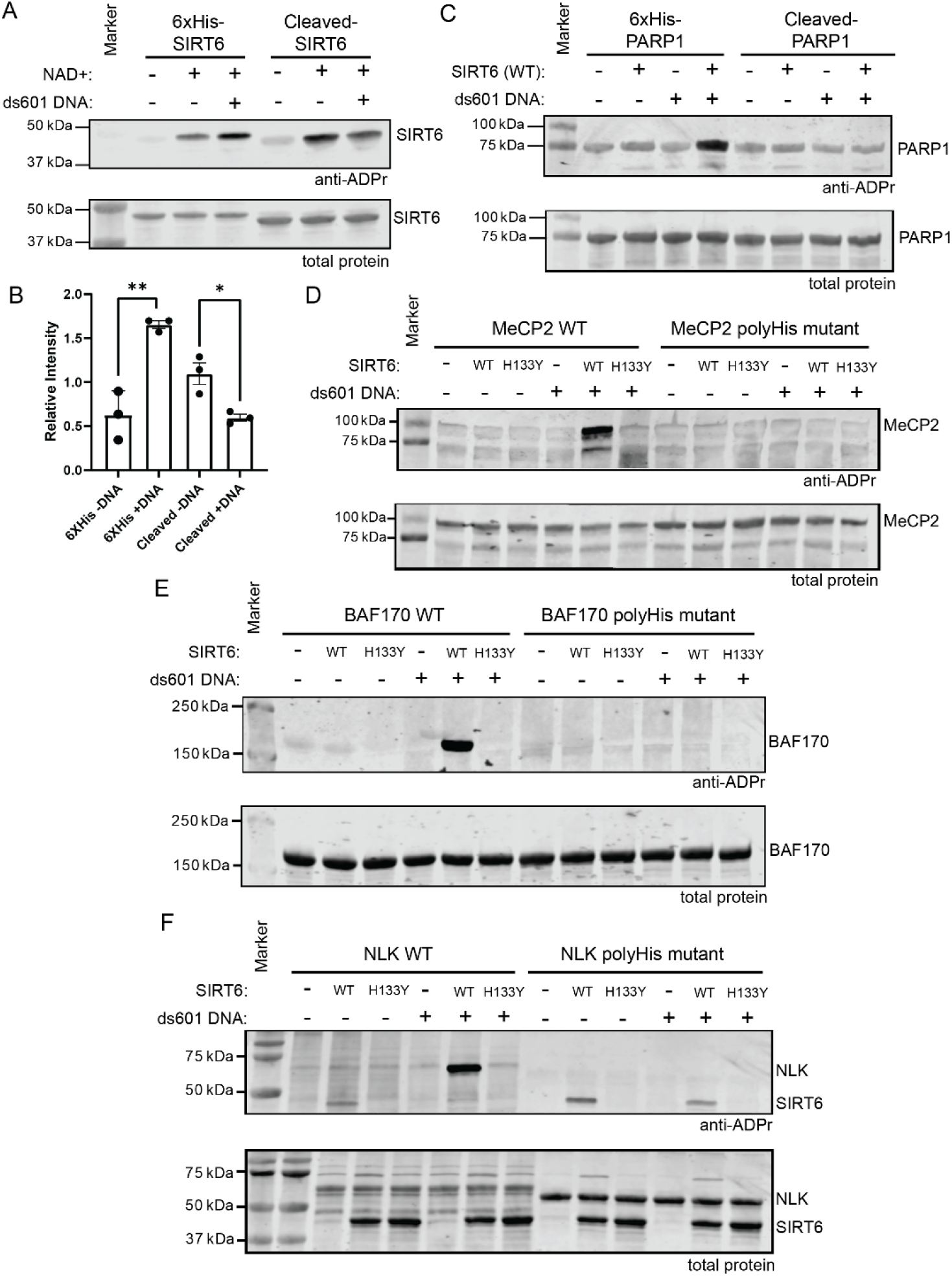
DNA stimulates SIRT6 to mARylate proteins containing polyHis. (A) Immunoblot analysis of 6xHis- and Cleaved-SIRT6 (2 µM) automARylation reactions with or without ds601 DNA (1 µM). The reactions were incubated at 37°C for 2 h +/-1 mM NAD^+^, n = 3. B) Quantification of the immunoblot data from panel A and replicate blots. The densitometry was performed in Image Studio (Licor) with normalization to the SIRT6 band in the total protein stain. An unpaired, two-tailed t-test was used: **p = 0.0034, *p = 0.0180, n = 3 independent replicate reactions, error bars show ± S.D. C) Immunoblot analysis of SIRT6 (10 µM) mARylation reactions with 6xHis-PARP1(2-655) or Cleaved-PARP1(2-655) (2 µM) as substrates. The reactions were incubated at 37°C for 2 h with 1 mM NAD^+^ and +/-1 µM ds601 DNA, n = 2. D-F) Immunoblot analyses of SIRT6 (2 µM) mARylation reactions with (D) MeCP2 WT or MeCP2 polyHis mutant, (E) BAF170-WT or BAF170 polyHis mutant, or (F) NLK-WT or NLK polyHis mutant as the substrate (all at 2 µM). SIRT6-H133Y is an inactive mutant that was included as a negative control.

The reactions in panels D and F were incubated at 37°C for 2 h and in panel E for 20 min with 1 mM NAD^+^ and +/-1 µM ds601 DNA, n = 2. All the immunoblots in this figure were performed using a poly/mono-ADP-ribose antibody (Cell Signaling Technologies #83732).

Auto-mARylation by SIRT6 was shown previously to be an intramolecular (or “in *cis*”) reaction,^40^ so we next asked if other substrates would be affected since the intermolecular (or “in *trans*”) mARylation mechanism could differ. SIRT6 mARylation of PARP1 was shown to activate PARP1 and promote DNA damage repair in cells.^33,34^ We generated the same hPARP1 fragment (PARP1(2-655)) that was used in the biochemical assays in the previous study.^34^ This truncation lacks the entire catalytic domain and cannot auto-PARylate and was reported to be mARylated by SIRT6 (at PARP1 K521). We made versions of this truncated PARP1 with and without the N-terminal 6xHis tag. We note that it was not described in the previous study if the PARP1(2-655) that was used contained a 6xHis tag or not.^34^ We only observed SIRT6 dependent modification of PARP1(2-655) that contains a 6xHis tag in the presence of dsDNA (**Figure 1C**).

Given the discrepancy between 6xHis-tagged constructs, we wondered if the exogenous histidine tag on SIRT6 and PARP1(2-655) could mimic a motif in endogenous SIRT6 substrates. PolyHis tracts are relatively uncommon in the human proteome compared to other single amino acid repeat tracts.^43,44^ Most polyHis-containing proteins are annotated as nuclear proteins and many are transcription factors.^43,45–47^ Interestingly, when we looked at the polyHis proteins in humans (**Data File S1**), we identified four proteins that are known to interact with SIRT6: BAF170 (also called SMARCC2)^36^, Lamin A/C,^38,39^ MeCP2,^24^ and YY1^48^ (**Table 1**). BAF170^36^ and Lamin A/C^38^ are two of the reported substrates of mARylation by SIRT6. MeCP2^24^ is reported to form a complex with SIRT6 in a DNA-dependent manner, and YYI^48^ is reported to form a complex with SIRT6 as well. Given this information, we next tested if SIRT6 would mARylate three different polyHis-containing proteins.

**Table 1.** SIRT6-associated and/or polyHis proteins with relevance to this study.

| Protein | UniProt | Association with SIRT6 | Previously reported to be MAR Target of SIRT6? | mARylated in this study by SIRT6? | PolyHis sequence | Number of total residues in the protein |
| --- | --- | --- | --- | --- | --- | --- |
| PARP1 | **P09874** | DNA damage<sup>33,34</sup> | Yes | No* | None | 1014 |
| MeCP2 | **P51608** | DNA damage<sup>24</sup> | No | Yes | 366-HHHHHHH-372 | 486 |
| BAF170 | **Q8TAQ2** | DNA damage<sup>36</sup> | Yes | Yes | 1117-HGHHHH-1122 | 1214 |
| NLK | **Q9UBE8** | None | No | Yes | 18 His in first 54 residues | 527 |
| LAMA | **W8QEH3** | Longevity<sup>38,39</sup> | Yes | Not Tested | 563-HHHH-566 | 572 |
| GATA6 | **Q92908** | DNA damage<sup>49</sup> | No | Not Tested | 324-HHHHHHHHHH-333 | 595 |
| YY1 | **P25490** | Longevity<sup>48</sup> | No | Not Tested | 70-HHHHHHHHHHH-80 | 414 |

\*PARP1(2-655) lacking a 6xHis tag was not mARylated by Cleaved-SIRT6 in our assay.

We chose MeCP2, BAF170, and nemo-like kinase (NLK) as substrates for mARylation assays with SIRT6. NLK was chosen as a polyHis protein with no prior association with SIRT6 or ADP-ribosylation. To avoid using a 6xHis tag, we used a GST tag to purify MeCP2 from *E. coli*, and we used a FLAG tag to purify BAF170 and NLK from *S. frugiperda* (Sf9). We purified the full-length MeCP2 (2-486), BAF170 (1-1214), and NLK (1-527) as well as mutants of each lacking the histidine repeats (the full amino acid sequences are provided in the **Supporting Information**). For MeCP2 and BAF170, the polyHis tracts were mutated to glycines. For NLK, which has two polyHis tracts in the N-terminus, we truncated the N-terminus (Δ2-55) and mutated H124/H125/H126 to alanine. In our assay, SIRT6 mARylates MeCP2, BAF170, and NLK in the presence of dsDNA and is unable to mARylate the polyHis mutants (**Figure 1D-F**). We used a catalytically inactive mutant (SIRT6-H133Y^42^) to confirm that the mARylation was dependent on the enzymatic activity of SIRT6 (**Figure 1D-F**). We note that BAF170 mARylation is obvious within a 20 min reaction (**Figure 1E**), while MeCP2 and NLK mARylation required a longer incubation time to be apparent (2 h, **Figure 1D and 1F**). We also note that for the assay with PARP1 (**Figure 1C**), we used 10 µM SIRT6 to be able to observe modification. For the reactions in **Figure 1D-F**, we used only 2 µM SIRT6. These observations imply that SIRT6 exhibits substrate preferences based on factors other than simply the presence of a polyHis sequence. Since PARP1(2-655) also binds to DNA ends,^50,51^ there is probably competition with the SIRT6 for occupation of the ends. We tested the same conditions for MeCP2 using a pan-ADP-ribose binding reagent (Millipore) that is derived from a bacterial macrodomain^52^ (**Figure S1**). The same results were obtained with both the antibody and macrodomain detection methods.^53^ These results support our initial finding that DNA stimulates SIRT6 to mARylate and that this happens on substrates containing histidine repeats. This also identifies MeCP2, NLK, and other polyHis proteins as potential new SIRT6 mARylation substrates.

### SIRT6 mARylation activity is activated by binding to DNA ends

SIRT6 is important during the DNA damage response and has been shown to promote non-homologous end joining (NHEJ) and homologous recombination (HR) in cells.^29,54^ SIRT6 has also been shown to independently arrive at DNA double strand breaks in cells.^29–31^ Previous work indicates that more than one SIRT6 molecule is able to occupy one molecule of DNA, and it binds as a dimer to the DNA end.^29,30^ Given this information, we performed a DNA titration to determine how the binding stoichiometry would affect SIRT6 activity. We used the ds601 DNA as before and MeCP2 as the substrate. The titration showed that increasing ds601 led to increased modification of MeCP2 by SIRT6 up to 500 nM ds601 (or 4:1 ratio of SIRT6:ds601) after which point higher concentrations of ds601 led to reduced SIRT6 activity (**Figure 2A**). If SIRT6 simply needed to be bound to DNA to achieve its activated state, we would expect SIRT6 activity to increase with increasing amounts of DNA until reaching a plateau once the SIRT6 was saturated with DNA. Our observation of a decrease in activity at lower SIRT6:DNA ratios (i.e., 2:1, 1:2, 1:4) suggests that dimerization is important for activation. We tested single-stranded 601 DNA (ssDNA) and observed a lower magnitude of activation compared to ds601 (**Figure 2B**). We next tested circular plasmid DNA and found that it did not activate mARylation by SIRT6 (**Figure 2C**), which is consistent with previous studies of SIRT6 DNA binding preferences.^27,29,30^ Altogether, these data suggest that SIRT6 binding at DNA ends is important for the activation of its MARylase activity and are consistent with a model of SIRT6 dimerization at dsDNA ends.^29,30^

**Figure 2.**
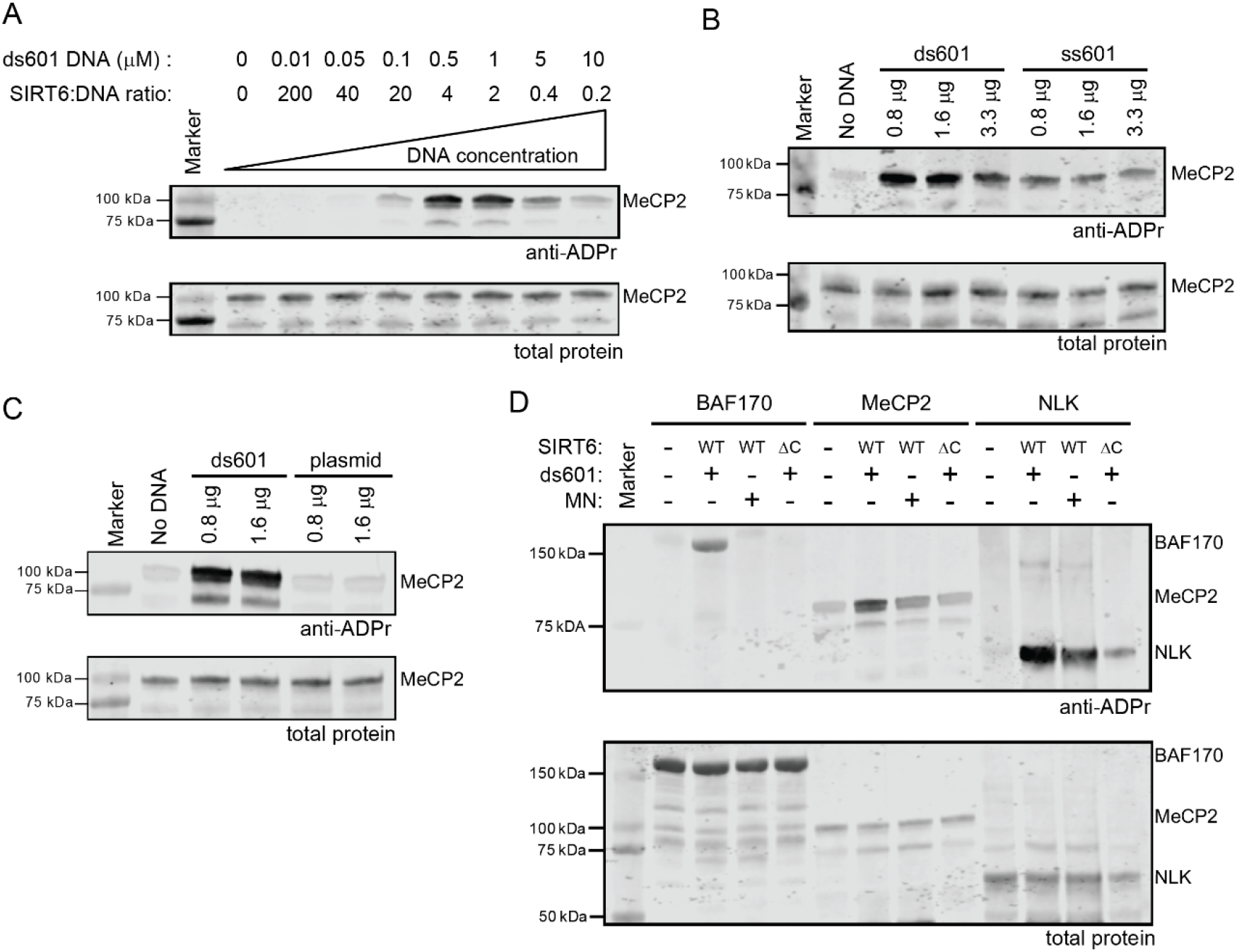
SIRT6 is activated in its mARylation activity by binding to DNA ends. A) Titration of ds601 in the MeCP2 (2 µM) mARylation assay by SIRT6 (2 µM), n = 2. B) Titration of ds601 and ss601 in the MeCP2 (2 µM) mARylation assay by SIRT6 (2 µM), n = 2. Note that there was no SIRT6 in the reaction for the “No DNA” lane. In terms of concentration, 0.8 µg = 0.5 µM ds601, 1.6 µg = 1 µM ds601, and 3.3 µg = 2 µM ds601. C) Immunoblot analysis of mARylation by SIRT6 (2 µM) of MeCP2 (2 µM) in the presence of plasmid DNA (pET28a), n = 2. Note that there was SIRT6 in the reaction for the “No DNA” lane. D) Immunoblot analysis of mARylation assays of SIRT6-WT or -ΔC (2 µM) with ds601 or WT mononucleosome (1 µM) with BAF170, MeCP2, or NLK (2 µM) as the substrate. The BAF170 reactions were incubated for 20 min. The poly/mono-ADP-ribose antibody (Cell Signaling Technologies #83732) was used for all the blots shown here. All the mARylation assays were incubated at 37°C for 2 h unless otherwise noted.

Various point mutations in the catalytic domain have been reported to decrease SIRT6’s DNA binding affinity.^29^ The C-terminal extension (residues 269-355) of SIRT6 is reported to contribute significantly to DNA binding.^28^ C-terminal truncations of SIRT6 have reduced deacetylase activity on nucleosomes (**Figure S2A**) but deacetylate peptides in our (**Figure S2B**) and others’ hands.^28,42^ We compared the binding of SIRT6-WT and a SIRT6 C-terminal truncation (SIRT6-Δ298-355, abbreviated as “SIRT6-ΔC”) to the ds601 DNA by gel shift assay. Our results recapitulated prior findings^27,28^ in that both proteins bind the ds601 DNA, but the binding of the SIRT6-ΔC is weaker than the WT. The SIRT6-WT has a higher affinity for the DNA and exhibits a larger shift beginning at the 2.0 µM SIRT6 concentration that is not observed with the SIRT6-ΔC up to 7 µM protein (**Figure S2C**). The multiple banding pattern in the SIRT6-WT assay is consistent with a higher SIRT6:DNA stoichiometry than 1:1. We then compared the mARylation activity of SIRT6-WT to -ΔC (both without a 6xHis tag) and found that the truncation retains auto-activity (**Figure S2D**). However, the truncation is impaired in its mARylation of BAF170, MeCP2, and NLK compared to SIRT6 WT (**Figure 2D**). SIRT6-ΔC has no apparent activity toward the substrates BAF170 and MeCP2. However, the truncation is still able to modify NLK, albeit to a much lower extent than the full-length enzyme.

SIRT6 deacetylates histone H3 lysine 9 acetylated (“H3K9ac”) mononucleosomes (**Figure S2A**).^24^ Recent cryo-EM structures of the nucleosome-bound SIRT6 show that it binds in a specific mode to the nucleosome surface, positioning it to place the H3 tail in the active site.^19,20,22^ While it was shown before that SIRT6 does not MARylate non-nucleosomal histones,^40^ we tested if SIRT6 could MARylate histone H3 in the context of nucleosomes (**Figure S2E**). We observed a faint, SIRT6-dependent band above histone H3 in the ADPr blot. To determine if this band could be MARylated H3, we probed with an H3 antibody (binding to a C-terminal epitope) and observed no indication that the new band in the ADPr blot was H3. Furthermore, MARylated H3 that was prepared semi-synthetically in another study (H3S10ADPr) showed a much smaller mobility shift^55^ than the band we observed in our ADPr blot. Thus, we conclude that SIRT6 does not MARylate H3 in the nucleosome context.

Next, we wanted to know if nucleosomes could activate SIRT6 to MARylate our previously characterized substrates. We note that the nucleosomes we used have 15 bp overhangs on either side of the Widom 601 sequence, which are sufficient to activate PARP1.^50^ For BAF170, there was no detectable activation of SIRT6. For MeCP2 and NLK, there was some activation, although it was attenuated compared to ds601 (**Figure 2D**). These results support the previous biochemical and structural data that nucleosome-bound SIRT6 is primed for lysine deacetylation. Based on our data, the nucleosome-bound state is not conducive to SIRT6’s mARylation activity in the way that the DNA end-bound state is, supporting context-dependent activities of SIRT6.

### Histidine is mARylated by SIRT6

Our data suggest that SIRT6 mARylates histidine repeats since mutation of the histidine repeats completely abolishes SIRT6-dependent modification (**Figure 1**). MARylation has been identified on different nucleophilic amino acid including glutamate, aspartate, serine, lysine, arginine, and cysteine and there are known enzymes responsible for these modifications.^56^ Diphtheria toxin (DT) and *Pseudomonas* exotoxin A (ETA) irreversibly mARylate a diphthamide analog of histidine in elongation factor-2 (EF-2) to halt protein translation.^57,58^ However, mARylation of histidine itself has only recently been reported as occurring in human cells.^59,60^ A group subsequently synthesized mARylated histidine-containing peptides (or a triazole mimic thereof) as tools to study this modification.^59,61^ There are no human enzymes yet shown to specifically mARylate histidine.

To further test our hypothesis that SIRT6 is mARylating histidine, we created an MeCP2 construct with the histidine repeats mutated to arginine (polyArg), which would provide a poly-cationic alternative to polyHis. Since arginine is known to be mARylated by some ADP-ribosyltransferases,^35,62^ we reasoned that SIRT6 would be able to modify this MeCP2 construct if the modification was not histidine specific. Furthermore, SIRT6 was reported to mARylate one of its reported substrates, KDM2A, on an arginine residue.^35^ However, SIRT6 does not mARylate the polyArg MeCP2 (**Figure 3A**). Moreover, SIRT6 will not mARylate the lysine- and arginine-rich histone H3 tail of a nucleosome (**Figure S2E**) to which it is known to bind with high specificity.^19,20,28^ We next titrated a polyHis peptide (HHHHHHGGG) into the MeCP2 mARylation assay and observed that the peptide inhibits MeCP2 mARylation by SIRT6 as well as SIRT6 auto-mARylation (**Figure S3A**). This finding is consistent with the polyHis peptide competing for the SIRT6 active site, rather than binding to an allosteric site.

**Figure 3.**
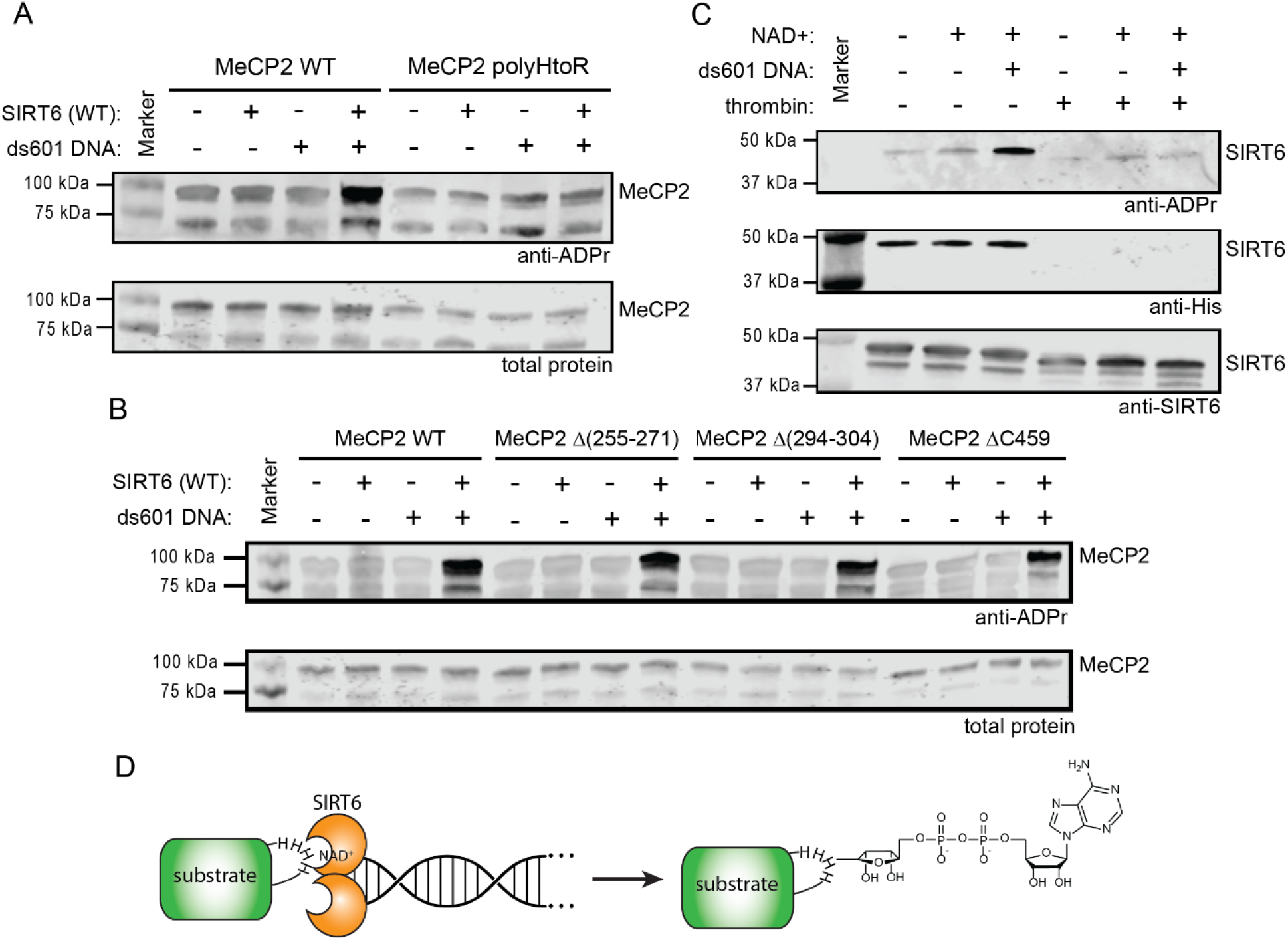
SIRT6 mARylates Histidine. A) Immunoblot analysis of SIRT6 (2 µM) mARylation reactions with MeCP2-WT or -polyHtoR mutant (2 µM) with or without 1 µM ds601 (37°C for 2 h), n = 2. B) Immunoblot analysis of SIRT6 (2 µM) mARylation reactions with MeCP2-WT, (Δ255-271), (Δ294-304), or ΔC459 (2 µM) with or without 1 µM ds601 (37°C for 2 h), n = 2. C) Immunoblot analysis of 6xHis-SIRT6 (2 µM) mARylation (1 µM ds601, 1 mM NAD+, 37°C for 20 min) before and after cleavage by thrombin (1 unit), n = 2. D) Schematic of SIRT6 mARylation activation by binding to dsDNA ends toward polyHis substrates. The poly/mono-ADP-ribose antibody (Cell Signaling Technologies #83732) was used for all the blots shown here.

We analyzed the mARylated MeCP2 by mass spectrometry to directly observe the modified residues. However, the polyHis tract was in the 15% of the MeCP2 sequence that was not covered in the analysis (**Figure S3B, Data File S2**). The analysis found relatively low levels of mARylation on three MeCP2 residues: D260, T299, and S464. We generated MeCP2 deletions (Δ255-271, Δ294-304, and a C-terminal deletion Δ459) that correlated to the entire tryptic peptides that were identified as being mARylated, in case the specific residue assignment was not accurate within the peptide. These proteins were all robustly mARylated by SIRT6 (**Figure 3B**), unlike the polyHis mutant (**Figure 1D**). This finding indicates that D260, T199, and S464 are not the predominant sites of SIRT6-catalyzed mARylation in MeCP2.

We next utilized our initial finding that auto-mARylation of the 6xHis-tagged SIRT6 is stimulated by DNA (**Figure 1A**). This SIRT6 construct has a thrombin cleavage site (“LVPR|GS”) between the 6xHis tag and the beginning of the SIRT6 sequence. We performed the mARylation reaction followed by cleavage with thrombin. There is a large decrease in the ADP-ribosylation signal on the SIRT6 subsequently treated with thrombin compared to the SIRT6 not treated with thrombin (**Figure 3C**). The ADP-ribosylation signal that is retained after treatment with thrombin can be attributed to the automodification in SIRT6 that is polyHis independent (**Figure 1A**) and which is presumed to be an intramolecular reaction^40^ and distinct from the DNA-induced histidine modification. We also looked at the sequences surrounding the polyHis tracts in the three substrates that we tested and in the full set of 129 human polyHis proteins (**Data File S1**), and there are no consensus motifs present in terms of other amino acids in the flanking sequences that could act as acceptor residues for the mARylation (**Figure S3C**,**D**). Even so, serine was one of the most common amino acids near the polyHis tracts, and it appears near the polyHis tract in several of the substrates we tested (**Figure S3C**). To test if SIRT6 could be modifying a serine instead, we used the serine-specific ADP-ribosyl-hydrolase ARH3.^63–65^ Incubation of ARH3 after the ADP-ribosylation reaction of MeCP2 resulted in no difference in the degree of modification of the MeCP2 (**Figure S3E**), indicating that the ADP-ribosylation is not on serine. We note that ADP-ribosylation of the SIRT6 (with no 6xHis tag) was diminished in the presence of ARH3, suggesting that the SIRT6 automodification reaction occurs, at least in part, on serine. Altogether, our results support that the DNA-dependent mARylation by SIRT6 occurs within the polyHis tract.

## Discussion

Sirtuins exhibit multiple enzymatic functions, and it has remained unclear to what extent the mARtransferase activity that some sirtuins display is a “side reaction” to their canonical deacylase activity.^66^ Here we show that SIRT6 mARylation activity is robustly activated by binding to DNA ends toward substrates containing a polyHis tract (**Figure 3D**). These findings are consistent with previous work showing that SIRT6 is rapidly recruited to DNA damage sites^29–31^ and that its mARtransferase activity is associated with oxidative stress and DNA damage conditions.^33–36^ We observed that the nucleosome-bound state of SIRT6 is not conducive to mARylation, suggesting that SIRT6 performs deacylation and mARylation under distinct circumstances. It is still unclear what drives SIRT6 to be recruited to DNA breaks in cells. It was shown that SIRT6 arrives at DNA breaks independently of MRE11, Ku80, or PARP1/2,^29^ suggesting that SIRT6 directly recognizes breaks similarly to PARP1/2. However, SIRT6 binds to the nucleosome with low nanomolar affinity^19,28^ and to “free” DNA ends with high nanomolar^30^ to low micromolar affinity.^29^ In contrast, PARP1 (zinc fingers F1 and F2) binds to a dsDNA break with low nanomolar affinity and to circular or unbroken DNA with low micromolar affinity.^51^ Given this information, it is unclear how SIRT6 could spontaneously reorganize from nucleosomes to DNA ends on its own. Further investigation is needed to understand this mechanism.

When we initially observed that SIRT6 modified proteins with a polyHis tract, we were unsure of the significance of this reaction given that the first substrates we tested had an exogenous polyHis motif (i.e., 6xHis-SIRT6 and 6xHis-PARP1(2-655)). When we looked at polyHis proteins in the human proteome, we noted that lamin A/C and BAF170 were previously shown to be mARylated by SIRT6 in cells.^36,38^ For the lamin A/C study, the site of mARylation was not determined. However, an earlier study^39^ showed that a peptide comprising lamin A residues 567-596 was sufficient to bind to SIRT6 and that those 30 residues were necessary for association with SIRT6 (among the peptides comprising up to residues 567-646 that were tested in the study). Intriguingly, this peptide (567-596) immediately follows the polyHis tract (residues 563-566) in lamin A suggesting the polyHis track could be the region of mARylation and the following region is used for binding of SIRT6 to lamin A. For BAF170, K312 was identified by mass spectrometry as being mARylated by SIRT6 in mouse cells and tissue,^36^ but the full datasets are not available for further analysis. Two other proteins that were reported to bind to SIRT6 have polyHis tracts, MeCP2 and YY1. Altogether, these data motivated us to ask if a polyHis tract is an endogenous SIRT6 target sequence.

Based on this evidence, we biochemically tested BAF170, MeCP2, and NLK as described herein and found that all three were modified by SIRT6 in a dsDNA- and polyHis-dependent manner. While we tested a small subset of polyHis proteins in this study, this polyHis motif now provides a candidate list of other potential SIRT6 mARylation substrates. By demonstrating that SIRT6 is a histidine mARyltransferase, we provide a new avenue of study for understanding histidine ADP-ribosylation. This work also provides important clues about the context in which SIRT6 mARylation can occur that will be useful in studying this activity in cells. The regulatory purpose of SIRT6-catalyzed mARylation of these polyHis tracts is an area for further investigation. The histidine repeat tracts occur almost exclusively in disordered regions and often occur along with other repetitive tracts (e.g., glutamine, proline).^43^ It was shown that mutation or removal of polyHis tracts inhibits localization of the protein to nuclear speckles and inhibits phase separation.^43,45^ Thus, this modification could modulate the liquid-liquid phase separation behavior of these polyHis proteins. Delineating the full scope of SIRT6’s biochemistry will be critical in delineating its many cellular roles. This understanding is of particular importance in disease states in which SIRT6 or its substrates behave aberrantly and in considering the therapeutic potential of small molecule inhibitors or activators of SIRT6 which could possibly be used to modulate the enzymatic duality of SIRT6.^16–18^

### Experimental Procedures

#### Materials

All salt, buffers, and other chemicals were obtained from Fisher Scientific unless otherwise noted. Chemicals used for solid-phase peptide synthesis (SPPS) were obtained from ChemImpex International unless otherwise noted. Other reagents for cloning, protein purification, and the biochemical assays are listed in **Table 2**. SIRT6 was a gift from Cheryl Arrowsmith (Addgene plasmid #41565; http://n2t.net/addgene:41565;RRID:Addgene_41565). pFastBac1 Flag Baf170 was a gift from Robert Kingston (Addgene plasmid #1955; http://n2t.net/addgene:1955 ; RRID:Addgene_1955).^67^ The pACeBac1-PARP1 plasmid was generated in a previous study.^50^ The MeCP2 and NLK genes were synthesized as GeneBlocks by Integrated DNA Technologies (IDT).

**Table 2.** Materials.

| Reagent | Manufacturer | Catalog number |
| --- | --- | --- |
| Q5 High-Fidelity DNA Polymerase | New England Biolabs (NEB) | M0491 |
| Dpn1 | New England Biolabs (NEB) | R0176 |
| NEBuilder HiFi DNA Assembly Master Mix | New England Biolabs (NEB) | E2621 |
| T4 PNK | New England Biolabs (NEB) | M0201 |
| T4 DNA ligase | New England Biolabs (NEB) | M0202 |
| 6x NEB Purple DNA loading dye | New England Biolabs (NEB) | B7025 |
| 100 bp ladder | New England Biolabs (NEB) | N0551 |
| 1 kb ladder | New England Biolabs (NEB) | N3232 |
| DH5α Mach1 competent cells | Invitrogen | C862003 |
| MAX Efficiency™ DH10Bac Competent Cells | Gibco | 10361012 |
| Rosetta(DE3) Competent cells- Novagen | Sigma | 70954 |
| QG buffer | Qiagen | 19063 |
| Miniprep buffers and columns | Qiagen | 27104 |
| Protease Inhibitor Mini Tablets | Pierce | A32953 |
| High Affinity Ni-Charged Resin | GenScript | L00223 |
| Glutathione Resin | GenScript | L00206 |
| Anti-DYKDDDDK (FLAG) G1 Affinity Resin | GenScript | L00432 |
| DYKDDDDK Peptide | GenScript | RP10586 |
| HiLoad Superdex 16/600 200pg column | Cytiva | 28989335 |
| Superdex 200 Increase 10/300 GL | Cytiva | 28990944 |
| Thrombin | Cytiva | 27084601 |
| Dialysis Tubing | Thermofisher | 88244 |
| XBridge BEH C18 columns | Waters | 186003624,<br>186008193,<br>186003673 |
| NAD<sup>+</sup> | New England Biolabs (NEB) | B9007S |
| Rabbit anti-poly/mono-ADP-ribose | Cell Signaling Technologies (CST) | 83732* (Lot 5) |
| Rabbit anti-H3K9ac | Cell Signaling Technologies (CST) | 9649 (Lot 13) |
| Mouse anti-His Tag | Cell Signaling Technologies (CST) | 2366 (Lot 14) |
| Rabbit anti-pan-ADP-ribose binding reagent | Millipore | MABE1016 (Lot 3595120) |
| Mouse anti-H3 | Cell Signaling Technologies (CST) | 3638 (Lot 11) |
| Rabbit anti-SIRT6 | Cell Signaling Technologies (CST) | 12486 (Lot 3) |
| IRDye 680CW goat anti-mouse | LI-COR | 926-68070 |
| IRDye 800CW goat anti-rabbit | LI-COR | 926-32211 |
| Revert 700 total protein stain | LI-COR | 926-11010 |
| MES 20x buffer | Bio-Rad | 1610789 |
| Dual color ladder | Bio-Rad | 1610374 |
| PVDF membrane | Bio-Rad | 1620177 |
| SIRT6 Inhibitor (OSS_128167) | Med Chem Express | HY-107454 |

\*The #83732 mAb is discontinued and has been replaced by #89190.

#### Cloning

Q5 High-Fidelity DNA Polymerase (NEB) was used for all PCR amplification steps. All primers were obtained from the DNA/Peptide Core at the University of Utah. PCR products were treated with Dpn1 (NEB), and linear products were purified via a 1% agarose gel, extracted with QG buffer (Qiagen), and purified by spin column (Qiagen). NEBuilder HiFi DNA Assembly Master Mix (NEB) was used as described by the manufacturer. For blunt-end ligations, T4 PNK (NEB) and T4 DNA ligase (NEB) were used as indicated by the manufacturer. Reactions were transformed into DH5α Mach1 cells (Invitrogen) and plated on antibiotic-containing agar to select colonies for sequence verification. Plasmids were isolated from liquid cultures using Qiagen Miniprep buffers and spin columns as directed by the manufacturer. All plasmids were sequence verified by GENEWIZ (Aventa Life Sciences). Amino acid sequences are listed in the **Supporting Information**. Bacmids were prepared according to the MultiBac protocol from Geneva Biotech.

The Sirt6 C-terminal extension was added to the Sirt6 gene in the plasmid #41565 (pET28a-LIC vector) to generate the full-length Sirt6 gene (aa 1-355). All Sirt6 mutants were then subcloned from this template. PARP1(2-655) was subcloned from the pACeBac1-PARP1(full-length) plasmid into the pET28a-LIC vector. The pFastBac1 Flag Baf170 #1955 was used as is for protein expression. The MeCP2 and NLK GeneBlocks were assembled into the pET28a-LIC and pACeBac1 vectors, respectively.

#### Protein Expression

*E. coli*: Plasmids were transformed into BL21 Rosetta(DE3) cells (Millipore) and grown at 37 °C in LB Miller (Fisher) to OD600 = 0.6 with shaking at 180 rpm with the appropriate antibiotic. After reaching an OD = 0.6 the temperature was changed to 18 °C, and cells were induced with 0.5 mM IPTG and grown for 18 h before harvesting and storage at -80°C. SIRT6, MeCP2, PARP1(2-655) ARH3, and histones were grown in *E. coli*.

Sf9: Bacmids were transfected into Sf9 cells to produce baculovirus and the desired protein as previously described.^50^ BAF170 and NLK were grown in Sf9 cells.

#### Protein Purification

Coomassie-stained SDS-PAGE gels are shown in **Figure S4**. for the purified proteins.

##### MeCP2

Cell pellets were thawed on ice for 1 h followed by resuspension in lysis buffer (50 mM Tris pH 8.0, 500 mM NaCl, 2 mM MgCl_2_, 5 mM DTT, 1 mM PMSF). Cells were lysed via sonication and clarified by centrifugation at 20,000 x g for 30 min. Soluble lysate was applied to equilibrated glutathione resin from GenScript (5 mL of resin per liter of growth) and allowed to batch bind with nutation at room temperature for 2 h. The resin was washed with 15 column volumes of wash buffer (50 mM Tris pH 8.0, 500 mM NaCl, 2 mM MgCl_2_, 2 mM DTT) and eluted with 30 mL of elution buffer (50 mM Tris pH 8.0, 500 mM NaCl, 2 mM MgCl_2_, 2 mM DTT, 10 mM reduced glutathione). Proteins were either dialyzed overnight against storage buffer or injected on a HiLoad Superdex 16/600 200pg column (Cytiva) for elution in storage buffer (storage buffer: 50 mM Tris pH 8.0, 100 mM NaCl, 10 % glycerol, 2 mM DTT). The fractions were pooled, and protein was concentrated down to the desired concentration as determined by absorbance at 280 nm and an extinction coefficient calculated by ProtParam (https://web.expasy.org/protparam/) and stored at -80°C.

Note:Addition of 5 mM DTT in the lysis/binding buffer with the addition of excess of GST resin and longer binding at room temperature allowed for a better protein yield.

##### Proteins cleaved with ULP1 (SIRT6 WT, PARP1(2-655), ARH3)

Cell pellets were thawed on ice for 1 h followed by resuspension in lysis buffer (40 mM Tris, 500 mM NaCl, 10 mM MgCl2, 5 mM β-ME, and 1 mM PMSF, pH 7.5 for SIRT6, 8.0 for PARP1 (2-655)). Lysis buffer for PARP1 (2-655) was supplemented with Protease Inhibitor Mini Tablets (Pierce). Cells were lysed via sonication and clarified by centrifugation at 20,000 x g for 30 min. Soluble lysate was applied to lysis buffer-equilibrated nickel resin (Genscript) and allowed to batch bind with nutation at 4°C for 1 h. The resin was washed with 100 mL lysis buffer per liter of growth supplemented with 30 mM imidazole. Proteins were eluted using 25 mL of lysis buffer supplemented with 300 mM imidazole. Then 25 µL of Ulp1 was added to the elution. The solution was transferred to dialysis tubing (3.5k MWCO, Thermofisher) and allowed to dialyze overnight against lysis buffer containing no imidazole or PMSF at 4°C with stirring. The next morning, the buffer was replaced with fresh dialysis buffer and allowed 1 h of dialysis. The solution was removed from the dialysis tubing and filtered through 0.45 µm filter to remove precipitation before being applied to equilibrated nickel resin to bind the Ulp1 and SUMO tags (reverse nickel). The solution was nutated with nickel resin for 15 min at 4°C. The flow through was obtained and concentrated down to 2 mL for injection on a HiLoad 16/600 Superdex 200 pg (Cytiva) and eluted in storage buffer (50 mM Tris, 100 mM NaCl, 1 mM MgCl2,10 % glycerol, 2 mM DTT, pH 7.5 for SIRT6 and ARH3 and 8.0 for PARP1 2-655). The fractions were pooled and concentrated down to the desired concentration as determined by absorbance at 280 nm and extinction coefficient calculated by ProtParam (https://web.expasy.org/protparam/) and stored at -80°C.

Note: The reverse nickel resin volumes and incubation times need to be empirically determined. The cleaved SIRT6 appears to non-specifically bind the nickel resin so too much resin and/or time will decrease yield, whereas too little resin/time will lead to incomplete removal of the Ulp1 and SUMO.

##### TEV cleavage (SIRT6-H133Y and SIRT6-ΔC298)

The same method was used as for the Ulp1 cleavage except 25 µL of TEV protease was added.

Note:We do not suggest using TEV to cleave SIRT6. In our experience, around 95% of the SIRT6 crashed out during the TEV cleavage and reverse nickel, and there was incomplete cleavage. Also, the SEC cannot separate cleaved versus uncleaved SIRT6, leading to a mixture of both proteins. The purification of the SIRT6 with SUMO/Ulp1 worked much better for us.

##### Nickel purification with no cleavage (6XHis SIRT6 WT and 6XHis PARP1 2-655)

Cell pellets were thawed on ice for 1 h followed by resuspension in lysis buffer (40 mM Tris, 500 mM NaCl, 10 mM MgCl_2_, 5 mM β-ME, and 1 mM PMSF, pH 7.5 for SIRT6, 8.0 for PARP1 (2-655)). Lysis buffer for PARP1 1-655 was also supplemented with Protease Inhibitor Mini Tablets (Pierce). Cells were lysed via sonication and clarified by centrifugation at 20,000 x g for 30 min. Soluble lysate was applied to equilibrated nickel resin and allowed to batch bind with nutation at 4°C for 1 h. The resin was washed with 100 mL lysis buffer per liter of growth supplemented with 30 mM imidazole. Proteins were eluted using 25 mL of lysis buffer supplemented with 300 mM imidazole. Eluant was concentrated down to about 2 mL for injection on the HiLoad 16/600 Superdex 200 pg (Cytiva) and eluted in storage buffer (50 mM Tris, 100 mM NaCl, 1 mM MgCl2, 10% glycerol, 2 mM DTT, pH 7.5 for SIRT6 and 8.0 for PARP1 2-655). The fractions were pooled and concentrated down to the desired concentration as determined by absorbance at 280 nm and extinction coefficient calculated by ProtParam (https://web.expasy.org/protparam/) and stored at -80°C.

##### FLAG Purifications

BAF170-WT, BAF170-4XHtoG, NLK WT, NLK Δ(2-54) H(78,79,80)toA were all FLAG purified according to the protocol below (adapted from^68^).

*Buffer A:* 20 mM HEPES, 10 mM KCl, 0.1 mM EDTA, 0.1 mM EGTA, 1 mM DTT, Pierce Protease Inhibitor Mini Tablets (1 per 10 mL), and 0.5 mM PMSF

*Buffer B:* 20 mM HEPES, 400 mM KCl, 1 mM EDTA, 1 mM EGTA, 10% glycerol, 1 mM DTT, Pierce Protease Inhibitor Mini Tablets (1 per 10 mL), and 0.5 mM PMSF

*Buffer BC 0*: 20 mM HEPES, 0.2 mM EDTA, 10% glycerol, 1 mM DTT, 0.2 mM PMSF *Buffer BC 100*: 20 mM HEPES, 100 mM KCl, 0.2 mM EDTA, 10% glycerol, 1 mM DTT, 0.2 mM PMSF

*Buffer BC 300*: 20 mM HEPES, 300 mM KCl, 0.2 mM EDTA, 10% glycerol, 1 mM DTT, 0.2 mM PMSF

*BAF170 WT and 4XHtoG Storage Buffer:* 20 mM HEPES pH 7.0, 100 mM KCL, 10% glycerol, 1 mM DTT

*NLK WT and NLK Δ(2-54) H(78,79,80)toA Storage Buffer*: 20 mM HEPES pH 7.5, 100 mM KCL, 10% glycerol, 1 mM DTT

Buffers were prepared at pH 7.0 for BAF170 constructs and at pH 7.5 for NLK constructs. Purifications were from 1 liter of growth from Sf9 cells, stored at -80°C prior to use. Cell pellets were thawed on ice for 1 h followed by resuspension in 10 mL of buffer A and allowed to swell on ice for 15 min. 625 µL of 10% IGEPAL CA-630 per 10 mL of buffer A was added, and the cells were vortexed briefly for lysis. Nuclei were pelleted at 4 °C for 30 s at 17,000 x g and the supernatant was removed. The nuclei were washed once with 10 mL of buffer A and were pelleted at 4 °C for 30 s at 17,000 x g. The resulting nuclear pellet was resuspended in 10 mL of cold nuclear extraction buffer B. Nuclear pellets were then incubated with nutation for 15 min at 4 °C. The supernatant (nuclear extract) was removed and diluted 2-fold with buffer BC-0. This solution was pelleted at 4 °C for 10 min at 17,000 x g to remove precipitation. Soluble nuclear extract was then applied to 1 mL of FLAG resin (GenScript) and incubated for 1 h at 4 °C with nutation. FLAG beads were washed once with 10 bed volumes of buffer BC-100, once with 10 bed volumes of buffer BC-300, and once more with 10 bed volumes of buffer BC-100. The protein of interest was eluted by 3 successive 1-mL washes with buffer BC-100 supplemented with 0.25 mg/mL FLAG peptide (Genscript) for 20 min at 4 °C with nutation. Elutions were combined and either dialyzed overnight against storage buffer or injected on the HiLoad Superdex 16/600 200 pg (Cytiva) and eluted in storage buffer. The proteins were concentrated down to the desired concentration as determined by absorbance at 280 nm and extinction coefficient calculated by ProtParam (https://web.expasy.org/protparam/) and stored at -80°C.

##### Histones

The histones (full-length H3, H4, H2A, and H2B and 6xHis-SUMO-H3A15C) were purified as described previously.^50^

#### Peptide and Histone Semi-Synthesis

The H3(1-14)K9ac-MESNa thioester, H3(5-13)K9ac-amide, and polyHis peptides (sequences below) were synthesized by SPPS and purified by RP-HPLC (Waters C18 column) as described previously.^50^ The thioester peptide was synthesized on a hydroxy-trityl resin (Chemmatrix) functionalized with hydrazine. The amide-terminated peptides were synthesized on a Rink amide resin (Chemmatrix).

The H3K9ac histone was assembled by native chemical ligation between the H3(1-14)K9ac-MESNa thioester peptide and the H3A15C protein followed by desulfurization and RP-HPLC purification as described previously.^50^ The HPLC and mass spectrometric characterization of the three peptides and the ligated, desulfurized histone are shown in **Figure S5 and S6A**. The RP-HPLC characterization was performed on an Agilent 1260 Infinity HPLC system using a Waters C18 column (#186003624) in a water/acetonitrile/0.1% TFA solvent system. The mass spec analysis was performed on a Waters Acuity QDa.

H3(1-14)K9ac-MESNa thioester: ARTKQTAR(Kac)STGGK-C=OS(MESNa)

H3(5-13)K9ac-amide: Ac-QTAR(Kac)STGG-NH_2_

polyHis peptide: HHHHHHGGG-NH_2_

#### Octamer and Mononucleosome Preparation

H3WT and H3K9ac octamers were assembled by dialysis and purified by size exclusion chromatography (Superdex 200 increase column, Cytiva) as described previously.^50^ The mononucleosomes were prepared by mixing the respective octamer with ds601 DNA (sequence in **Supporting Information**) and dialyzing as described previously.^50^ The mononucleosomes were analyzed on a native 5% TBE gel to assess their quality (**Figure S6**).

#### Western Blotting

Reactions were loaded on a 4-12% bis tris gel (unless otherwise noted) and run with Bio-Rad MES buffer at 180V for about 45 min. Gels were transferred to Bio-Rad PVDF membrane with cold towbin buffer (25 mM Tris, 192 mM glycine, 10% (v/v) methanol, 0.06% (w/v) SDS) using a semi-dry transblot turbo set (Bio-Rad) for 30 min at 20 V. The membranes with total protein quantification were subjected to Revert Total Protein Stain 700 (Licor) according to the manufacturer’s protocol. All membranes were blocked with 3% nonfat milk (Genesee) in TBST (50 mM Tris pH 7.5, 150 mM NaCl, 0.05% Tween-20) at RT for 1 h. Anti-ADPr blots were performed using a 1:1000 antibody dilution in TBST of the Cell Signaling Technologies pan/mono antibody (#83732) overnight at 4°C. The blot in **Figure S1** was incubated with a 1:1000 dilution in TBST of the Anti-ADPr Millipore binding reagent overnight at 4°C. Anti-H3K9ac and anti-H3 (Cell Signaling Technologies) were used at a 1:2000 dilution. Anti-SIRT6 and anti-His tag (Cell Signaling Technologies) were used at a 1:1000 dilution. The blots were imaged on a Licor Odyssey system.

The specificity of the ADP-ribose antibodies was shown by using no cofactor (i.e., NAD^+^) and/or no enzyme lanes in the assays and by agreement between the two different antibodies (i.e., **Figure 1D and S1**). The H3K9ac antibody was similarly validated by using a “no SIRT6” condition along with the other conditions tested (**Figure S2A**) on semisynthetically-generated H3K9ac nucleosomes. The SIRT6 and H3 antibodies were assessed by the band aligning with the correct molecular weight for the respective recombinant protein as observed in a total protein stain. The His tag antibody was assessed by blotting for the SIRT6 before and after removal of the 6xHis tag by proteolytic cleavage.

#### Ribosylation Assays

The reactions (15 μL total volume) were set up and incubated as described in the respective figure legends. NAD^+^ was added last to initiate the reaction. A 5x reaction buffer was diluted to 1x for the assay. The 1x reaction buffer was 50 mM Tris pH 7.5, 20 mM NaCl, 2 mM MgCl_2_, 2 mM DTT and 1 mM NAD^+^ (NEB) was used when not otherwise specified in figure legends. The reactions were quenched by addition of 6x SDS loading buffer and boiled for 5 min.

Note: Reactions with SIRT6 and PARP1 (2-655) contained 10 μm SIRT6 and 2 μm PARP1 (2-655) to observe SIRT6 modification of PARP1 (2-655).

Note:SIRT6 auto mARylation reactions were set up (2 μm SIRT6) with no other substrate.

#### Electrophoretic Mobility Shift Assays

The SIRT6-WT or SIRT6-ΔC were incubated with 50 nM of ds601 DNA in reaction buffer (50 mM Tris pH 7.5, 20 mM NaCl, 2 mM MgCl_2_, 2 mM DTT) at 10 μL total volume. The reactions were incubated at RT for 15 min followed by the addition of 6X purple DNA loading dye (NEB). The reactions were analyzed on a 5% TBE gel with cold 1x TBE buffer (89 mM Tris, 89 mM boric acid, mM EDTA) for 100 min at 120 V, stained with ethidium bromide, and imaged using a Bio-Rad ChemiDoc system.

#### HPLC Deacetylation Assays

The reactions (50 μL total volume) included 10 μm 6xHis-SIRT6, 600 μm H3K9ac peptide, 1 mM NAD^+^ in reaction buffer (50 mM Tris pH 7.5, 20 mM NaCl, 2 mM MgCl_2_, 2 mM DTT) and were incubated at 30°C for 1 h. The reactions were quenched by the addition of 50 μL solvent A (water, 0.1% TFA). The samples were analyzed on a C18 column (Waters #186003624) using a gradient of 0-20% solvent B over 20 min (solvent B = 90% acetonitrile, 10% water, 0.1% TFA). After 20 min, the column washed with 100% solvent B for 5 min, and then re-equilibrated in 100% solvent A for 5 min. An Agilent 1260 Infinity system was used.

#### H3K9ac Mononucleosome Deacetylation Assay

The reactions (15 μL total volume) included 2 μm 6xHis-SIRT6 and 150 nM H3K9ac mononucleosome in reaction buffer (50 mM Tris pH 7.5, 20 mM NaCl, 2 mM MgCl_2_, 2 mM DTT, 1 mM NAD^+^) and were incubated at 30°C for 10 min. The reactions were quenched with 6x SDS loading buffer and boiled for 5 min.

#### Thrombin Cleavage Assay

The SIRT6 auto-ADP-ribosylation assay was set up as before with incubation for 20 min at 37 °C. To the reactions was added 1 U of thrombin (Cytiva) reconstituted in 1x PBS at 1 U/μL. The reactions were incubated for an additional 20 min at 37 °C and then quenched with 6X SDS loading buffer.

#### ARH3 Assay

MeCP2 mARylation assay was set up as previously described. Reactions were quenched with 1 mM SIRT6 inhibitor (Med Chem Express, OSS_128167) after 2 h and additionally incubated for either 30 min or 2 h more at 30 °C with 1 μM ARH3. We note that the SIRT6 inhibitor was dissolved to 100 mM in 100% DMSO and a working stock of 10 mM in 90% water and 10% DMSO was diluted from that. This 10 mM stock was cloudy and was homogenized thoroughly before adding to the reactions. **Figure S3E** shows the inhibition of SIRT6 by OSS_128167 (e.g., lane 2 compared to lane 3).

#### Mass Spectrometric Analysis of ADP-ribosylated MeCP2

The mass spectrometric analysis was carried out by Creative Proteomics as described in **Data File S3**. The full results are tabulated in **Data File S2**.

#### Software and Data Analysis

Prism GraphPad was used for preparing the plots in **Figure 1B, S2B, S5**, and **S6**. The unpaired, two-tailed t-test in **Figure 1B** was performed in Prism Graphpad. The exact p values are shown in the figure legends. The densitometric analysis was performed in Licor Image Studio with normalization to the SIRT6 band in the total protein stain. The sequence alignments were performed using NCBI Protein BLAST by searching in the UniprotKB/Swiss-prot database. The uncropped blots and gels, including all experimental replicates are in **Data File S4**. Each experiment was performed a minimum of 2 independent times, and the exact number of replicates is noted in each figure legend.

## Author Contributions

NP and KLD conceived the paper and designed all experiments. NP carried out the cloning, protein purification, and all biochemical assays. KLD prepared the peptides, histones, and histone octamers. NP and KLD wrote the manuscript.

## Acknowledgements

The authors acknowledge the NIH (R35GM143080) for funding. The content is solely the responsibility of the authors and does not necessarily represent the official views of the National Institutes of Health. We also acknowledge the DNA/Peptide Synthesis Core at the University of Utah.

## Data Availability

All the data from this study are contained within this article.

## Supporting Information

This article contains supporting information.

