## Supporting Information for "DNA stimulates SIRT6 to mono-ADP-ribosylate proteins within histidine repeats"

**Contents:**

Supporting Figures S1-S6
Annotated amino acid sequences for all proteins
Sequences for the DNA used in the enzyme assays

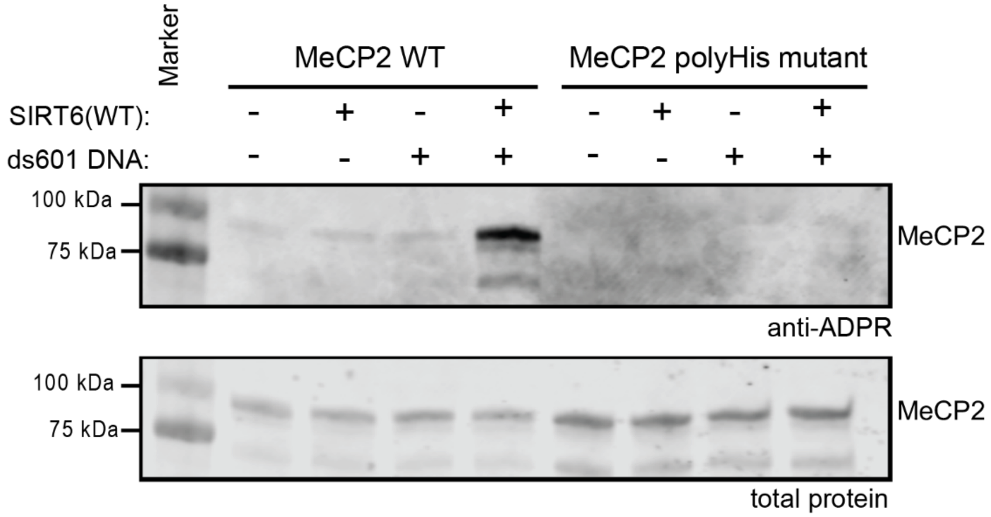


**Figure S1. Millipore pan/mono-ADPr binding reagent also detects polyHis-dependent mARylation of MeCP2 by SIRT6.** Immunoblot analysis of MeCP2-WT or -polyHis mutant (2 μM) mARylation by SIRT6 WT (2 μM) in the presence of ds601 DNA (1 μM) at 37°C for 2 h Western Blot developed with 1:1000 dilution of the Millipore pan/mono ADPR binding reagent (MABE1016), n = 2.

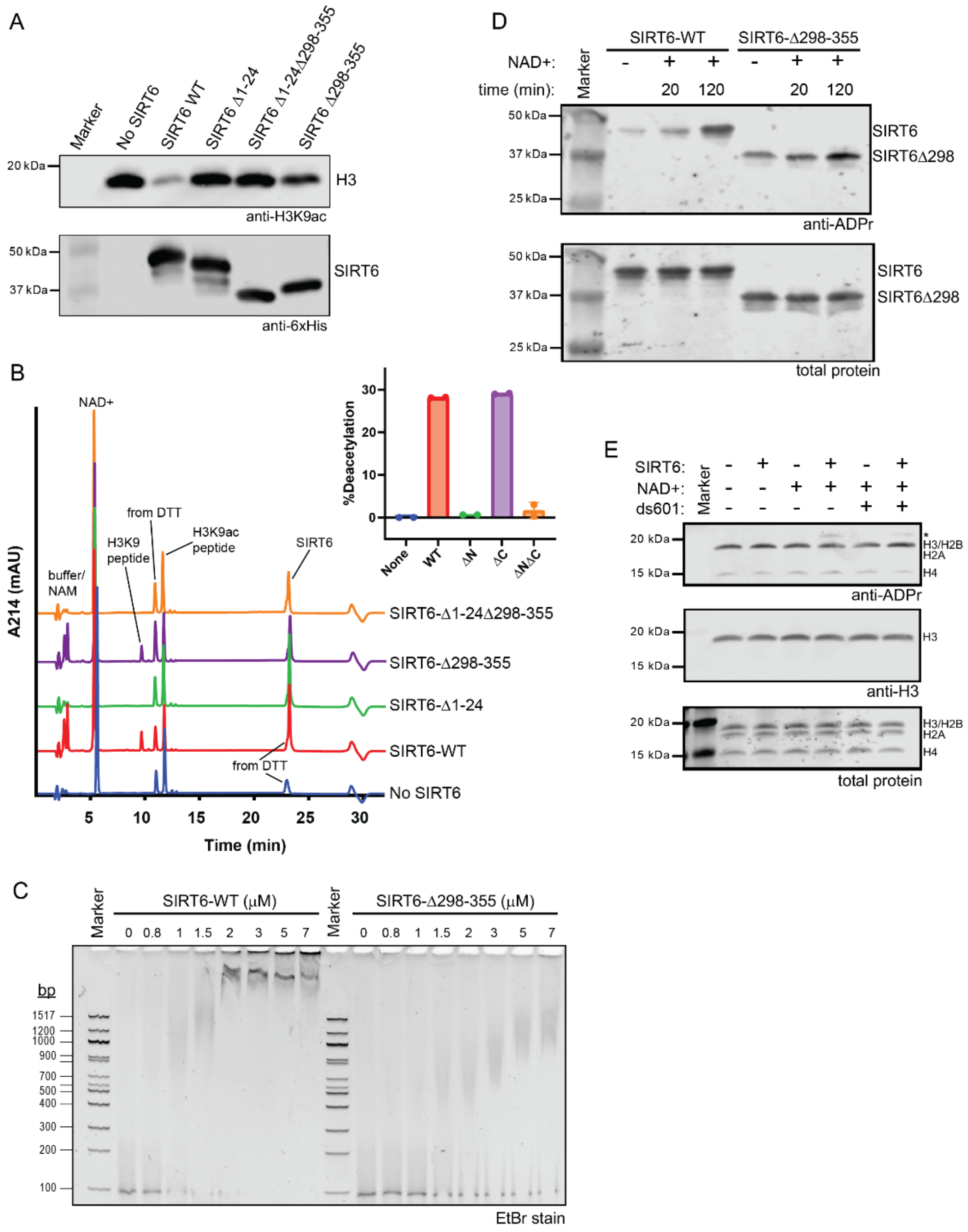


**Figure S2. SIRT6 binds to nucleosomes and to dsDNA.** A) Immunoblot analysis of H3K9ac nucleosome (150 nM) deacetylation assays with 6xHis-SIRT6 mutants (2 μM) at 30°C for 20 min, n = 2. B) Chromatograms of H3K9ac peptide (600 μM) assays with 6xHis-SIRT6 (10 μM),1 mM $NAD^+$, at 37°C for 1 h, n = 2, λ = 214 nm. The inset shows the extent of deacetylation that was calculated by quantifying the area under the deacetylated peptide peak and normalizing to the SIRT6 peak. The error bars represent ± S.D. C) Electrophoretic mobility shift assay (EMSA) of ds601 (100 nM) with Cleaved-SIRT6-WT or -ΔC at the indicated concentrations (in μM). The TBE gel was stained with EtBr, n = 3. D) Immunoblot analysis of autoMARylation by SIRT6-WT or -ΔC (2 μM) and 1 mM $NAD^+$, n = 2. E) Immunoblot analysis of WT nucleosomes (200 nM) incubated with SIRT6-WT (2 μM), ds601 (1 μM), and 1 mM $NAD^+$ at 37°C for 2 h, n = 2. The starred band in the ADPr blot is SIRT6-dependent, but it does not appear in the H3 blot.

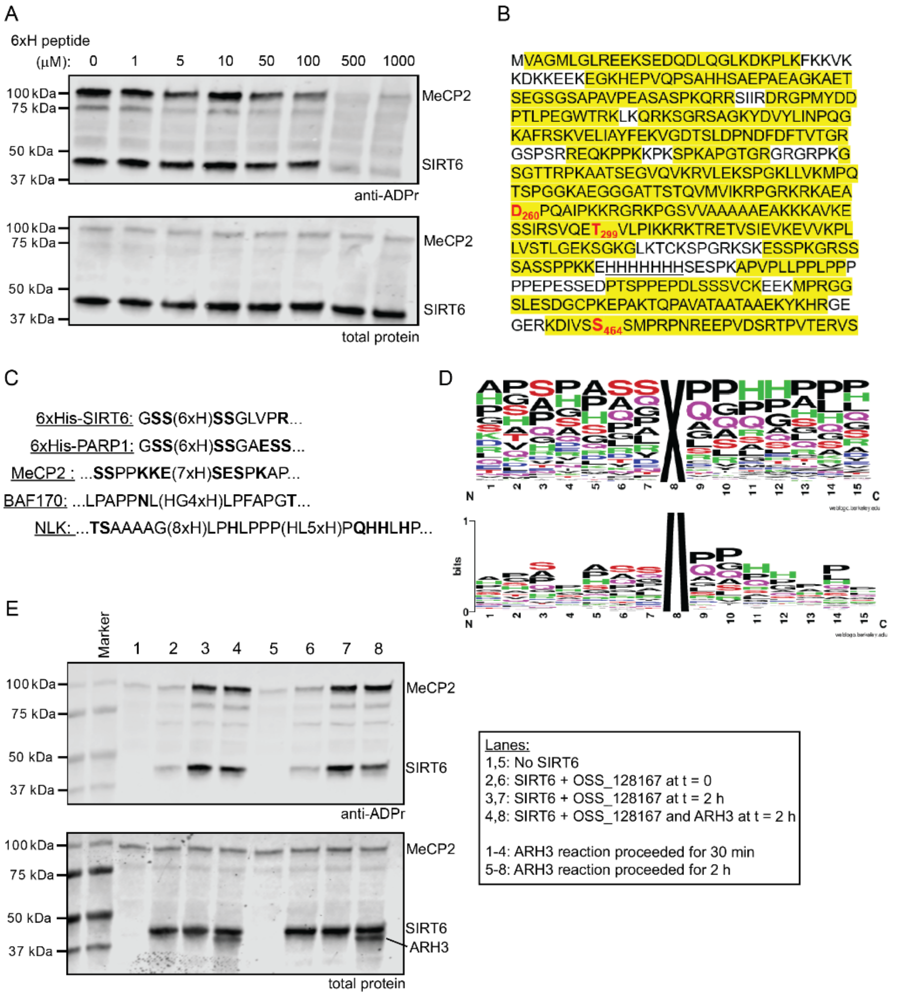


**Figure S3. SIRT6 mARylates histidine.** A) Immunoblot analysis of MeCP2 (2 μM) and SIRT6 (2 μM) mARylation reactions (1 μM ds601, 1 mM NAD$^+$, 37°C for 2 h) with the 6xH peptide (HHHHHHGGG), n = 2. B) Sequence of MeCP2. The yellow-highlighted portions were observed in the analysis by the identification of at least one peptide containing those amino acids (85% sequence coverage for this 486-amino acid protein), while the non-highlighted amino acids were not observed. The amino acids in red were found to be mARylated. The polyHis tract is underlined. All peptides that were identified are listed in **Data File S2**. C) Sequences surrounding the polyHis tract in the substrates tested in this study. Any residues that could be mARylation acceptors are bolded. D) Frequency map (top) and sequence map (bottom) for the surrounding residues of the polyHis tracts in all 129 of human polyHis proteins (listed in **Data File S1**). Position 8 ("X") represents the polyHis tract, which was defined as at least four sequential histidine residues plus histidines that were separated from the sequential histidine tract by no more than one intervening amino acid (e.g., HGHHHH or HHHHHSHRHH). E) Immunoblot analysis of MeCP2 (2 μM) and SIRT6 (2 μM) mARylation reactions (1 μM ds601, 1 mM NAD$^+$, 37°C for 2 h). After the 2 h mARylation reaction, SIRT6 inhibitor (OSS_128167, 1 mM) was added to quench the SIRT6. At the same time, ARH3 (1 μM) was added to reactions 7 and 8. Reactions 1-4 were then allowed to proceed for an additional 30 min. Reactions 5-8 were allowed to proceed for an additional 2 h.

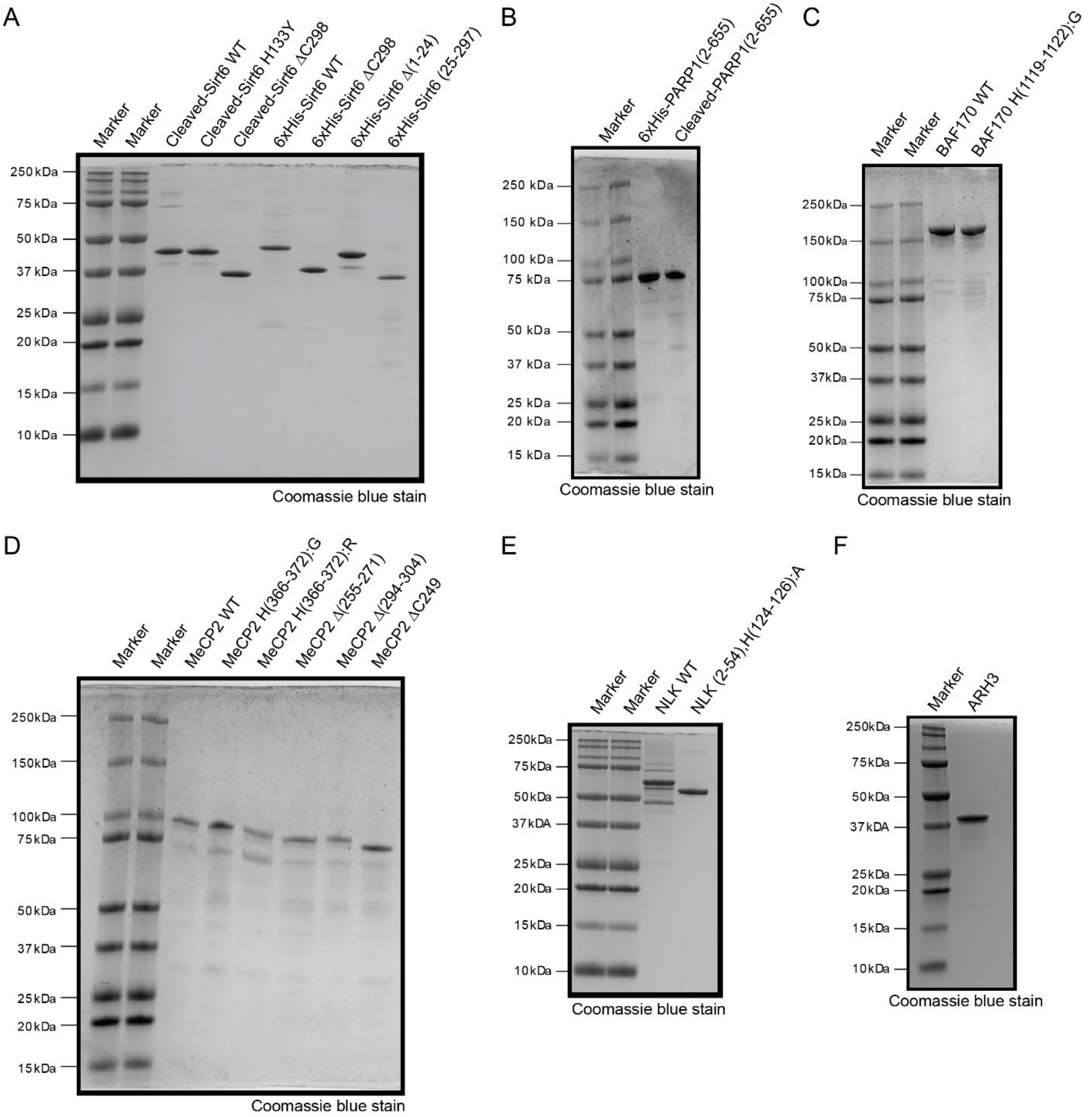


**Figure S4. Protein gels of proteins used in this study.** A) SIRT6, B) PARP1, C) BAF170/SMARCC2, D) MeCP2, E) NLK, F) ARH3.

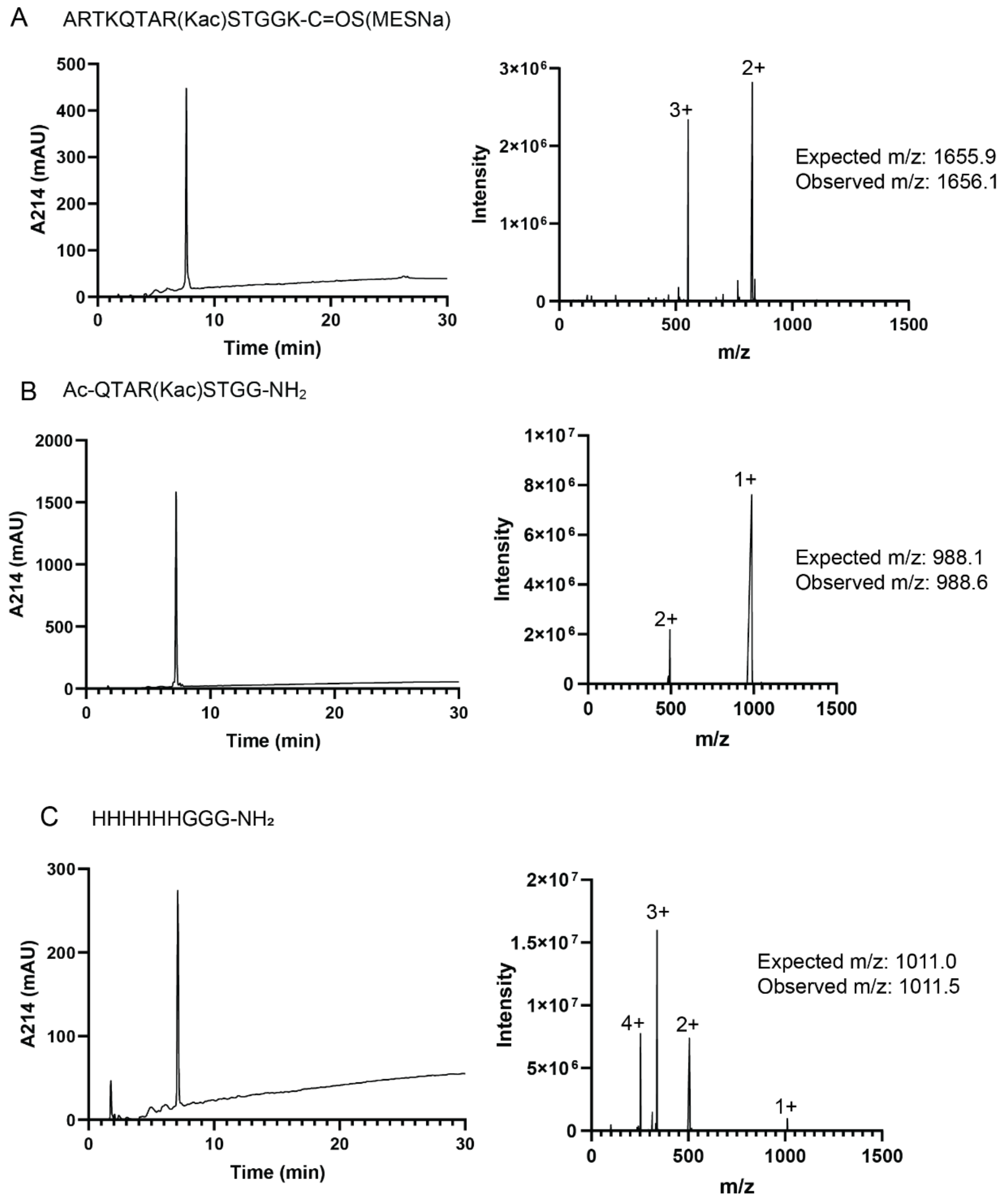


**Figure S5. Chromatographic and mass spectrometric analysis of the peptides used in this study.** A) H3(1-14)K9ac-MESNa thioester: ARTKQTAR(Kac)STGGK-COS(MESNa), B) H3(5-13)K9ac-amide: Ac-QTAR(Kac)STGG-$NH_2$, C) polyHis peptide: HHHHHHGGG-$NH_2$. The chromatograms (left) were collected using a 0-70% solvent B gradient over 30 min (solvent A = 100% water, 0.1% TFA and solvent B = 90% water, 10% water, 0.1% TFA) on a Waters BEH XBridge column (186003624) with detection of absorbance of 214 nm. The peak at 2 min is the void volume. The extracted ion chromatograms (right) were collected using a Waters QDa.

A

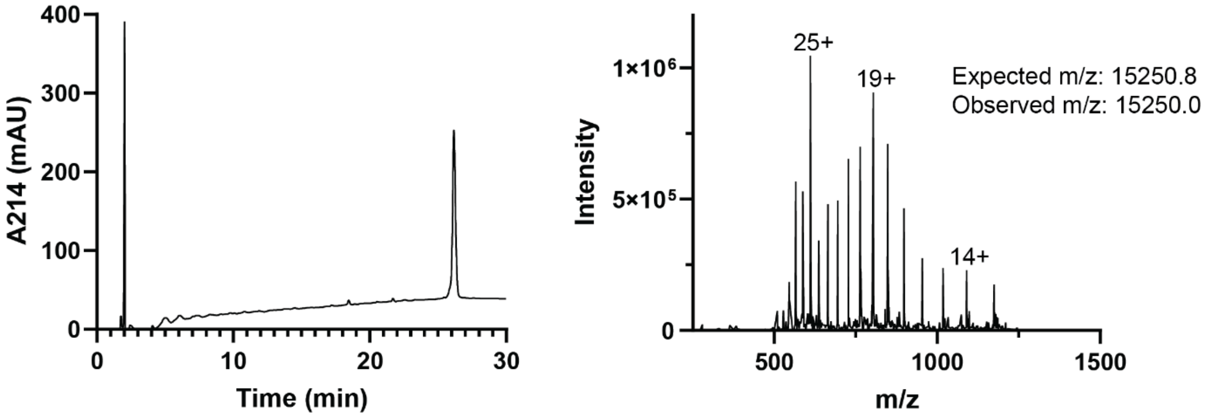


B

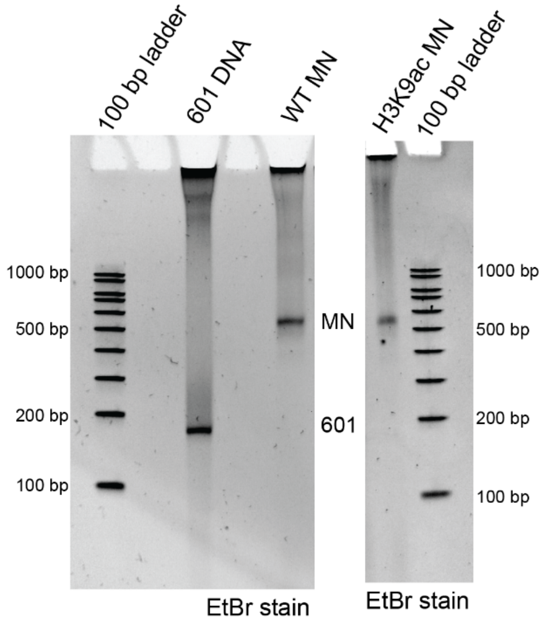


**Figure S6. Analysis of the H3K9ac histone and WT and H3K9ac mononucleosomes.** A) UV and extracted ion chromatograms for the H3K9ac histone. This sample was analyzed as described in the Figure S5 legend. B) Native gel analysis of the WT (left) and H3K9ac (right) mononucleosomes ("MN") and the 601 DNA (left) (5% TBE gel, EtBr staining). Note that Apex 100 bp ladder (Genesee #19-109) was used for these gels.

**Protein Sequences:**

**PARP1:**

**6XHis-PARP1 (2-655): Highlight = 6XHis Tag, in pet28a plasmid**

MGSSHHHHHHSSGAESSDKLYRVEYAKSGRASCKKCSESIPKDSLRMAIMVQSPMFDGKVPH
WYHFSCFWKVGHSIRHPDVEVDGFSELRWDDQQKVKKTAEAGGVTGKGQDGIGSKAEKTLG
DFAAEYAKSNRSTCKGCMEKIEKGQVRLSKKMVDPEKPQLGMIDRWYHPGCFVKNREELGFR
PEYSASQLKGFSLLATEDKEALKKQLPGVKSEGKRKGDEVDGVDEVAKKKSKKEKDKDSKLEK
ALKAQNDLIWNIKDELKKVCSTNDLKELLIFNKQQVPSGESAILDRVADGMVFGALLPCEECSG
QLVFKSDAYYCTGDVTAWTKCMVKTQTPNRKEWVTPKEFREISYLKKLKVKKQDRIFPPETSA
SVAATPPPSTASAPAAVNSSASADKPLSNMKILTLGKLSRNKDEVKAMIEKLGGKLTGTANKAS
LCISTKKEVEKMNKKMEEVKEANIRVVSEDFLQDVSASTKSLQELFLAHILSPWGAEVKAEPVE
VVAPRGKSGAALSKKSKGQVKEEGINKSEKRMKLTLKGGAAVDPDSGLEHSAHVLEKGGKVF
SATLGLVDIVKGTNSYYKLQLLEDDKENRYWIFRSWGRVGTVIGSNKLEQMPSKEDAIEHFMKL
YEEKTGNAWHSKNFTKYPKKFYPLEIDYGQDEEAVKKL

**Cleaved PARP1 (2-655): in pet28a plasmid**

AESSDKLYRVEYAKSGRASCKKCSESIPKDSLRMAIMVQSPMFDGKVPHWYHFSCFWKVGHS
IRHPDVEVDGFSELRWDDQQKVKKTAEAGGVTGKGQDGIGSKAEKTLGDFAAEYAKSNRSTC
KGCMEKIEKGQVRLSKKMVDPEKPQLGMIDRWYHPGCFVKNREELGFRPEYSASQLKGFSLL
ATEDKEALKKQLPGVKSEGKRKGDEVDGVDEVAKKKSKKEKDKDSKLEKALKAQNDLIWNIKD
ELKKVCSTNDLKELLIFNKQQVPSGESAILDRVADGMVFGALLPCEECSGQLVFKSDAYYCTGD
VTAWTKCMVKTQTPNRKEWVTPKEFREISYLKKLKVKKQDRIFPPETSASVAATPPPSTASAPA
AVNSSASADKPLSNMKILTLGKLSRNKDEVKAMIEKLGGKLTGTANKASLCISTKKEVEKMNKK
MEEVKEANIRVVSEDFLQDVSASTKSLQELFLAHILSPWGAEVKAEPVEVVAPRGKSGAALSKK
SKGQVKEEGINKSEKRMKLTLKGGAAVDPDSGLEHSAHVLEKGGKVFSATLGLVDIVKGTNSY
YKLQLLEDDKENRYWIFRSWGRVGTVIGSNKLEQMPSKEDAIEHFMKLYEEKTGNAWHSKNFT
KYPKKFYPLEIDYGQDEEAVKKL

**MeCP2:**

**MeCP2 WT (2-486): Highlight = GST tag, in pet28a plasmid**

MSPILGYWKIKGLVQPTRLLLEYLEEKYEEHLYERDEGDKWRNKKFELGLEFPNLPYYIDGDVK
LTQSMAIIRYIADKHNMLGGCPKERAEISMLEGAVLDIRYGVSRIAYSKDFETLKVDFLSKLPEML
KMFEDRLCHKTYLNGDHVTHPDFMLYDALDVVLYMDPMCLDAFPKLVCFKKRIEAIPQIDKYLK
SSKYIAWPLQGWQATFGGGDHPPKLVPRGSVAGMLGLREEKSEDQDLQGLKDKPLKFKKVKK
DKKEEKEGKHEPVQPSAHHSAEPAEAGKAETSEGSGSAPAVPEASASPKQRRSIIRDRGPMY
DDPTLPEGWTRKLKQRKSGRSAGKYDVYLINPQGKAFRSKVELIAYFEKVGDTSLDPNDFDFT
VTGRGSPSRREQKPPKKPKSPKAPGTGRGRGRPKGSGTTRPKAATSEGVQVKRVLEKSPGK
LLVKMPFQTSPGGKAEGGGATTSTQVMVIKRPGRKRKAEADPQAIPKKRGRKPGSVVAAAAA
EAKKKAVKESSIRSVQETVLPIKKRKTRETVSIEVKEVVKPLLVSTLGEKSGKGLKTCKSPGRKS
KESSPKGRSSSASSPPKKEHHHHHHHSESPKAPVPLLPPLPPPPPEPESSEDPTSPPEPQDLS
SSVCKEEKMPRGGSLESDGCPKEPAKTQPAVATAATAAEKYKHRGEGERKDIVSSSMPRPNR
EEPVDSRTPVTERVS

**MeCP2 H (366-372) toG: Underline = HtoG mutation, Highlight= GST tag, in pet28a plasmid**

MSPILGYWKIKGLVQPTRLLLEYLEEKYEEHLYERDEGDKWRNKKFELGLEFPNLPYYIDGDVK
LTQSMAIIRYIADKHNMLGGCPKERAEISMLEGAVLDIRYGVSRIAYSKDFETLKVDFLSKLPEML
KMFEDRLCHKTYLNGDHVTHPDFMLYDALDVVLYMDPMCLDAFPKLVCFKKRIEAIPQIDKYLK
SSKYIAWPLQGWQATFGGGDHPPKLVPRGSVAGMLGLREEKSEDQDLQGLKDKPLKFKKVKK
DKKEEKEGKHEPVQPSAHHSAEPAEAGKAETSEGSGSAPAVPEASASPKQRRSIIRDRGPMY
DDPTLPEGWTRKLKQRKSGRSAGKYDVYLINPQGKAFRSKVELIAYFEKVGDTSLDPNDFDFT
VTGRGSPSRREQKPPKKPKSPKAPGTGRGRGRPKGSGTTRPKAATSEGVQVKRVLEKSPGK
LLVKMPFQTSPGGKAEGGGATTSTQVMVIKRPGRKRKAEADPQAIPKKRGRKPGSVVAAAAA
EAKKKAVKESSIRSVQETVLPIKKRKTRETVSIEVKEVVKPLLVSTLGEKSGKGLKTCKSPGRKS
KESSPKGRSSSASSPPKKEGGGGGGGSESPKAPVPLLPPLPPPPPEPESSEDPTSPPEPQDL
SSSVCKEEKMPRGGSLESDGCPKEPAKTQPAVATAATAAEKYKHRGEGERKDIVSSSMPRPN
REEPVDSRTPVTERVS

**MeCP2 H(366-372)toR: Underline = HtoR mutation, Highlight= GST tag, in pet28a plasmid**

MSPILGYWKIKGLVQPTRLLLEYLEEKYEEHLYERDEGDKWRNKKFELGLEFPNLPYYIDGDVK
LTQSMAIIRYIADKHNMLGGCPKERAEISMLEGAVLDIRYGVSRIAYSKDFETLKVDFLSKLPEML
KMFEDRLCHKTYLNGDHVTHPDFMLYDALDVVLYMDPMCLDAFPKLVCFKKRIEAIPQIDKYLK
SSKYIAWPLQGWQATFGGGDHPPKLVPRGSVAGMLGLREEKSEDQDLQGLKDKPLKFKKVKK
DKKEEKEGKHEPVQPSAHHSAEPAEAGKAETSEGSGSAPAVPEASASPKQRRSIIRDRGPMY
DDPTLPEGWTRKLKQRKSGRSAGKYDVYLINPQGKAFRSKVELIAYFEKVGDTSLDPNDFDFT
VTGRGSPSRREQKPPKKPKSPKAPGTGRGRGRPKGSGTTRPKAATSEGVQVKRVLEKSPGK
LLVKMPFQTSPGGKAEGGGATTSTQVMVIKRPGRKRKAEADPQAIPKKRGRKPGSVVAAAAA
EAKKKAVKESSIRSVQETVLPIKKRKTRETVSIEVKEVVKPLLVSTLGEKSGKGLKTCKSPGRKS
KESSPKGRSSSASSPPKKERRRRRRRSESPKAPVPLLPPLPPPPPEPESSEDPTSPPEPQDLS
SSVCKEEKMPRGGSLESDGCPKEPAKTQPAVATAATAAEKYKHRGEGERKDIVSSSMPRPNR
EEPVDSRTPVTERVS

**MeCP2 Δ255-271: Highlight= GST tag, in pet28a plasmid**

MSPILGYWKIKGLVQPTRLLLEYLEEKYEEHLYERDEGDKWRNKKFELGLEFPNLPYYIDGDVK
LTQSMAIIRYIADKHNMLGGCPKERAEISMLEGAVLDIRYGVSRIAYSKDFETLKVDFLSKLPEML
KMFEDRLCHKTYLNGDHVTHPDFMLYDALDVVLYMDPMCLDAFPKLVCFKKRIEAIPQIDKYLK
SSKYIAWPLQGWQATFGGGDHPPKLVPRGSVAGMLGLREEKSEDQDLQGLKDKPLKFKKVKK
DKKEEKEGKHEPVQPSAHHSAEPAEAGKAETSEGSGSAPAVPEASASPKQRRSIIRDRGPMY
DDPTLPEGWTRKLKQRKSGRSAGKYDVYLINPQGKAFRSKVELIAYFEKVGDTSLDPNDFDFT
VTGRGSPSRREQKPPKKPKSPKAPGTGRGRGRPKGSGTTRPKAATSEGVQVKRVLEKSPGK
LLVKMPFQTSPGGKAEGGGATTSTQVMVIKRPGRKPGSVVAAAAAEAKKKAVKESSIRSVQET
VLPIKKRKTRETVSIEVKEVVKPLLVSTLGEKSGKGLKTCKSPGRKSKESSPKGRSSSASSPPK
KEHHHHHHHSESPKAPVPLLPPLPPPPPEPESSEDPTSPPEPQDLSSSVCKEEKMPRGGSLES
DGCPKEPAKTQPAVATAATAAEKYKHRGEGERKDIVSSSMPRPNREEPVDSRTPVTERVS

**MeCP2 Δ294-304: Highlight= GST tag, in pet28a plasmid**

MSPILGYWKIKGLVQPTRLLLEYLEEKYEEHLYERDEGDKWRNKKFELGLEFPNLPYYIDGDVK
LTQSMAIIRYIADKHNMLGGCPKERAEISMLEGAVLDIRYGVSRIAYSKDFETLKVDFLSKLPEML
KMFEDRLCHKTYLNGDHVTHPDFMLYDALDVVLYMDPMCLDAFPKLVCFKKRIEAIPQIDKYLK
SSKYIAWPLQGWQATFGGGDHPPKLVPRGSVAGMLGLREEKSEDQDLQGLKDKPLKFKKVKK
DKKEEKEGKHEPVQPSAHHSAEPAEAGKAETSEGSGSAPAVPEASASPKQRRSIIRDRGPMY
DDPTLPEGWTRKLKQRKSGRSAGKYDVYLINPQGKAFRSKVELIAYFEKVGDTSLDPNDFDFT
VTGRGSPSRREQKPPKKPKSPKAPGTGRGRGRPKGSGTTRPKAATSEGVQVKRVLEKSPGK
LLVKMPFQTSPGGKAEGGGATTSTQVMVIKRPGRKRKAEADPQAIPKKRGRKPGSVVAAAAA
EAKKKAVKESSIKRKTRETVSIEVKEVVKPLLVSTLGEKSGKGLKTCKSPGRKSKESSPKGRSS
SASSPPKKEHHHHHHHSESPKAPVPLLPPLPPPPPEPESSEDPTSPPEPQDLSSSVCKEEKMP
RGGSLESDGCPKEPAKTQPAVATAATAAEKYKHRGEGERKDIVSSSMPRPNREEPVDSRTPV
TERVS

**MeCP2 ΔdelC459 (2-458): Highlight= GST tag, in pet28a plasmid**

MSPILGYWKIKGLVQPTRLLLEYLEEKYEEHLYERDEGDKWRNKKFELGLEFPNLPYYIDGDVK
LTQSMAIIRYIADKHNMLGGCPKERAEISMLEGAVLDIRYGVSRIAYSKDFETLKVDFLSKLPEML
KMFEDRLCHKTYLNGDHVTHPDFMLYDALDVVLYMDPMCLDAFPKLVCFKKRIEAIPQIDKYLK
SSKYIAWPLQGWQATFGGGDHPPKLVPRGSVAGMLGLREEKSEDQDLQGLKDKPLKFKKVKK
DKKEEKEGKHEPVQPSAHHSAEPAEAGKAETSEGSGSAPAVPEASASPKQRRSIIRDRGPMY
DDPTLPEGWTRKLKQRKSGRSAGKYDVYLINPQGKAFRSKVELIAYFEKVGDTSLDPNDFDFT
VTGRGSPSRREQKPPKKPKSPKAPGTGRGRGRPKGSGTTRPKAATSEGVQVKRVLEKSPGK
LLVKMPFQTSPGGKAEGGGATTSTQVMVIKRPGRKRKAEADPQAIPKKRGRKPGSVVAAAAA
EAKKKAVKESSIRSVQETVLPIKKRKTRETVSIEVKEVVKPLLVSTLGEKSGKGLKTCKSPGRKS
KESSPKGRSSSASSPPKKEHHHHHHHSESPKAPVPLLPPLPPPPPEPESSEDPTSPPEPQDLS
SSVCKEEKMPRGGSLESDGCPKEPAKTQPAVATAATAAEKYKHRGEGER

**SIRT6:**

**6XHis-Thrombin-SIRT6 WT: Highlight = 6XHis and thrombin site, in pet28a plasmid**

MGSSHHHHHHSSGLVPRGSVNYAAGLSPYADKGKCGLPEIFDPPEELERKVWELARLVWQSS
SVVFHTGAGISTASGIPDFRGPHGVWTMEERGLAPKFDTTFESARPTQTHMALVQLERVGLLR
FLVSQNVDGLHVRSGFPRDKLAELHGNMFVEECAKCKTQYVRDTVVGTMGLKATGRLCTVAK
ARGLRACRGELRDTILDWEDSLPDRDLALADEASRNADLSITLGTSLQIRPSGNLPLATKRRGG
RLVIVNLQPTKHDRHADLRIHGYVDEVMTRLMKHLGLEIPAWDGPRVLERALPPLPRPPTPKLE
PKEESPTRINGSIPAGPKQEPCAQHNGSEPASPKRERPTSPAPHRPPKRVKAKAVPS

**6XHis-Thrombin-SIRT6 ΔN24 (25-355): Highlight = 6XHis and thrombin site, in pet28a plasmid**

MGSSHHHHHHSSGLVPRGDPPEELERKVWELARLVWQSSSVVFHTGAGISTASGIPDFRGPH
GVWTMEERGLAPKFDTTFESARPTQTHMALVQLERVGLLRFLVSQNVDGLHVRSGFPRDKLA
ELHGNMFVEECAKCKTQYVRDTVVGTMGLKATGRLCTVAKARGLRACRGELRDTILDWEDSL
PDRDLALADEASRNADLSITLGTSLQIRPSGNLPLATKRRGGRLVIVNLQPTKHDRHADLRIHGY
VDEVMTRLMKHLGLEIPAWDGPRVLERALPPLPRPPTPKLEPKEESPTRINGSIPAGPKQEPCA
QHNGSEPASPKRERPTSPAPHRPPKRVKAKAVPS

**6XHis-Thrombin-SIRT6 ΔC298 (2-297): Highlight = 6XHis and thrombin site, in pet28a plasmid**

MGSSHHHHHHSSGLVPRGSVNYAAGLSPYADKGKCGLPEIFDPPEELERKVWELARLVWQSS
SVVFHTGAGISTASGIPDFRGPHGVWTMEERGLAPKFDTTFESARPTQTHMALVQLERVGLLR
FLVSQNVDGLHVRSGFPRDKLAELHGNMFVEECAKCKTQYVRDTVVGTMGLKATGRLCTVAK
ARGLRACRGELRDTILDWEDSLPDRDLALADEASRNADLSITLGTSLQIRPSGNLPLATKRRGG
RLVIVNLQPTKHDRHADLRIHGYVDEVMTRLMKHLGLEIPAWDGPRVLERALPPLPRPPTPKLE

**6XHis-Thrombin-SIRT6 ΔN24ΔC298 (25-297): Highlight = 6XHis and thrombin site, in pet28a plasmid**

MGSSHHHHHHSSGLVPRGDPPEELERKVWELARLVWQSSSVVFHTGAGISTASGIPDFRGPH
GVWTMEERGLAPKFDTTFESARPTQTHMALVQLERVGLLRFLVSQNVDGLHVRSGFPRDKLA
ELHGNMFVEECAKCKTQYVRDTVVGTMGLKATGRLCTVAKARGLRACRGELRDTILDWEDSL
PDRDLALADEASRNADLSITLGTSLQIRPSGNLPLATKRRGGRLVIVNLQPTKHDRHADLRIHGY
VDEVMTRLMKHLGLEIPAWDGPRVLERALPPLPRPPTPKLE

**SIRT6 WT (2-355):, in pet28a plasmid**

SVNYAAGLSPYADKGKCGLPEIFDPPEELERKVWELARLVWQSSSVVFHTGAGISTASGIPDFR
GPHGVWTMEERGLAPKFDTTFESARPTQTHMALVQLERVGLLRFLVSQNVDGLHVRSGFPRD
KLAELHGNMFVEECAKCKTQYVRDTVVGTMGLKATGRLCTVAKARGLRACRGELRDTILDWE
DSLPDRDLALADEASRNADLSITLGTSLQIRPSGNLPLATKRRGGRLVIVNLQPTKHDRHADLRI
HGYVDEVMTRLMKHLGLEIPAWDGPRVLERALPPLPRPPTPKLEPKEESPTRINGSIPAGPKQE
PCAQHNGSEPASPKRERPTSPAPHRPPKRVKAKAVPS

**SIRT6 H133Y (2-355): Underline= H133 to Y mutation, in pet28a plasmid**

SVNYAAGLSPYADKGKCGLPEIFDPPEELERKVWELARLVWQSSSVVFHTGAGISTASGIPDFR
GPHGVWTMEERGLAPKFDTTFESARPTQTHMALVQLERVGLLRFLVSQNVDGLHVRSGFPRD
KLAELYGNMFVEECAKCKTQYVRDTVVGTMGLKATGRLCTVAKARGLRACRGELRDTILDWE
DSLPDRDLALADEASRNADLSITLGTSLQIRPSGNLPLATKRRGGRLVIVNLQPTKHDRHADLRI
HGYVDEVMTRLMKHLGLEIPAWDGPRVLERALPPLPRPPTPKLEPKEESPTRINGSIPAGPKQE
PCAQHNGSEPASPKRERPTSPAPHRPPKRVKAKAVPS

**SIRT6 ΔC298 (2-297): in pet28a plasmid**

SVNYAAGLSPYADKGKCGLPEIFDPPEELERKVWELARLVWQSSSVVFHTGAGISTASGIPDFR
GPHGVWTMEERGLAPKFDTTFESARPTQTHMALVQLERVGLLRFLVSQNVDGLHVRSGFPRD
KLAELHGNMFVEECAKCKTQYVRDTVVGTMGLKATGRLCTVAKARGLRACRGELRDTILDWE
DSLPDRDLALADEASRNADLSITLGTSLQIRPSGNLPLATKRRGGRLVIVNLQPTKHDRHADLRI
HGYVDEVMTRLMKHLGLEIPAWDGPRVLERALPPLPRPPTPKLE

**NLK WT: (1-527): Highlight = FLAG tag, in bacmid**

MSLCGARANAKMMAAYNGGTSAAAAGHHHHHHHHHLPHLPPPHLHHHHHPQHHLHPGSAAAV
HPVQQHTSSAAAAAAAAAAAAAMLNPGQQQPYFPSPAPGQAPGPAAAAPAQVQAAAAATVK
AHHHQHSHHPQQQLDIEPDRPIGYGAFGVVWSVTDPRDGKRVALKKMPNVFQNLVSCKRVFR

ELKMLCFFKHDNVLSALDILQPPHIDYFEEIYVVTELMQSDLHKIIVSPQPLSSDHVKVFLYQILRG
LKYLHSAGILHRDIKPGNLLVNSNCVLKICDFGLARVEELDESRHMTQEVVTQYYRAPEILMGSR
HYSNAIDIWSVGCIFAELLGRRILFQAQSPIQQLDLITDLLGTPSLEAMRTACEGAKAHILRGPHK
QPSLPVLYTLSSQATHEAVHLLCRMLVFDPSKRISAKDALAHPYLDEGRLRYHTCMCKCCFSTS
TGRVYTSDFEPVTNPKFDDTFEKNLSSVRQVKEIIHQFILEQQKGNRVPLCINPQSAAFKSFISS
TVAQPSEMPPSPLVWEGGSGGDYKDDDDK

**NLK1 Δ(2-54) H (78-80)to A: Underline = HtoA Highlight = FLAG tag in bacmid**

MPGSAAAVHPVQQHTSSAAAAAAAAAAAAAMLNPGQQQPYFPSPAPGQAPGPAAAAPAQVQ
AAAAATVKAAAAQHSHHPQQQLDIEPDRPIGYGAFGVVWSVTDPRDGKRVALKKMPNVFQNL
VSCKRVFRELKMLCFFKHDNVLSALDILQPPHIDYFEEIYVVTELMQSDLHKIIVSPQPLSSDHVK
VFLYQILRGLKYLHSAGILHRDIKPGNLLVNSNCVLKICDFGLARVEELDESRHMTQEVVTQYYR
APEILMGSRHYSNAIDIWSVGCIFAELLGRRILFQAQSPIQQLDLITDLLGTPSLEAMRTACEGAK
AHILRGPHKQPSLPVLYTLSSQATHEAVHLLCRMLVFDPSKRISAKDALAHPYLDEGRLRYHTC
MCKCCFSTSTGRVYTSDFEPVTNPKFDDTFEKNLSSVRQVKEIIHQFILEQQKGNRVPLCINPQ
SAAFKSFISSTVAQPSEMPPSPLVWEGGSGGDYKDDDDK

**BAF170 WT (1-1214): Highlight = 3XFLAG tag, in bacmid**

MAVRKKDGGPNVKYYEAADTVTQFDNVRLWLGKNYKKYIQAEPPTNKSLSSLVVQLLQFQ
EEVFGKHVSNAPLTKLPIKCFLDFKAGGSLCHILAAAYKFKSDQGWRRYDFQNPSRMDRN
VEMFMTIEKSLVQNNCLSRPNIFLCPEIEPKLLGKLKDIIKRHQGTVTEDKNNASHVVYP
VPGNLEEEEWVRPVMKRDKQVLLHWGYYPDSYDTWIPASEIEASVEDAPTPEKPRKVHAK
WILDTDTFNEWMNEEDYEVNDDKNPVSRRKKISAKTLTDEVNSPDSDRRDKKGGNYKKRK
RSPSPSPTPEAKKKNAKKGPSTPYTKSKRGHREEEQEDLTKDMDEPSPVPNVEEVTLPKT
VNTKKDSESAPVKGGTMTDLDEQEDESMETTGKDEDENSTGNKGEQTKNPDLHEDNVTEQ
THHIIIPSYAAWFDYNSVHAIERRALPEFFNGKNKSKTPEIYLAYRNFMIDTYRLNPQEY
LTSTACRRNLAGDVCAIMRVHAFLEQWGLINYQVDAESRPTPMGPPPTSHFHVLADTPSG
LVPLQPKTPQQTSASQQMLNFPDKGKEKPTDMQNFGLRTDMYTKKNVPSKSKAAASATRE
WTEQETLLLLEALEMYKDDWNKVSEHVGSRTQDECILHFLRLPIEDPYLEDSEASLGPLA
YQPIPFSQSGNPVMSTVAFLASVVDPRVASAAAKSALEEFSKMKEEVPTALVEAHVRKVE
EAAKVTGKADPAFGLESSGIAGTTSDEPERIEESGNDEARVEGQATDEKKEPKEPREGGG
AIEEEAKEKTSEAPKKDEEKGKEGDSEKESEKSDGDPIVDPEKEKEPKEGQEEVLKEVVE
SEGERKTKVERDIGEGNLSTAAAAALAAAAVKAKHLAAVEERKIKSLVALLVETQMKKLE
IKLRHFEELETIMDREREALEYQRQQLLADRQAFHMEQLKYAEMRARQQHFQQMHQQQQQ
PPPALPPGSQPIPPTGAAGPPAVHGLAVAPASVVPAPAGSGAPPGSLGPSEQIGQAGSTA
GPQQQQPAGAPQPGAVPPGVPPPGPHGPSPFPNQQTPPSMMPGAVPGSGHPGVAGNAPLG
LPFGMPPPPPPPAPSIIPFGSLADSISINLPAPPNLHGHHHHLPFAPGTLPPPNLPVSMA
NPLHPNLPATTTMPSSLPLGPGLGSAAAQSPAIVAAVQGNLLPSASPLPDPGTPLPPDPT
APSPGTVTPVPPPQEFGGDYKDDDDKGGSDYKDDDDKGGSDYKDDDDK

**BAF170 4XHtoG: Underline = 4XHtoG Highlight = 3XFLAG tag, in bacmid**

MAVRKKDGGPNVKYYEAADTVTQFDNVRLWLGKNYKKYIQAEPPTNKSLSSLVVQLLQFQ
EEVFGKHVSNAPLTKLPIKCFLDFKAGGSLCHILAAAYKFKSDQGWRRYDFQNPSRMDRN
VEMFMTIEKSLVQNNCLSRPNIFLCPEIEPKLLGKLKDIIKRHQGTVTEDKNNASHVVYP
VPGNLEEEEWVRPVMKRDKQVLLHWGYYPDSYDTWIPASEIEASVEDAPTPEKPRKVHAK
WILDTDTFNEWMNEEDYEVNDDKNPVSRRKKISAKTLTDEVNSPDSDRRDKKGGNYKKRK
RSPSPSPTPEAKKKNAKKGPSTPYTKSKRGHREEEQEDLTKDMDEPSPVPNVEEVTLPKT

VNTKKDSESAPVKGGTMTDLDEQEDESMETTGKDEDENSTGNKGEQTKNPDLHEDNVTEQ
THHIIIPSYAAWFDYNSVHAIERRALPEFFNGKNKSKTPEIYLAYRNFMIDTYRLNPQEY
LTSTACRRNLAGDVCAIMRVHAFLEQWGLINYQVDAESRPTPMGPPPTSHFHVLADTPSG
LVPLQPKTPQQTSASQQMLNFPDKGKEKPTDMQNFGLRTDMYTKKNVPSKSKAAASATRE
WTEQETLLLLEALEMYKDDWNKVSEHVGSRTQDECILHFLRLPIEDPYLEDSEASLGPLA
YQPIPFSQSGNPVMSTVAFLASVVDPRVASAAAKSALEEFSKMKEEVPTALVEAHVRKVE
EAAKVTGKADPAFGLESSGIAGTTSDEPERIEESGNDEARVEGQATDEKKEPKEPREGGG
AIEEEAKEKTSEAPKKDEEKGKEGDSEKESEKSDGDPIVDPEKEKEPKEGQEEVLKEVVE
SEGERKTKVERDIGEGNLSTAAAAALAAAAVKAKHLAAVEERKIKSLVALLVETQMKKLE
IKLRHFEELETIMDREREALEYQRQQLLADRQAFHMEQLKYAEMRARQQHFQQMHQQQQQ
PPPALPPGSQPIPPTGAAGPPAVHGLAVAPASVVPAPAGSGAPPGSLGPSEQIGQAGSTA
GPQQQQPAGAPQPGAVPPGVPPPGPHGPSPFPNQQTPPSMMPGAVPGSGHPGVAGNAPLG
LPFGMPPPPPPPAPSIIPFGSLADSISINLPAPPNLHGGGGGLPFAPGTLPPPNLPVSMA
NPLHPNLPATTTMPSSLPLGPGLGSAAAQSPAIVAAVQGNLLPSASPLPDPGTPLPPDPT
APSPGTVTPVPPPQEFGGDYKDDDDKGGSDYKDDDDKGGSDYKDDDDK

**ARH3 (14-363): in SUMO-pET30 Highlight = HA tag**

AGAARSLSRFRGCLAGALLGDCVGSFYEAHDTVDLTSVLRHVQSLEPDP
GTPGSERTEALYYTD DTAMARALVQSLLAKEAFDEVDMAHRFAQEYK
KDPDRGYGAGVVTVFKKLLNPKCRDVFEPARA QFNGKGSYGNGGAM
RVAGISLAYSSVQDVQKFARLSAQLTHASSLGYNGAILQALAVHLALQGES
SSEHFLKQLLGHMEDLEGDAQSVLDARELGMEERPYSSRLKKIGELLDQASV
TREEVVSELGNGIA AFESVPTAIYCFLRCMEPDPEIPSAFNSLQRTL
IYSISLGGDTDTIATMAGAIAGAYYGMDQVPES
WQQSCEGYEETDILAQSLHR VFQKSAQSLHRVFQKSYPYDVPDYARS

**DNA Sequences:**

ds601 DNA:

CTACTGGTACGGCAGACAGGATGTATATATCTGACACGTGCCTGGAGACTAGGGAGTAAT
CCCCTTGGCGGTTAAAACGCGGGGGACAGCGCGTACGTGCGTTTAAGCGGTGCTAGAGC
TGTCTACGACCAATTGAGCGGCCTCGGCACCGGGATTCTCCAGTATTCGAGGCCGTTC

Note: Used ss601 for **Figure 2B**.

Note: For the “plasmid” DNA used in **Figure 2C**, a pET28a plasmid with the TEV-SIRT6(WT) gene was used. The full sequence of that plasmid is below:

CCGCACCAACGCGCAGCCCGGACTCGGTAATGGCGCGCATTGCGCCCAGCGCCATCTGA
TCGTTGGCAACCAGCATCGCAGTGGGAACGATGCCCTCATTCAGCATTTGCATGGTTTGTT
GAAAACCGGACATGGCACTCCAGTCGCCTTCCCGTTCCGCTATCGGCTGAATTTGATTGC
GAGTGAGATATTTATGCCAGCCAGCCAGACGCAGACGCGCCGAGACAGAACTTAATGGGC
CCGCTAACAGCGCGATTTGCTGGTGACCCAATGCGACCAGATGCTCCACGCCCAGTCGCG
TACCGTCTTCATGGGAGAAAATAATACTGTTGATGGGTGTCTGGTCAGAGACATCAAGAAA
TAACGCCGGAACATTAGTGCAGGCAGCTTCCACAGCAATGGCATCCTGGTCATCCAGCGG
ATAGTTAATGATCAGCCCACTGACGCGTTGCGCGAGAAGATTGTGCACCGCCGCTTTACA

GGCTTCGACGCCGCTTCGTTCTACCATCGACACCACCACGCTGGCACCCAGTTGATCGGC
GCGAGATTTAATCGCCGCGACAATTTGCGACGGCGCGTGCAGGGCCAGACTGGAGGTGG
CAACGCCAATCAGCAACGACTGTTTGCCCGCCAGTTGTTGTGCCACGCGGTTGGGAATGT
AATTCAGCTCCGCCATCGCCGCTTCCACTTTTTCCCGCGTTTTCGCAGAAACGTGGCTGGC
CTGGTTCACCACGCGGGAAACGGTCTGATAAGAGACACCGGCATACTCTGCGACATCGTA
TAACGTTACTGGTTTCACATTCACCACCCTGAATTGACTCTCTTCCGGGCGCTATCATGCCA
TACCGCGAAAGGTTTTGCGCCATTCGATGGTGTCCGGGATCTCGACGCTCTCCCTTATGC
GACTCCTGCATTAGGAAGCAGCCCAGTAGTAGGTTGAGGCCGTTGAGCACCGCCGCCGC
AAGGAATGGTGCATGCAAGGAGATGGCGCCCAACAGTCCCCCGGCCACGGGGCCTGCCA
CCATACCCACGCCGAAACAAGCGCTCATGAGCCCGAAGTGGCGAGCCCGATCTTCCCCAT
CGGTGATGTCGGCGATATAGGCGCCAGCAACCGCACCTGTGGCGCCGGTGATGCCGGCC
ACGATGCGTCCGGCGTAGAGGATCGAGATCTCGATCCCGCGAAATTAATACGACTCACTAT
AGGGGAATTGTGAGCGGATAACAATTCCCCTCTAGAAATAATTTTGTTTAACTTTAAGAAGG
AGATATACCATGGGCAGCAGCCATCATCATCATCATCACAGCAGCGGCGAGAATCTGTACT
TCCAGAGTGTGAATTACGCGGCGGGGCTGTCGCCGTACGCGGACAAGGGCAAGTGCGGC
CTCCCGGAGATCTTCGACCCCCCGGAGGAGCTGGAGCGGAAGGTGTGGGAACTGGCGAG
GCTGGTCTGGCAGTCTTCCAGTGTGGTGTTCCACACGGGTGCCGGCATCAGCACTGCCTC
TGGCATCCCCGACTTCAGGGGTCCCCACGGAGTCTGGACCATGGAGGAGCGAGGTCTGG
CCCCCAAGTTCGACACCACCTTTGAGAGCGCGCGGCCCACGCAGACCCACATGGCGCTG
GTGCAGCTGGAGCGCGTGGGCCTCCTCCGCTTCCTGGTCAGCCAGAACGTGGACGGGCT
CCATGTGCGCTCAGGCTTCCCCAGGGACAAACTGGCAGAGCTCCACGGGAACATGTTTGT
GGAAGAATGTGCCAAGTGTAAGACGCAGTACGTCCGAGACACAGTCGTGGGCACCATGG
GCCTGAAGGCCACGGGCCGGCTCTGCACCGTGGCTAAGGCAAGGGGGCTGCGAGCCTG
CAGGGGAGAGCTGAGGGACACCATCCTAGACTGGGAGGACTCCCTGCCCGACCGGGACC
TGGCACTCGCCGATGAGGCCAGCAGGAACGCCGACCTGTCCATCACGCTGGGTACATCG
CTGCAGATCCGGCCCAGCGGGAACCTGCCGCTGGCTACCAAGCGCCGGGGAGGCCGCC
TGGTCATCGTCAACCTGCAGCCCACCAAGCACGACCGCCATGCTGACCTCCGCATCCATG
GCTACGTTGACGAGGTCATGACCCGGCTCATGAAGCACCTGGGGCTGGAGATCCCCGCC
TGGGACGGCCCCCGTGTGCTGGAGAGGGCGCTGCCACCCCTGCCCCGCCCGCCCACCC
CCAAGCTGGAGCCCAAGGAGGAATCTCCCACCCGGATCAACGGCTCTATCCCCGCCGGC
CCCAAGCAGGAGCCCTGCGCCCAGCACAACGGCTCAGAGCCCGCCAGCCCCAAACGGGA
GCGGCCCACCAGCCCTGCCCCCCACAGACCCCCCAAAAGGGTGAAGGCCAAGGCGGTCC
CCAGCTGATGACGAAGCTTGCGGCCGCACTCGAGCACCACCACCACCACCACTGAGATCC
GGCTGCTAACAAAGCCCGAAAGGAAGCTGAGTTGGCTGCTGCCACCGCTGAGCAATAACT
AGCATAACCCCTTGGGGCCTCTAAACGGGTCTTGAGGGGTTTTTTGCTGAAAGGAGGAAC
TATATCCGGATTGGCGAATGGGACGCGCCCTGTAGCGGCGCATTAAGCGCGGCGGGTGT
GGTGGTTACGCGCAGCGTGACCGCTACACTTGCCAGCGCCCTAGCGCCCGCTCCTTTCG
CTTTCTTCCCTTCCTTTCTCGCCACGTTCGCCGGCTTTCCCCGTCAAGCTCTAAATCGGGG
GCTCCCTTTAGGGTTCCGATTTAGTGCTTTACGGCACCTCGACCCCAAAAAACTTGATTAG
GGTGATGGTTCACGTAGTGGGCCATCGCCCTGATAGACGGTTTTTCGCCCTTTGACGTTG
GAGTCCACGTTCTTTAATAGTGGACTCTTGTTCCAAACTGGAACAACACTCAACCCTATCTC
GGTCTATTCTTTTGATTTATAAGGGATTTTGCCGATTTCGGCCTATTGGTTAAAAAATGAGC
TGATTTAACAAAAATTTAACGCGAATTTTAACAAAATATTAACGCTTACAATTTAGGTGGCAC
TTTTCGGGGAAATGTGCGCGGAACCCCTATTTGTTTATTTTTCTAAATACATTCAAATATGTA
TCCGCTCATGAATTAATTCTTAGAAAAACTCATCGAGCATCAAATGAAACTGCAATTTATTCA
TATCAGGATTATCAATACCATATTTTTGAAAAAGCCGTTTCTGTAATGAAGGAGAAAACTCA
CCGAGGCAGTTCCATAGGATGGCAAGATCCTGGTATCGGTCTGCGATTCCGACTCGTCCA
ACATCAATACAACCTATTAATTTCCCCTCGTCAAAAATAAGGTTATCAAGTGAGAAATCACC
ATGAGTGACGACTGAATCCGGTGAGAATGGCAAAAGTTTATGCATTTCTTTCCAGACTTGTT
CAACAGGCCAGCCATTACGCTCGTCATCAAAATCACTCGCATCAACCAAACCGTTATTCATT
CGTGATTGCGCCTGAGCGAGACGAAATACGCGATCGCTGTTAAAAGGACAATTACAAACA

GGAATCGAATGCAACCGGCGCAGGAACACTGCCAGCGCATCAACAATATTTTCACCTGAAT
CAGGATATTCTTCTAATACCTGGAATGCTGTTTTCCCGGGGATCGCAGTGGTGAGTAACCA
TGCATCATCAGGAGTACGGATAAAATGCTTGATGGTCGGAAGAGGCATAAATTCCGTCAGC
CAGTTTAGTCTGACCATCTCATCTGTAACATCATTGGCAACGCTACCTTTGCCATGTTTCAG
AAACAACTCTGGCGCATCGGGCTTCCCATACAATCGATAGATTGTCGCACCTGATTGCCCG
ACATTATCGCGAGCCCATTTATACCCATATAAATCAGCATCCATGTTGGAATTTAATCGCGG
CCTAGAGCAAGACGTTTCCCGTTGAATATGGCTCATAACACCCCTTGTATTACTGTTTATGT
AAGCAGACAGTTTTATTGTTCATGACCAAAATCCCTTAACGTGAGTTTTCGTTCCACTGAGC
GTCAGACCCCGTAGAAAAGATCAAAGGATCTTCTTGAGATCCTTTTTTTCTGCGCGTAATCT
GCTGCTTGCAAACAAAAAAACCACCGCTACCAGCGGTGGTTTGTTTGCCGGATCAAGAGCT
ACCAACTCTTTTTCCGAAGGTAACTGGCTTCAGCAGAGCGCAGATACCAAATACTGTCCTT
CTAGTGTAGCCGTAGTTAGGCCACCACTTCAAGAACTCTGTAGCACCGCCTACATACCTCG
CTCTGCTAATCCTGTTACCAGTGGCTGCTGCCAGTGGCGATAAGTCGTGTCTTACCGGGTT
GGACTCAAGACGATAGTTACCGGATAAGGCGCAGCGGTCGGGCTGAACGGGGGGTTCGT
GCACACAGCCCAGCTTGGAGCGAACGACCTACACCGAACTGAGATACCTACAGCGTGAGC
TATGAGAAAGCGCCACGCTTCCCGAAGGGAGAAAGGCGGACAGGTATCCGGTAAGCGGC
AGGGTCGGAACAGGAGAGCGCACGAGGGAGCTTCCAGGGGGAAACGCCTGGTATCTTTA
TAGTCCTGTCGGGTTTCGCCACCTCTGACTTGAGCGTCGATTTTTGTGATGCTCGTCAGGG
GGGCGGAGCCTATGGAAAAACGCCAGCAACGCGGCCTTTTTACGGTTCCTGGCCTTTTGC
TGGCCTTTTGCTCACATGTTCTTTCCTGCGTTATCCCCTGATTCTGTGGATAACCGTATTAC
CGCCTTTGAGTGAGCTGATACCGCTCGCCGCAGCCGAACGACCGAGCGCAGCGAGTCAG
TGAGCGAGGAAGCGGAAGAGCGCCTGATGCGGTATTTTCTCCTTACGCATCTGTGCGGTA
TTTCACACCGCAATGGTGCACTCTCAGTACAATCTGCTCTGATGCCGCATAGTTAAGCCAG
TATACACTCCGCTATCGCTACGTGACTGGGTCATGGCTGCGCCCCGACACCCGCCAACAC
CCGCTGACGCGCCCTGACGGGCTTGTCTGCTCCCGGCATCCGCTTACAGACAAGCTGTGA
CCGTCTCCGGGAGCTGCATGTGTCAGAGGTTTTCACCGTCATCACCGAAACGCGCGAGGC
AGCTGCGGTAAAGCTCATCAGCGTGGTCGTGAAGCGATTCACAGATGTCTGCCTGTTCATC
CGCGTCCAGCTCGTTGAGTTTCTCCAGAAGCGTTAATGTCTGGCTTCTGATAAAGCGGGCC
ATGTTAAGGGCGGTTTTTTCCTGTTTGGTCACTGATGCCTCCGTGTAAGGGGGATTTCTGT
TCATGGGGGTAATGATACCGATGAAACGAGAGAGGATGCTCACGATACGGGTTACTGATG
ATGAACATGCCCGGTTACTGGAACGTTGTGAGGGTAAACAACTGGCGGTATGGATGCGGC
GGGACCAGAGAAAAATCACTCAGGGTCAATGCCAGCGCTTCGTTAATACAGATGTAGGTGT
TCCACAGGGTAGCCAGCAGCATCCTGCGATGCAGATCCGGAACATAATGGTGCAGGGCG
CTGACTTCCGCGTTTCCAGACTTTACGAAACACGGAAACCGAAGACCATTCATGTTGTTGC
TCAGGTCGCAGACGTTTTGCAGCAGCAGTCGCTTCACGTTCGCTCGCGTATCGGTGATTC
ATTCTGCTAACCAGTAAGGCAACCCCGCCAGCCTAGCCGGGTCCTCAACGACAGGAGCAC
GATCATGCGCACCCGTGGGGCCGCCATGCCGGCGATAATGGCCTGCTTCTCGCCGAAAC
GTTTGGTGGCGGGACCAGTGACGAAGGCTTGAGCGAGGGCGTGCAAGATTCCGAATACC
GCAAGCGACAGGCCGATCATCGTCGCGCTCCAGCGAAAGCGGTCCTCGCCGAAAATGAC
CCAGAGCGCTGCCGGCACCTGTCCTACGAGTTGCATGATAAAGAAGACAGTCATAAGTGC
GGCGACGATAGTCATGCCCCGCGCCCACCGGAAGGAGCTGACTGGGTTGAAGGCTCTCA
AGGGCATCGGTCGAGATCCCGGTGCCTAATGAGTGAGCTAACTTACATTAATTGCGTTGCG
CTCACTGCCCGCTTTCCAGTCGGGAAACCTGTCGTGCCAGCTGCATTAATGAATCGGCCA
ACGCGCGGGGAGAGGCGGTTTGCGTATTGGGCGCCAGGGTGGTTTTTCTTTTCACCAGTG
AGACGGGCAACAGCTGATTGCCCTTCACCGCCTGGCCCTGAGAGAGTTGCAGCAAGCGG
TCCACGCTGGTTTGCCCCAGCAGGCGAAAATCCTGTTTGATGGTGGTTAACGGCGGGATA
TAACATGAGCTGTCTTCGGTATCGTCGTATCCCACTACCGAGATAT
