## Supplementary material for "DNA stimulates SIRT6 to mono-ADP-ribosylate proteins within histidine repeats": Data File S3

Creative Proteomics

SUITE 115, 17 Ramsey Road, Shirley, NY 11967, USA
www.creative-proteomics.com


### Analysis Report

| | |
|---|---|
| **Project Name** | Protein ADP-ribosylation Analysis |
| **Sample Description** | Protein sample in solution |
| **Sample Quantity** | One sample |
| **Project Client** | Katharine Diehl |
| **Project Date** | March 24, 2023 |
| **Order Number** | Order-CPJS09152205 |
| **Remark** | |

Creative Proteomics

SUITE 115, 17 Ramsey Road, Shirley, NY 11967, USA
www.creative-proteomics.com


### Analytical Service Report

#### 1. Sample Information

One protein sample was ran on SDS-PAGE gel followed by in-gel digested by trypsin, and then identified by applying our nanoLC-MS/MS platform.

#### 2. Materials and Methods

##### 2.1 Chemicals and Instrumentation

DL-dithiothreitol (DTT), iodoacetamide (IAA), formic acid (FA), acetonitrile (ACN), were purchased from Sigma (St. Louis, MO, USA), trypsin was purchased from Promega (Madison, WI, USA). Ultrapure water was prepared from a Millipore purification system (Billerica, MA, USA). An Ultimate 3000 nano UHPLC system coupled with a Q Exactive HF mass spectrometer (Thermo Fisher Scientific, USA) with an ESI nanospray source.

##### 2.2 SDS-PAGE

1) Load samples and protein marker on SDS-PAGE gel (12% separating gel) and make sure not to overflow. Then cover the top and connect the anodes.
2) Run the gel at 80 kV for 20 min. Then increase the voltage to 120 kV and run for 1 h.
3) Stain the SDS-PAGE gel with Coomassie Brilliant Blue.

##### 2.3 In-gel Digestion

1) Cut the gel slice from each gel into 1 mm3 cubes and transfer the gel cubes to a 1.5 mL microcentrifuge tube. Centrifuge the tube for 1-2 sec to spin the gel slice to the bottom of the tube. Add 1 mL 50 mM $NH_4HCO_3$: ACN =50 : 50. Remove the supernatant until the brown disappears. Stop the reaction by adding 200 μL of water. Discard the supernatant after 10 min. Add 1 mL of NH4HCO3 standing 30 min, and then remove the supernatant.
2) Add 500 μL of ACN and incubate for 30 min. The gel pieces should become opaque and stick together.
3) Remove the ACN and rehydrate the gel slice in 10 mM DTT. Add enough solution to completely cover the gel slice. Incubate at 56 ℃ for 1 h.
4) Carefully remove the DTT and add 500 μL of ACN, incubate for 10 min at room

SUITE 115, 17 Ramsey Road, Shirley, NY 11967, USA
www.creative-proteomics.com


temperature.
5) Remove the ACN and add 50 mM IAA to completely cover the gel slice. Incubate for 30 min at room temperature in the dark.
6) Remove the IAA and add 500 μL of ACN to the gel slice. Incubate for 10 min at room temperature. Carefully remove the ACN solution.
7) Add enough trypsin digestion solution to cover the gel slices. Incubate the gel pieces on ice for 45 min. Add more digestion solution if all the initial solution is absorbed by the gel pieces.
8) Remove the excess digestion solution and and 5-20 μL of 50 mM $NH_4HCO_3$ to keep the gel pieces wet during enzymatic digestion.
9) Recover the supernatant and transfer into a fresh 1.5 mL microcentrifuge tube. Add 50 mM ammonium bicarbonate/acetonitrile solution (1:2, v/v) to cover gel slices. Incubate for 1 h at 37 ℃. Transfer the solution to the 1.5 mL microcentrifuge tube.
10) Lyophilize the extracted peptides to near dryness.
11) Resuspend peptides in 20 μL of 0.1% formic acid before LC-MS/MS analysis.

**2.4 Nano LC-MS/MS Analysis**

**2.4.1 NanoLC**

Nanoflow UPLC: Ultimate 3000 nano UHPLC system (ThermoFisher Scientific, USA)
Nanocolumn: trapping column (PepMap C18, 100Å, 100 μm × 2 cm, 5 μm) and an analytical column (PepMap C18, 100Å, 75 μm × 50 cm, 2 μm)
Loaded sample: 1 μg
Mobile phase: A: 0.1% formic acid in water; B: 0.1% formic acid in 80% acetonitrile.
Total flow rate: 250 nL/min
LC linear gradient: from 2 to 8% buffer B in 3 min, from 8% to 20% buffer B in 50 min, from 20% to 40% buffer B in 36 min, then from 40% to 90% buffer B in 4 min.

**2.4.2 Mass Spectrometry**

The full scan was performed between 300-1,650 m/z at the resolution 60,000 at 200 m/z, the automatic gain control target for the full scan was set to 3e6. The MS/MS scan was operated in Top 20 mode using the following settings: resolution 15,000 at 200 m/z; automatic gain control target 1e5; maximum injection time 19ms; normalized collision

SUITE 115, 17 Ramsey Road, Shirley, NY 11967, USA
www.creative-proteomics.com


energy at 28%; isolation window of 1.4 Th; charge sate exclusion: unassigned, 1, > 6; dynamic exclusion 30 s.

##### 2.5 Data Analysis

Two raw MS files were analyzed and searched against MECP2 and SIRT6 reference sequences (provided by customers) using PEAKS STUDIO 8.5. The parameters were set as follows: the protein modifications were carbamidomethylation (C) (fixed), oxidation (M), ADP Ribose addition(variable); the enzyme specificity was set to trypsin; the maximum missed cleavages were set to 2; the precursor ion mass tolerance was set to 10 ppm, and MS/MS tolerance was 0.6 Da.

#### 3. Analytical Results

##### 3.1 SDS-PAGE

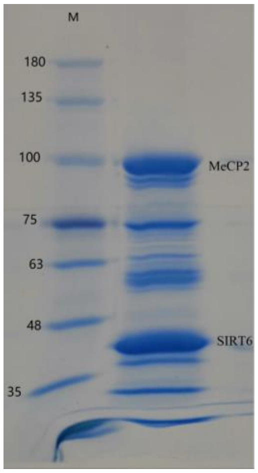


##### 3.2 MS based Protein Identification

The detailed identified peptides as well as ADP-ribosylation sites were listed in the supplemental excel sheets.
