## Supplementary material for "DNA stimulates SIRT6 to mono-ADP-ribosylate proteins within histidine repeats": Data File S4

Figure 1 A:

Main Text:
Anti-ADPr

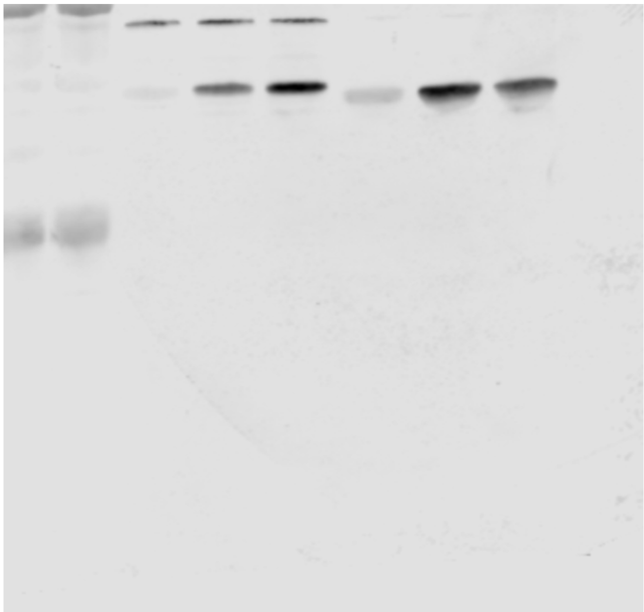

Total Protein

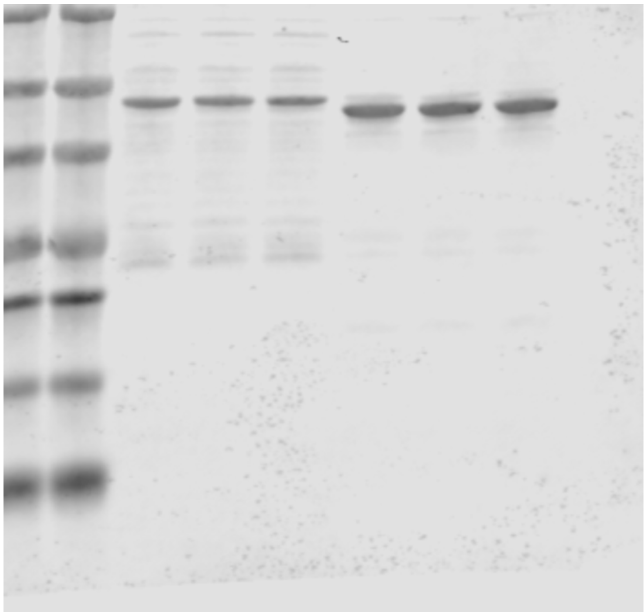

Figure 1 A:

Rep 2:

Anti-ADPr

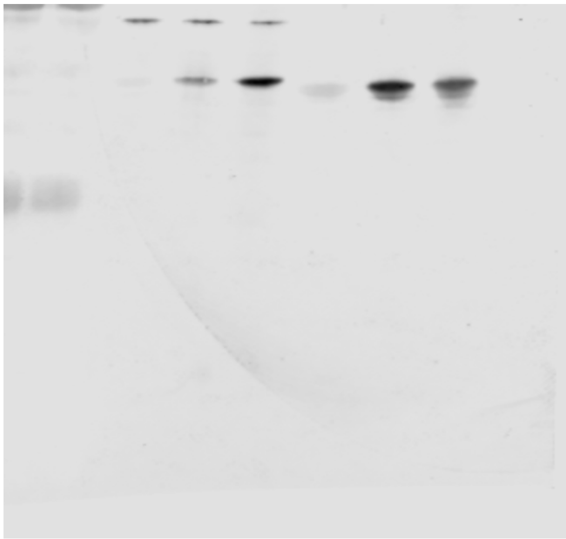

Total Protein

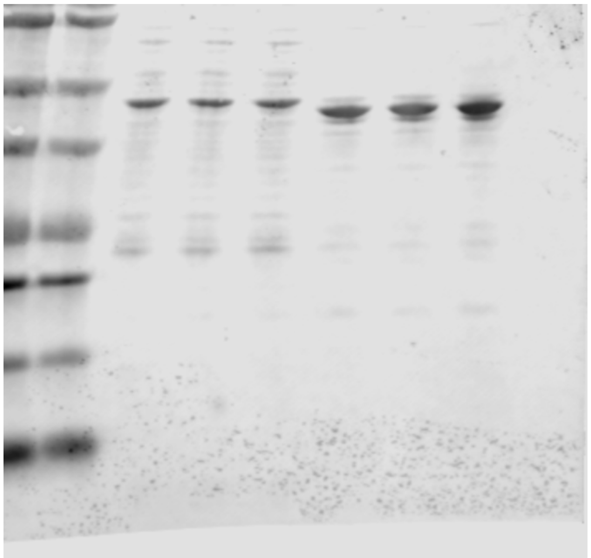

Figure 1 A:

Rep 3:

Anti-ADPr

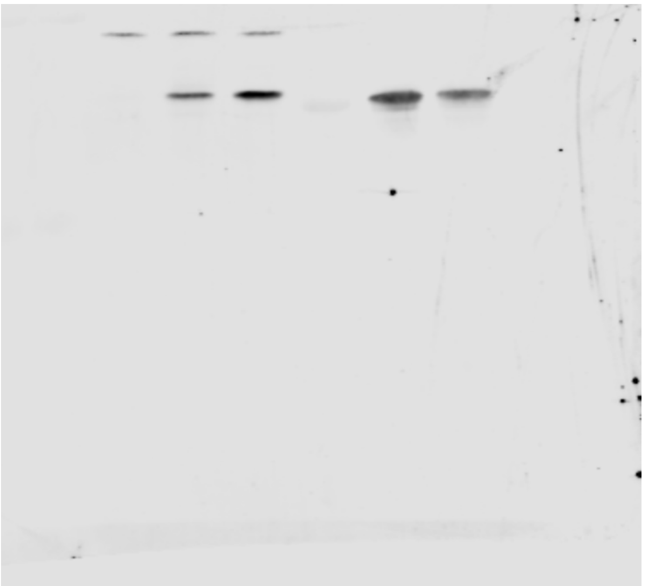

Total Protein

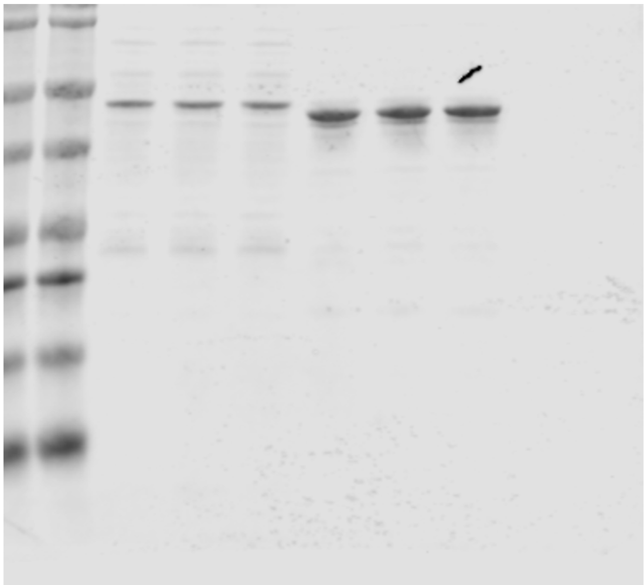

Figure 1 C:

Main Text:

Anti-ADPr

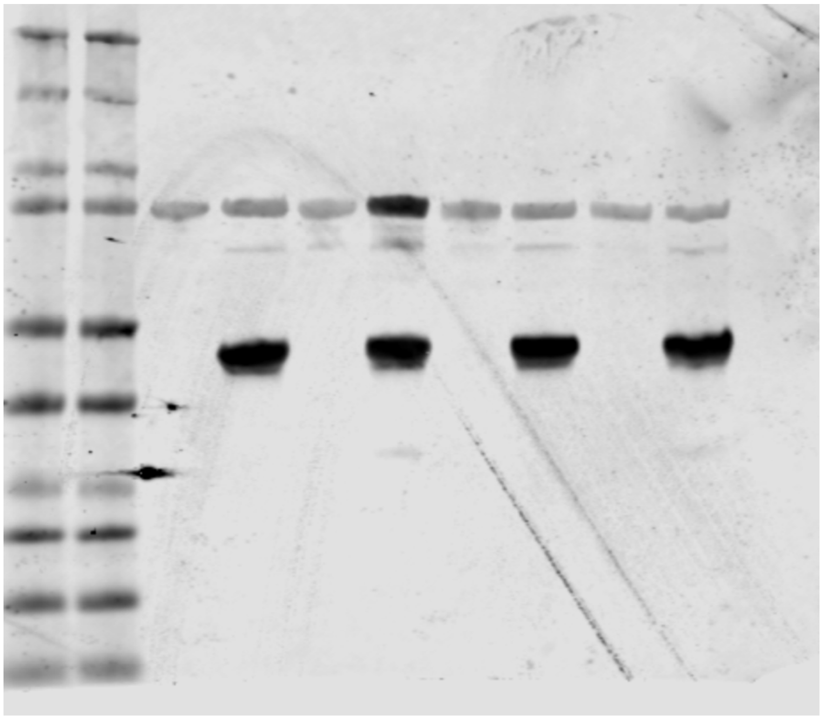

Total Protein

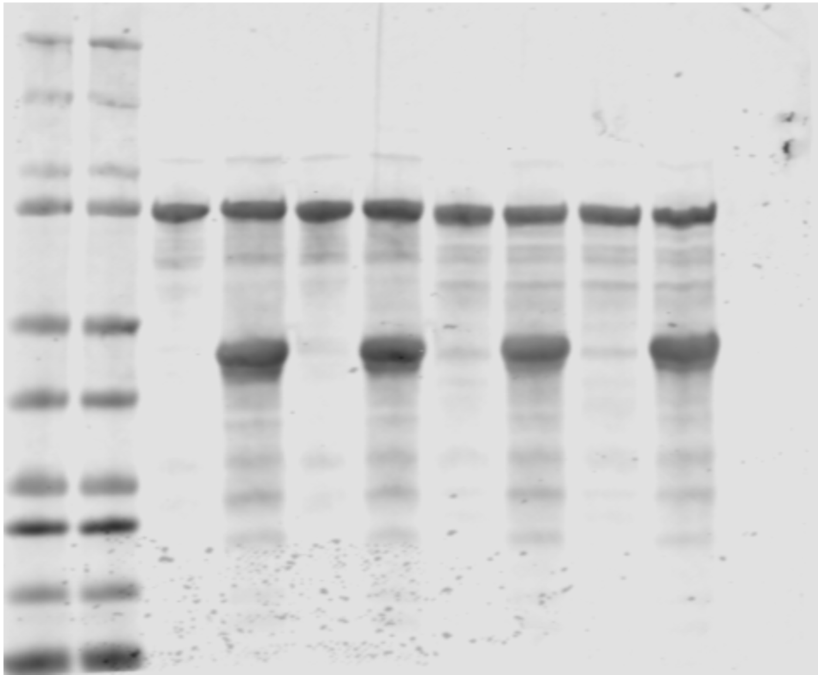

Figure 1 C:

Rep 2:
Anti-ADPr

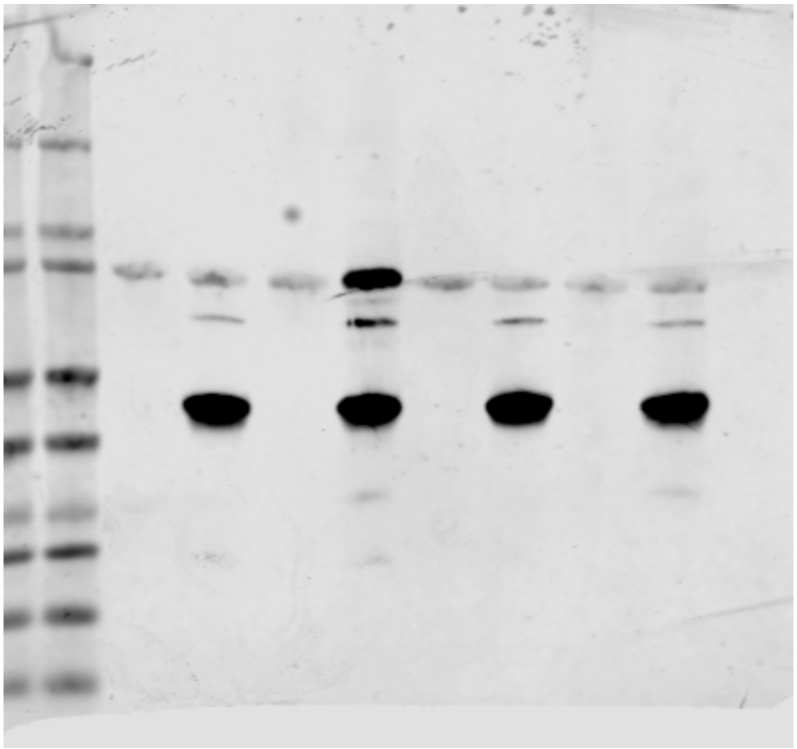

Total Protein

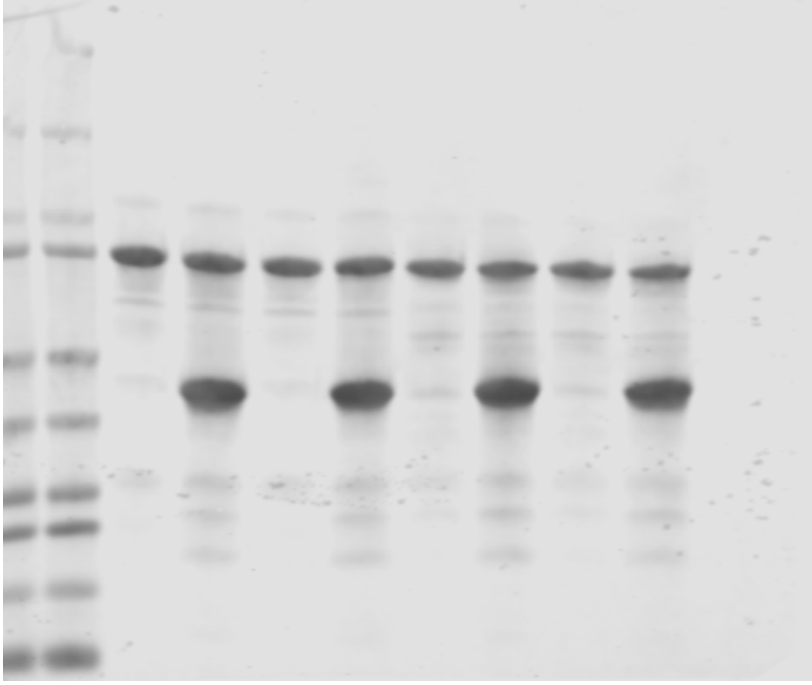

Figure 1 D:

Main Text:
Anti-ADPr

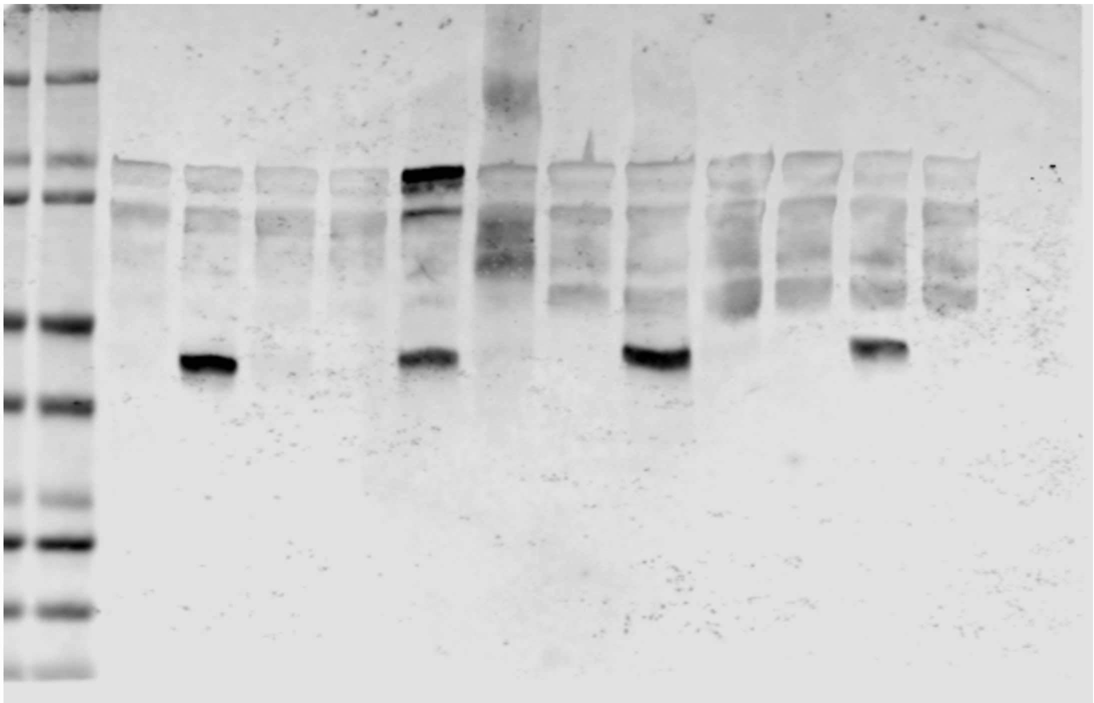

Total Protein

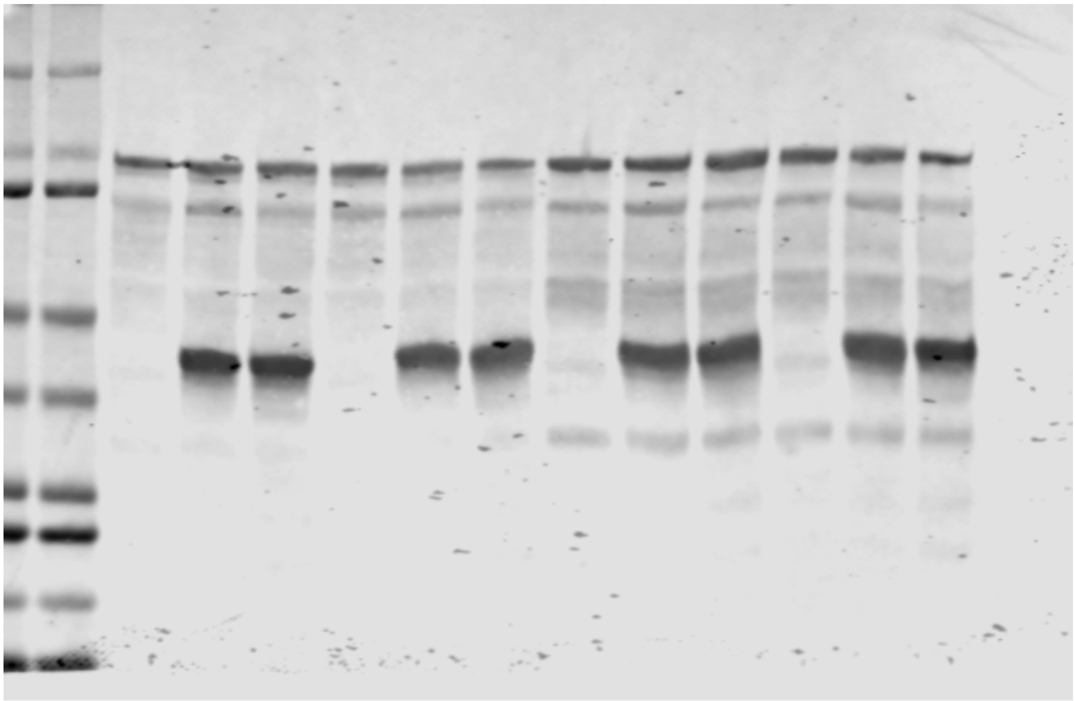

Figure 1 D:

Rep 2:

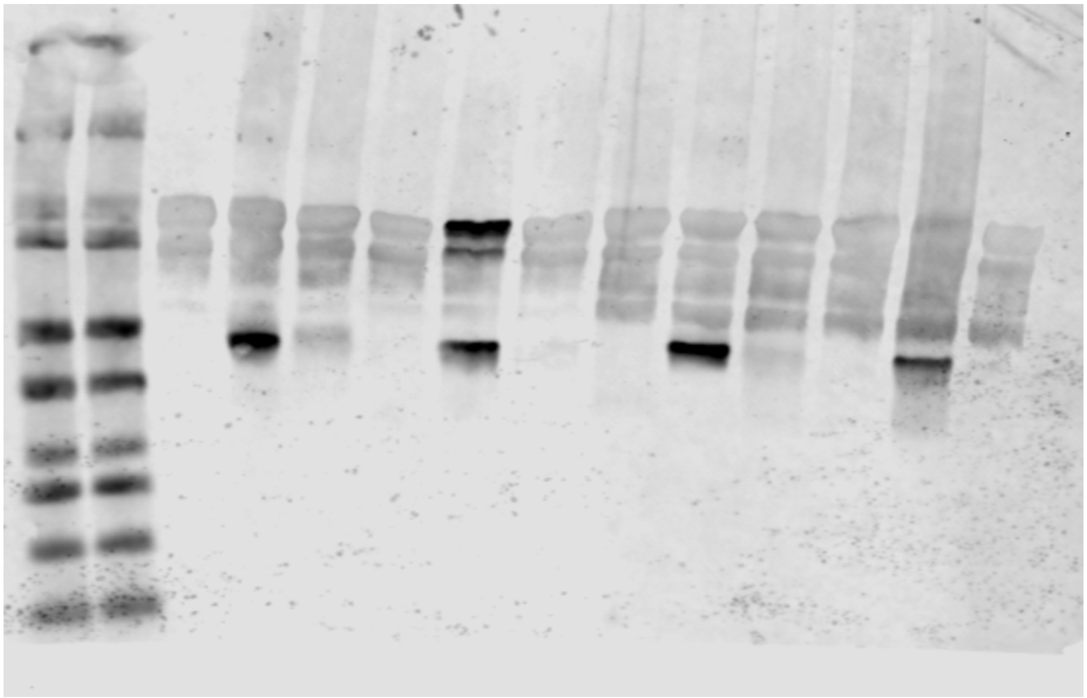

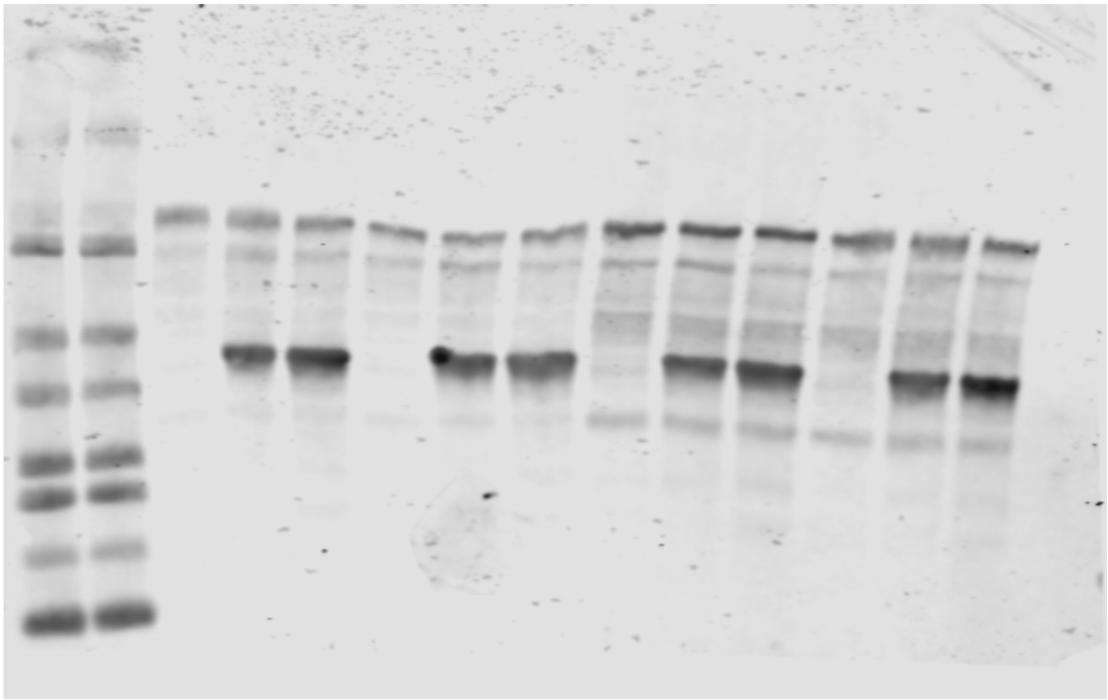

Figure 1 E:

Main Text:

Anti-ADPr

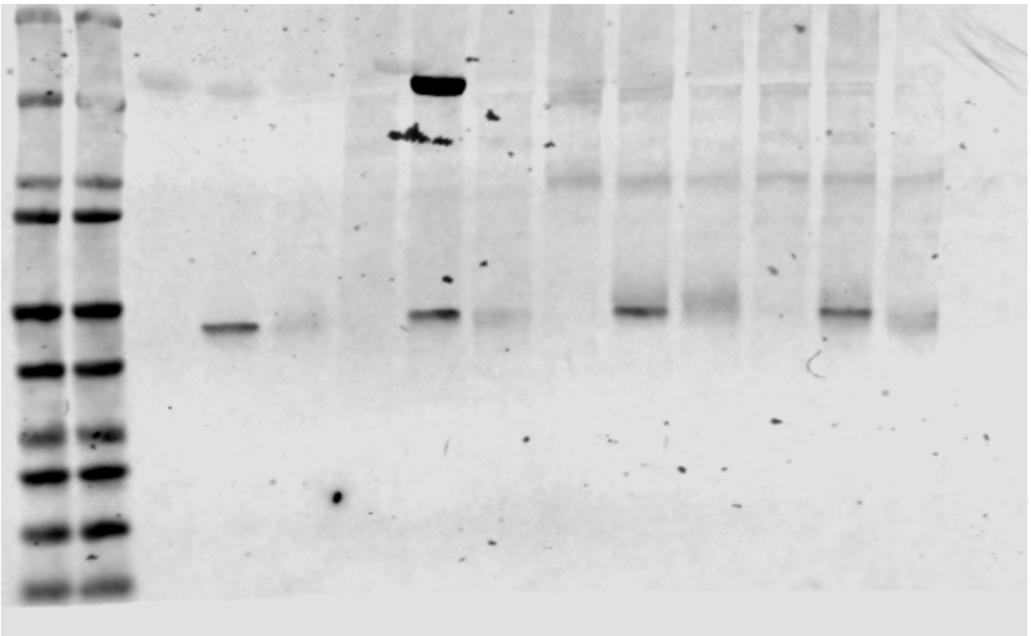

Total Protein

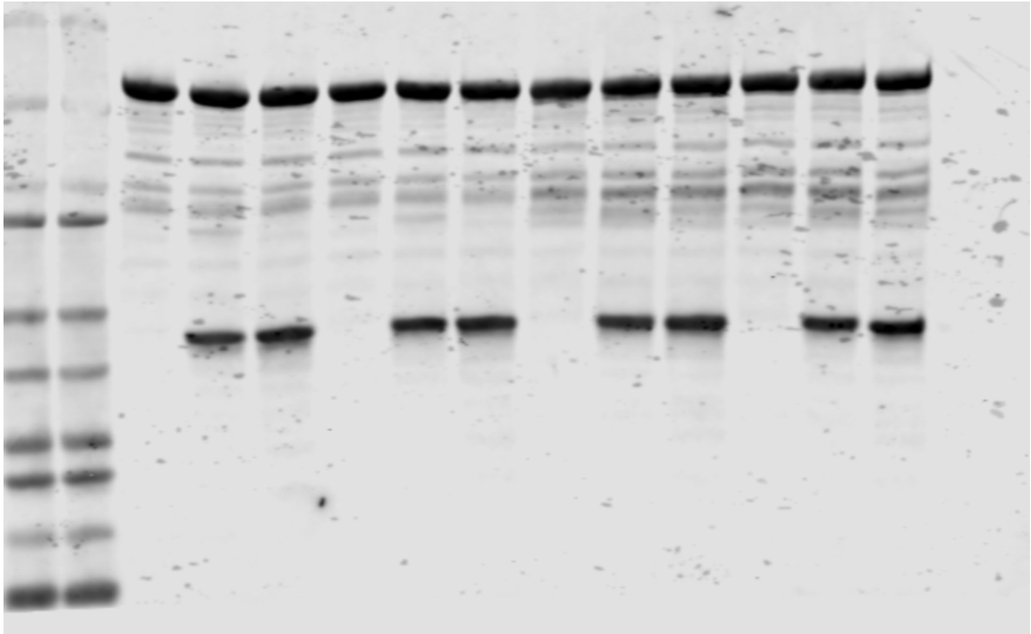

Figure 1 E:

Rep 2:

Anti-ADPr

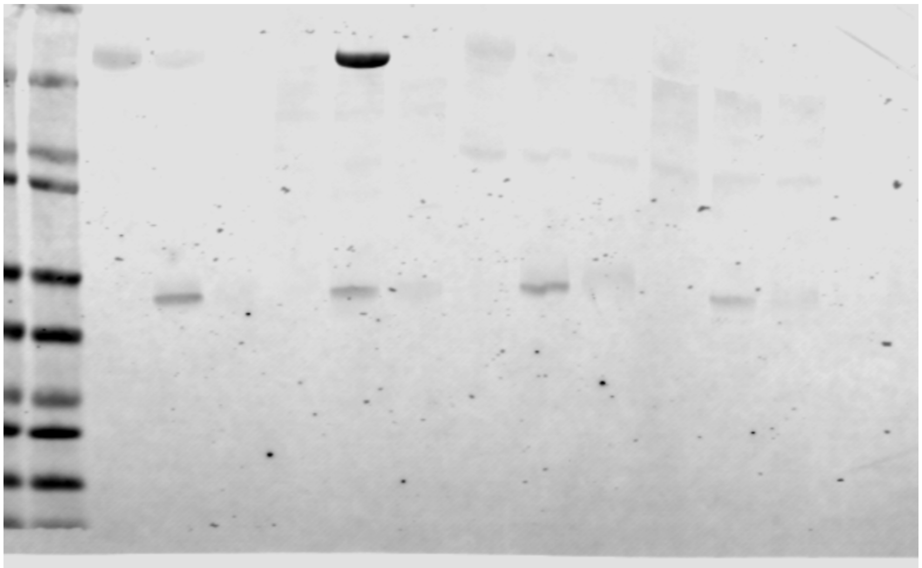

Total Protein

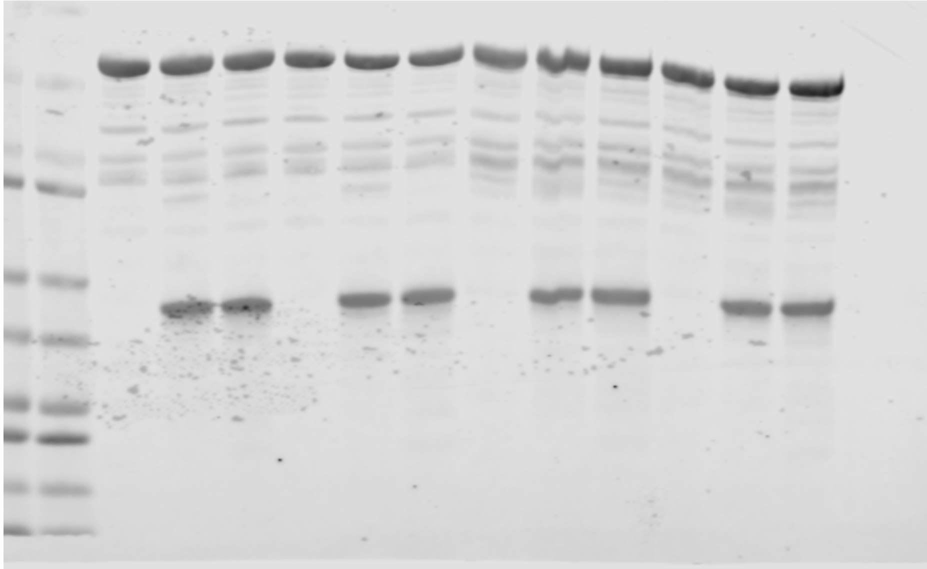

Figure 1 F:

Main Text:
Anti-ADPr

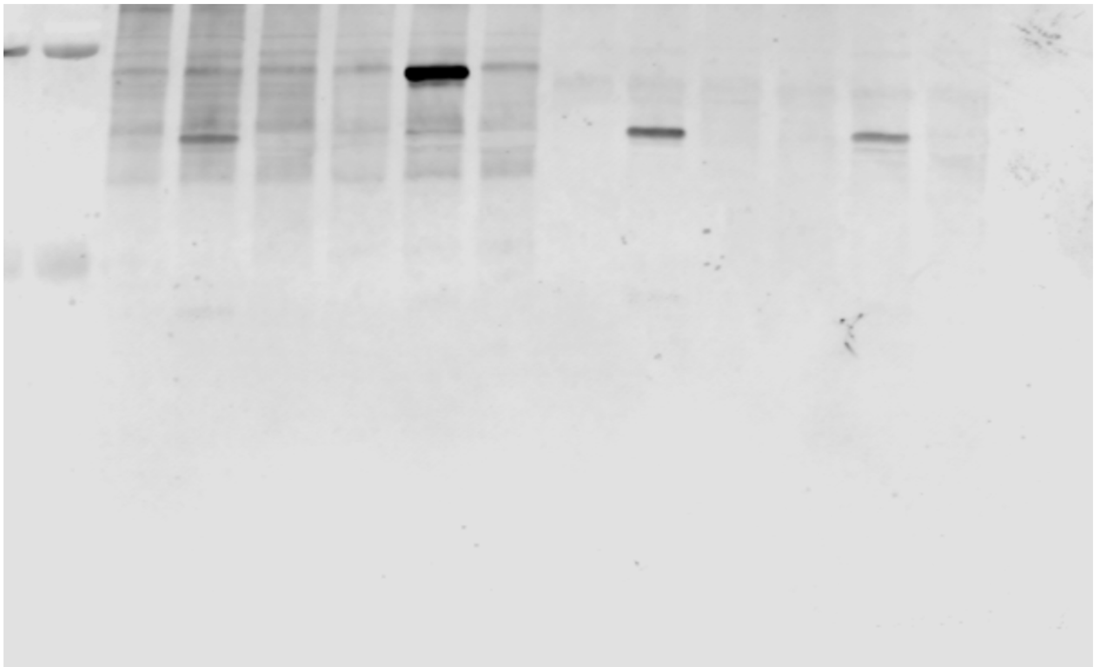

Total Protein

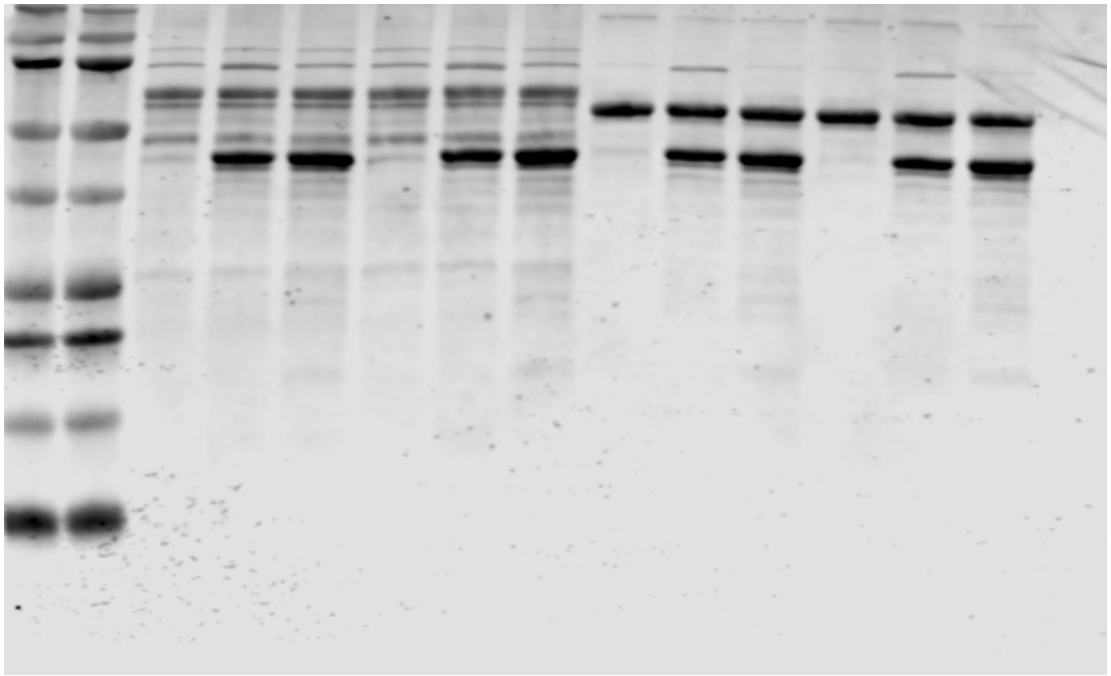

Figure 1 F:

Rep 2:
Anti-ADPr

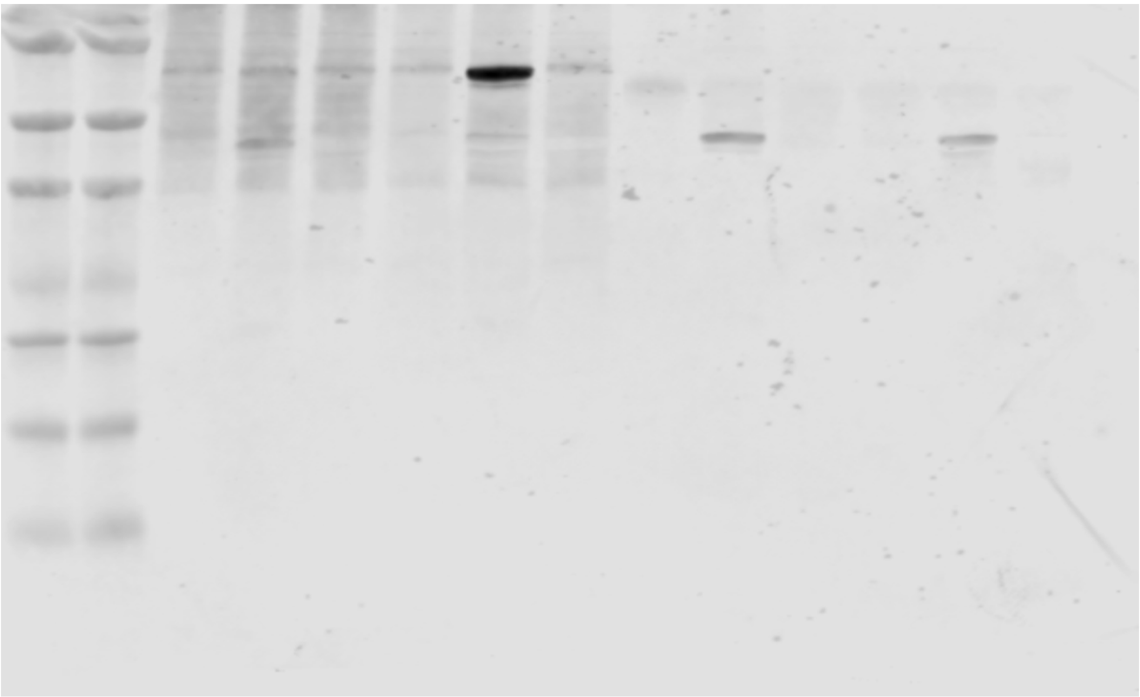

Total Protein

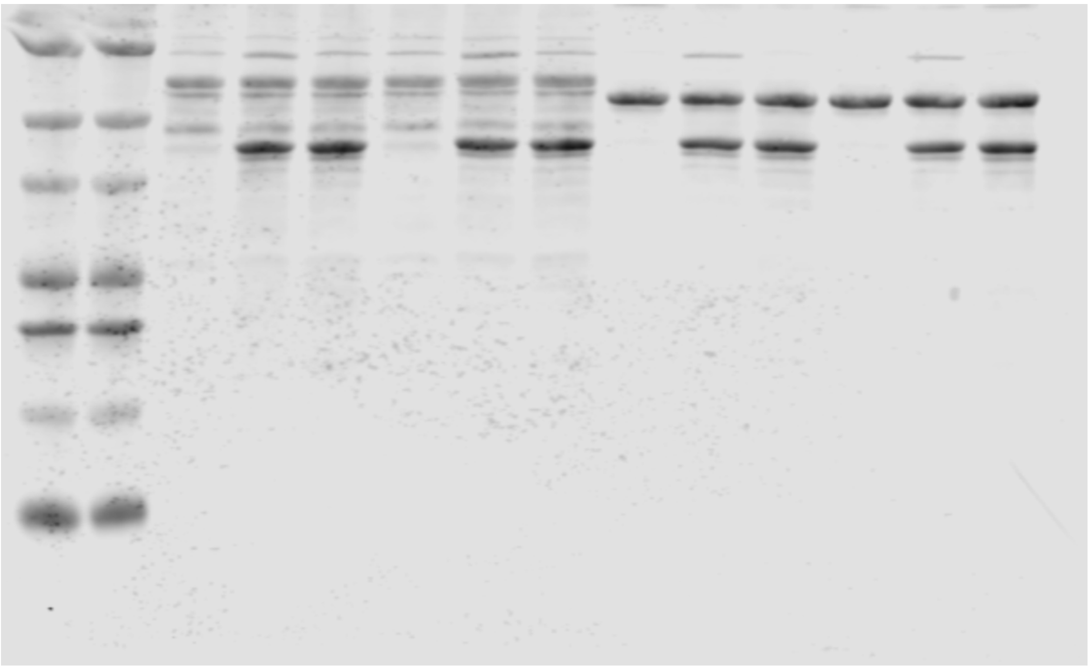

Figure S1 A:

Displayed image:
Anti-ADPr

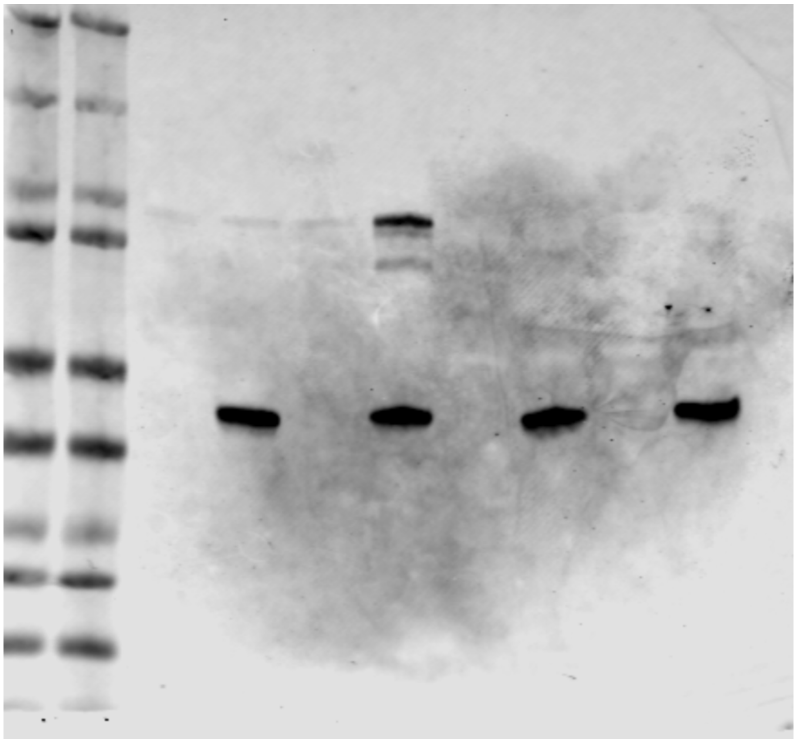

Total Protein

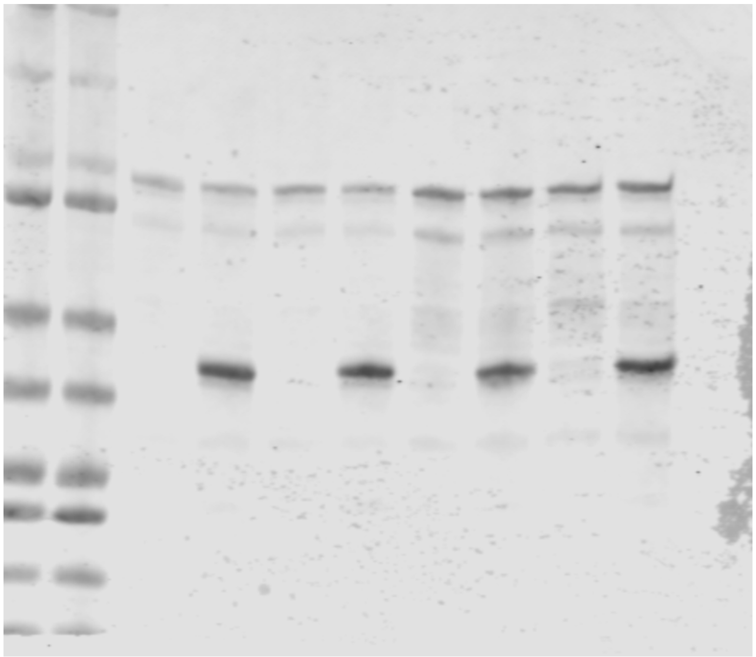

Figure S1 A:

Rep 2:
Anti-ADPr

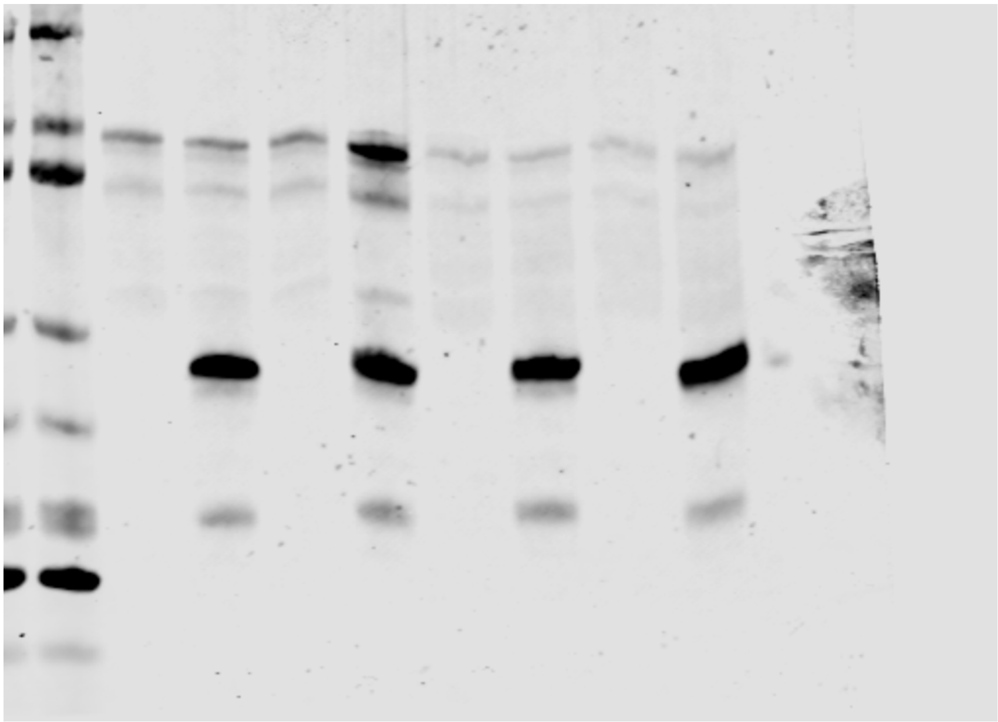

Total Protein

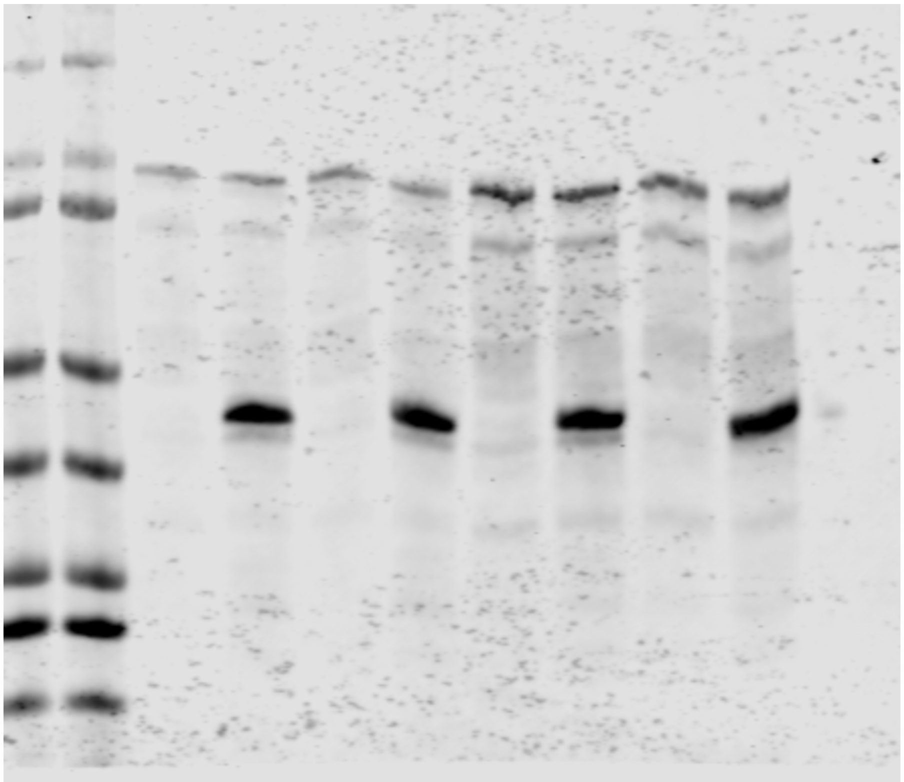

Figure 2 A:

Main Text:

Anti-ADPr (second half of blot technical replicate)

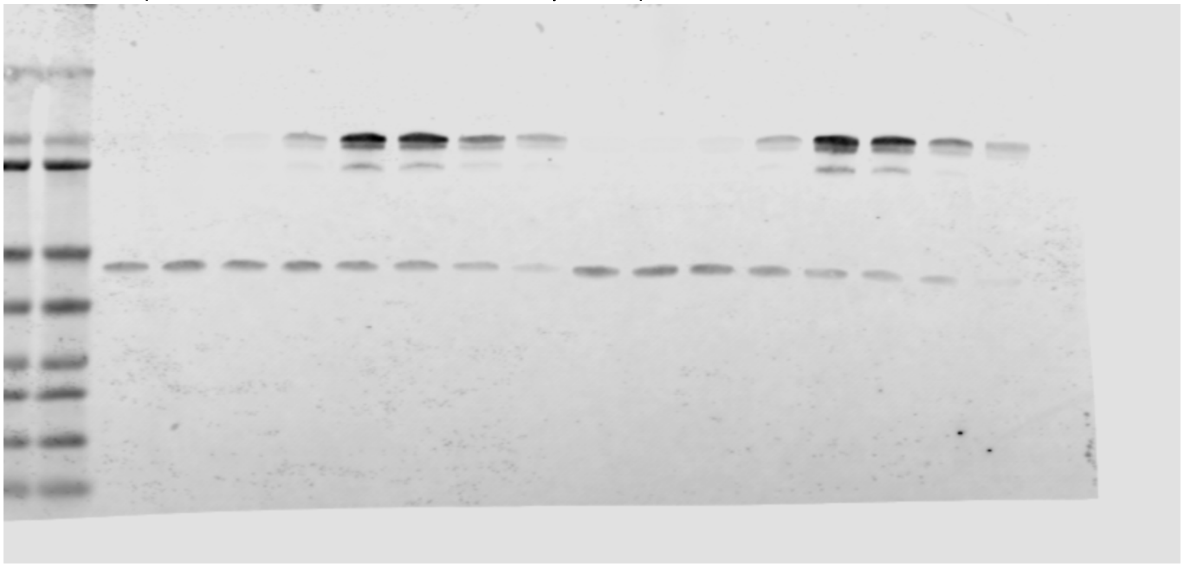

Total Protein (second half of blot technical replicate)

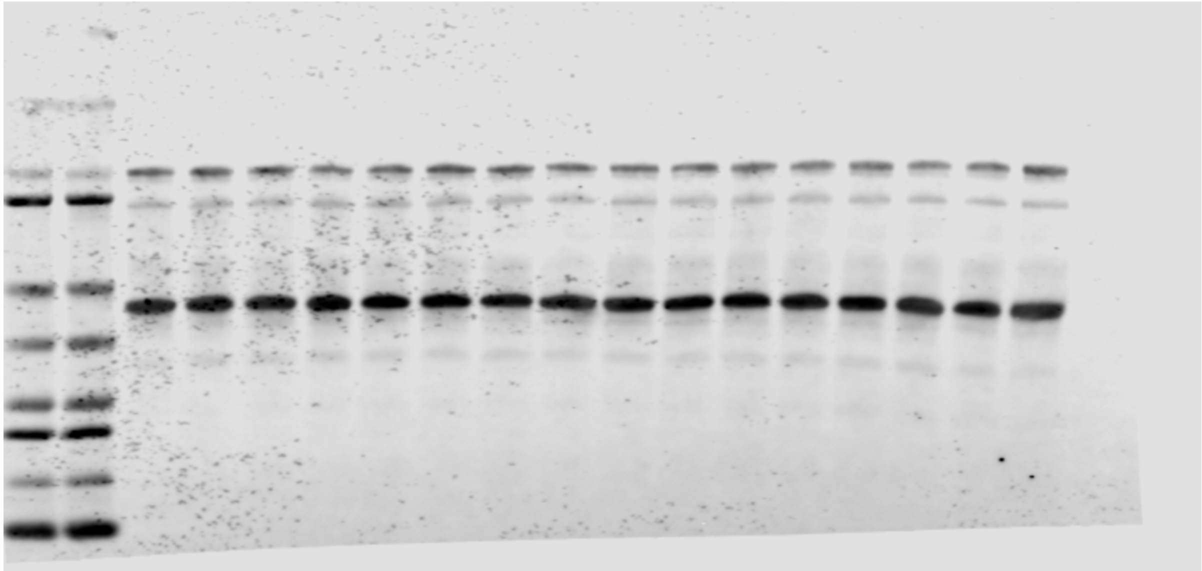

Figure 2 A:

Rep 2:
Anti-ADPr (second half of blot technical replicate)

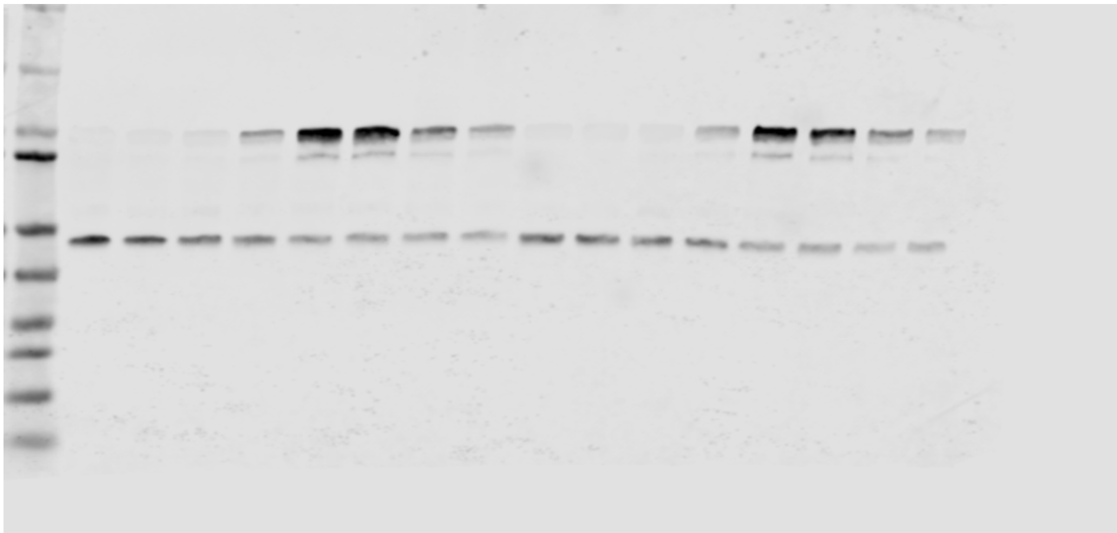

Total Protein (second half of blot technical replicate)

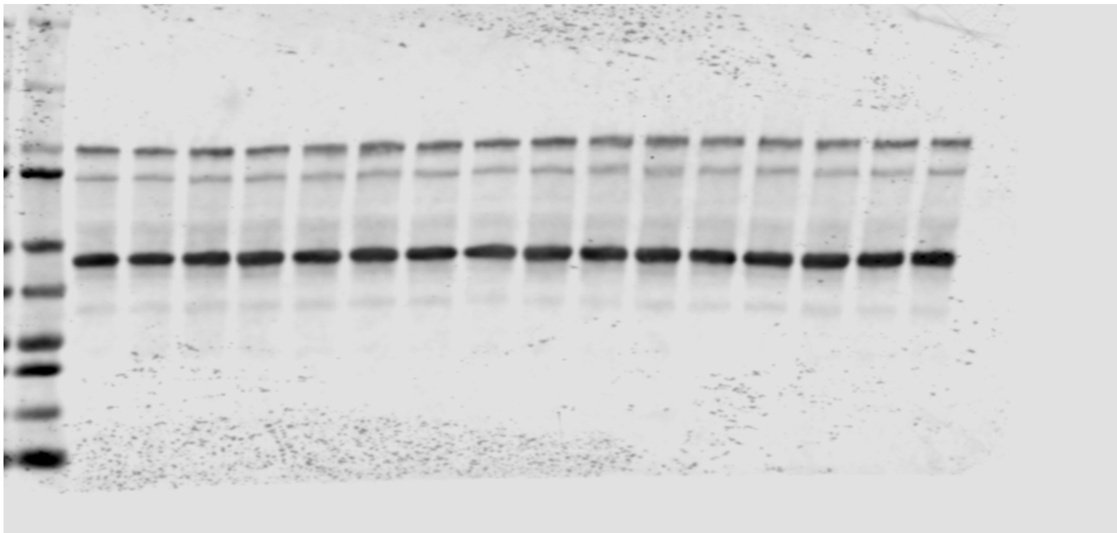

Figure 2 B:

Main Text:

Anti-ADPr (last 3 lanes not described or displayed in text)

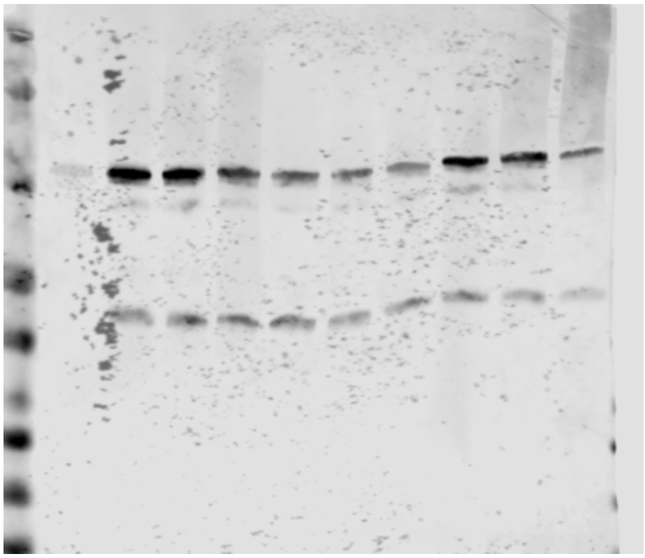

Total Protein (last 3 lanes not described or displayed in text)

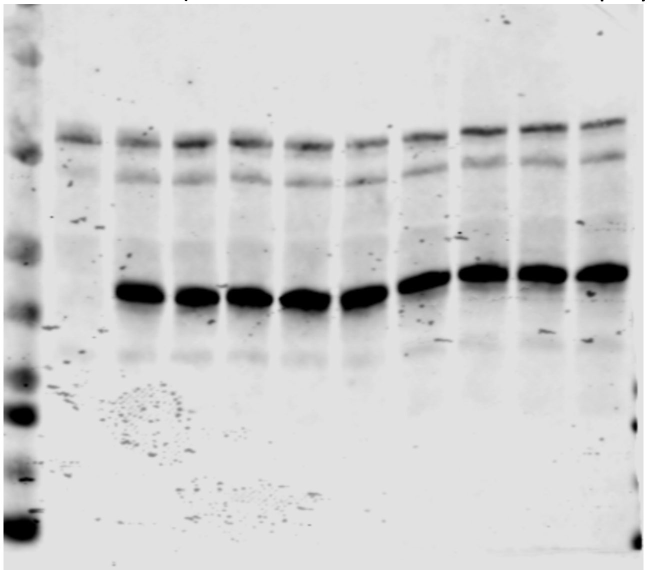

Figure 2 B:

Rep 2:

Anti-ADPr (last 3 lanes not described or displayed in text)

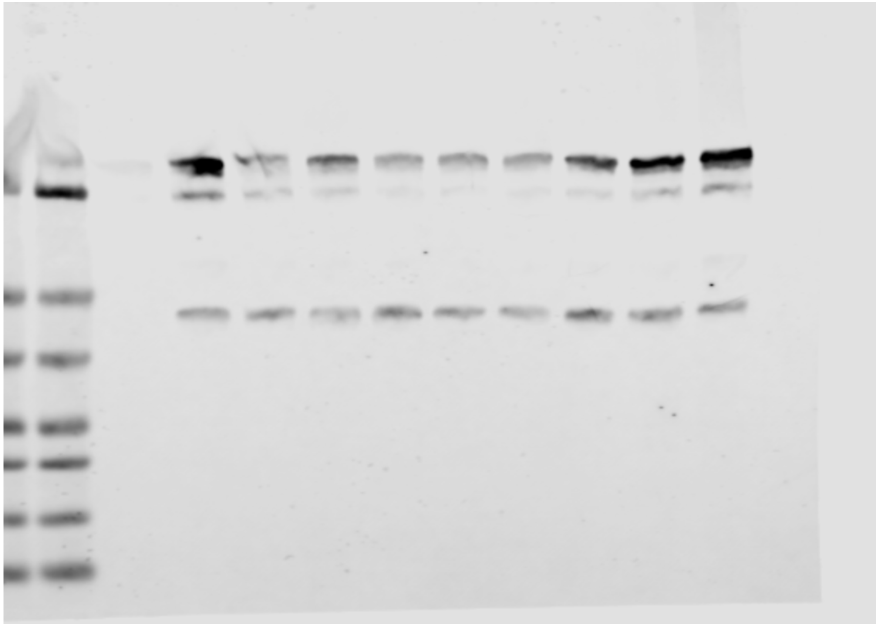

Total Protein (last 3 lanes not described or displayed in text)

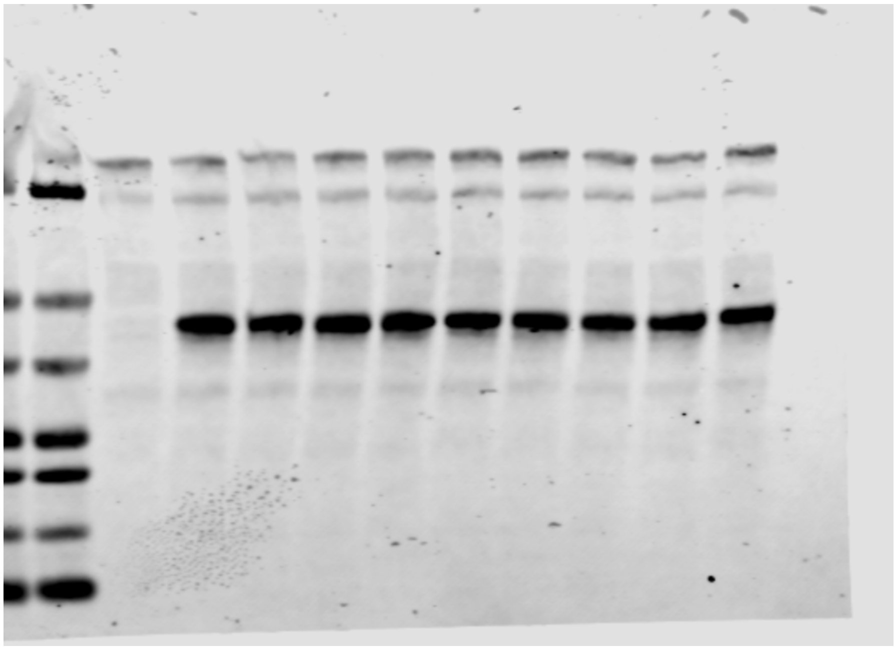

Figure 2 C:

Main Text:
Anti-ADPr

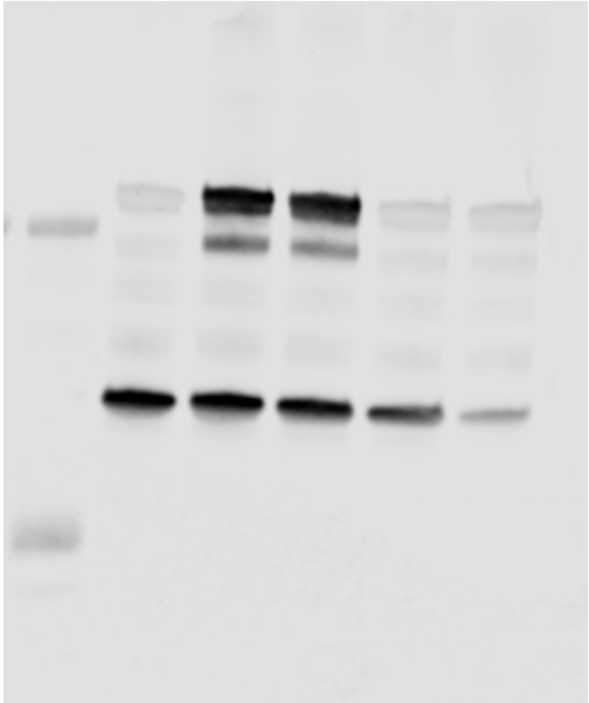

Total Protein

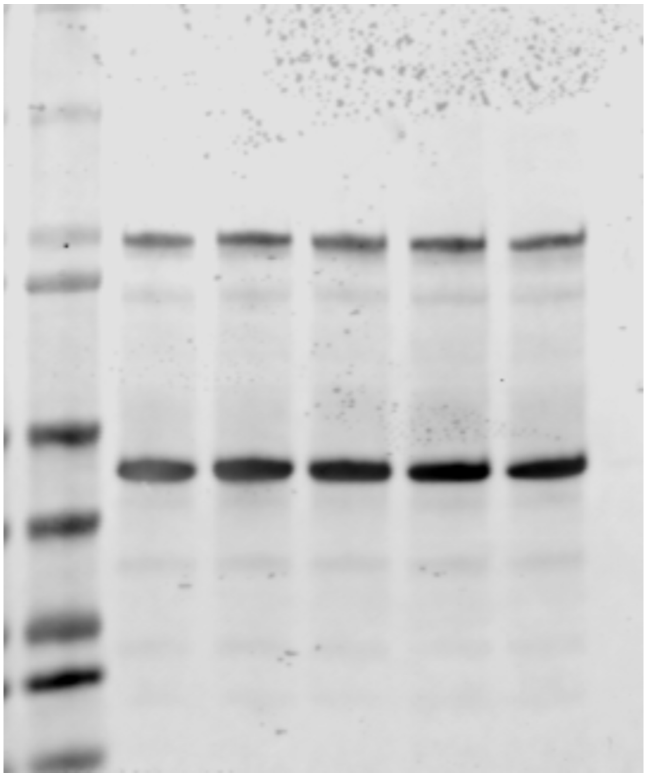

Figure 2 C:

Rep 2:
Anti-ADPr

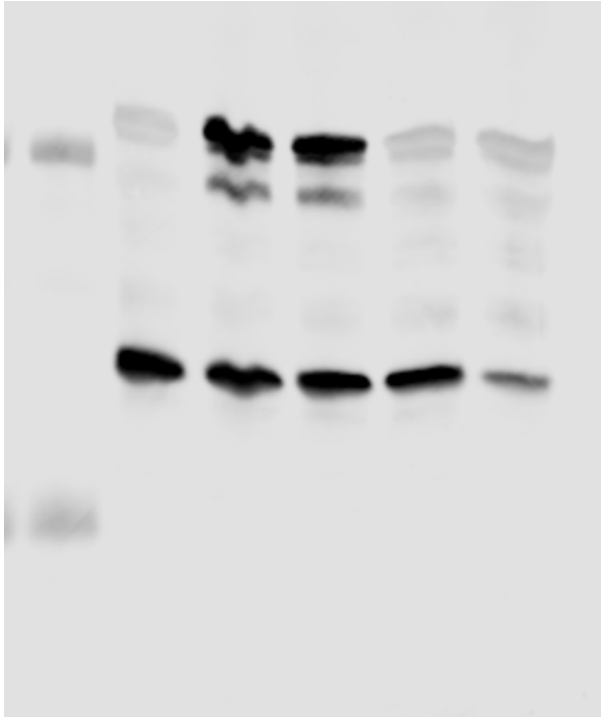

Total Protein

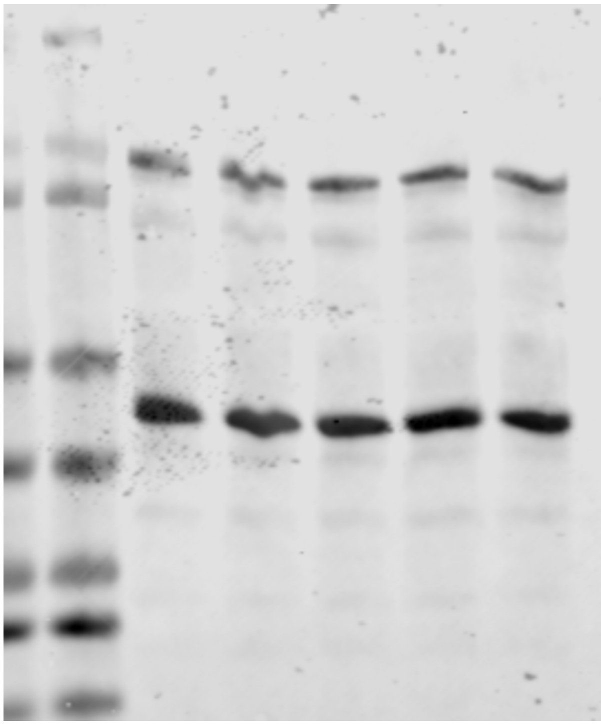

Figure 2 D:

Main Text:
Anti-ADPr

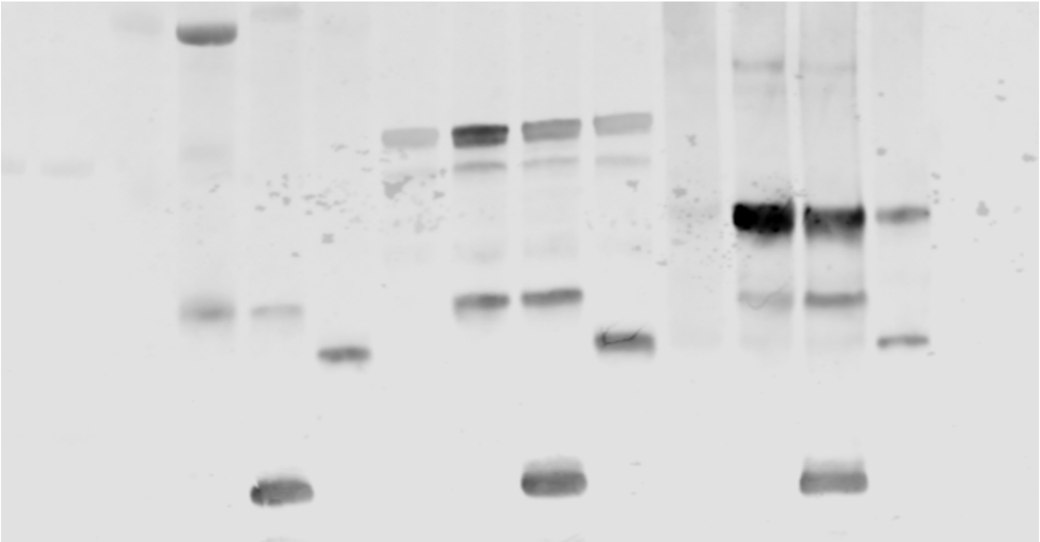

Total Protein

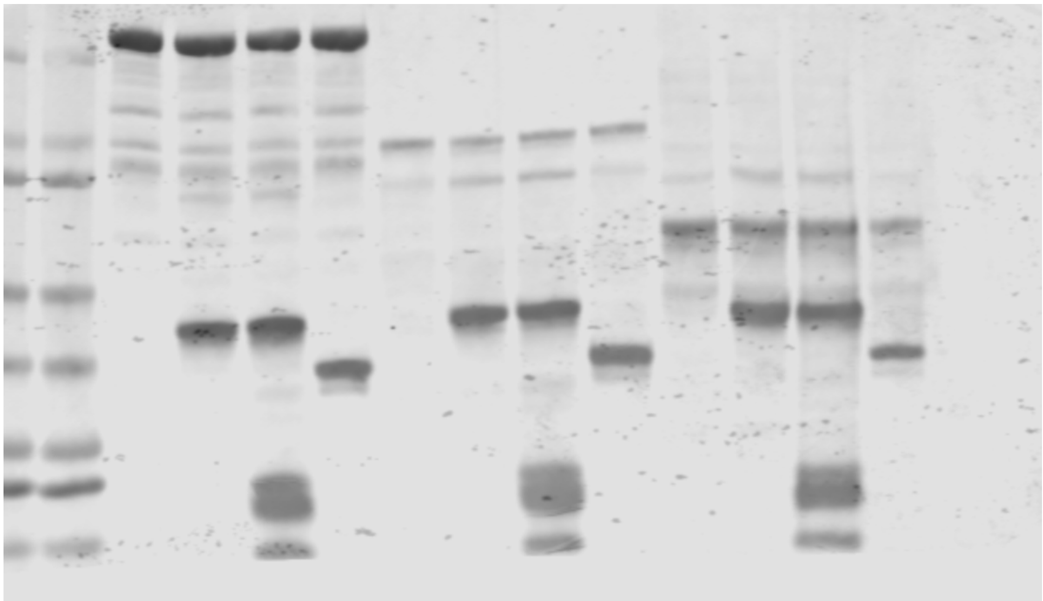

Figure 2 D:

Rep 2:
Anti-ADPr

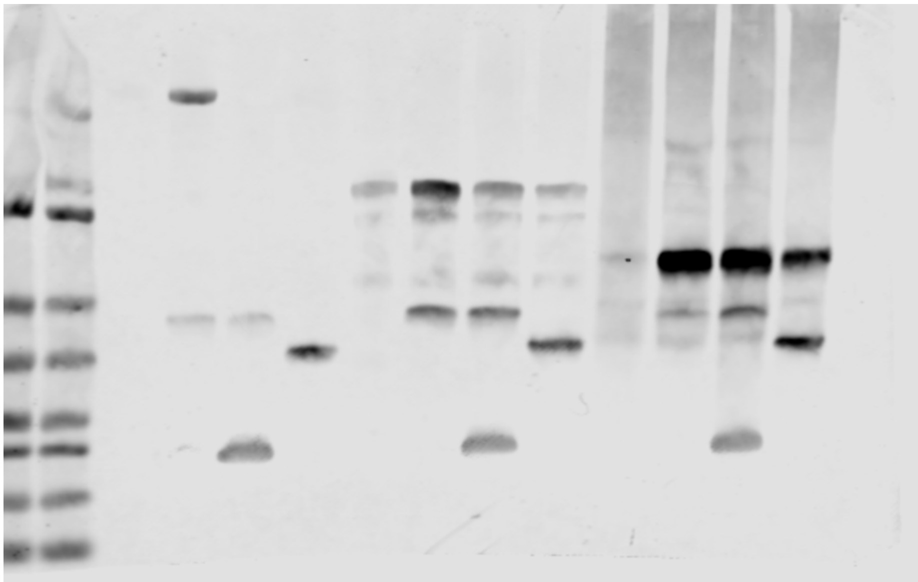

Total Protein

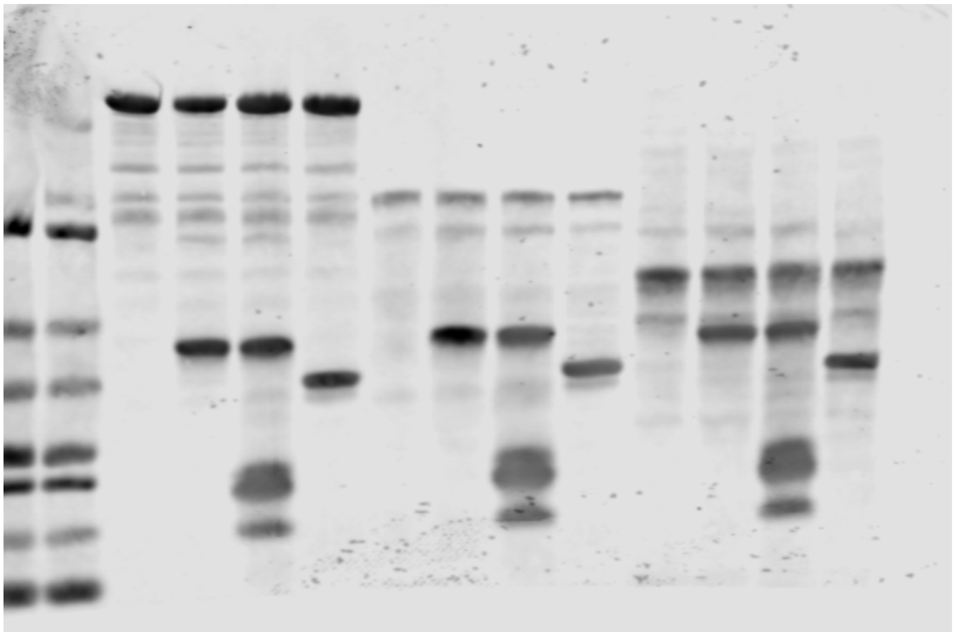

Figure S2 A:

Displayed Image:

Anti-H3K9ac

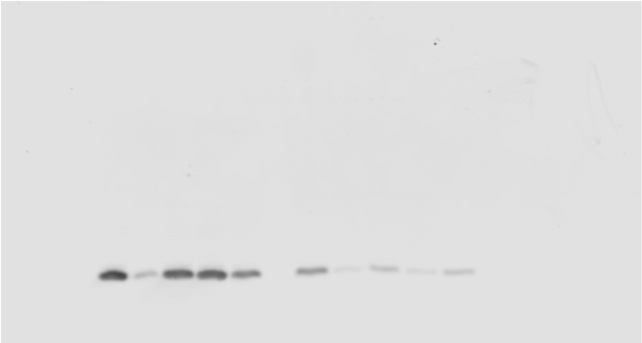

Anti-6XHis

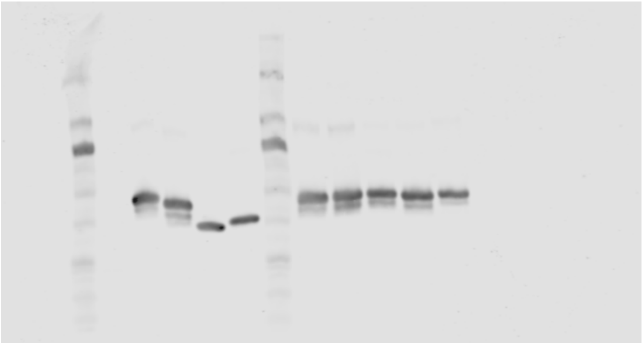

Figure S2 A:

Rep 2:

Anti-H3K9ac

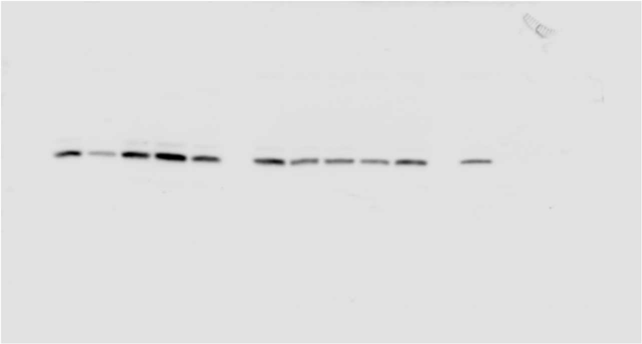

Anti-6XHis

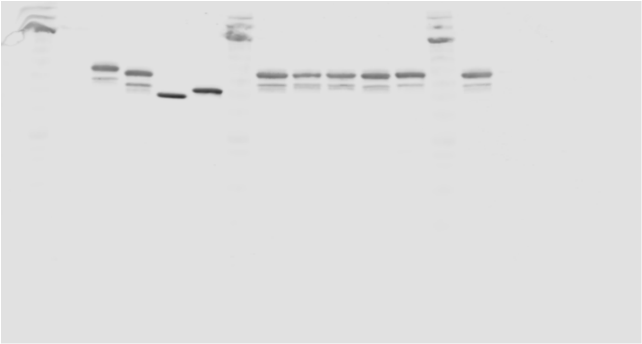

Figure S2 C:

Displayed Image:

EtBr Stain

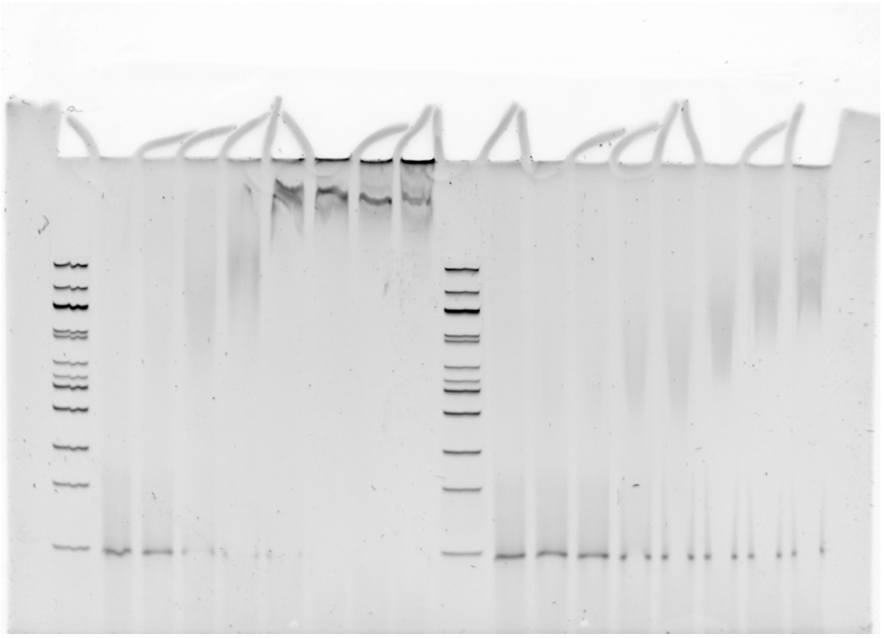

Figure S2 C:

Rep 2:

EtBr Stain

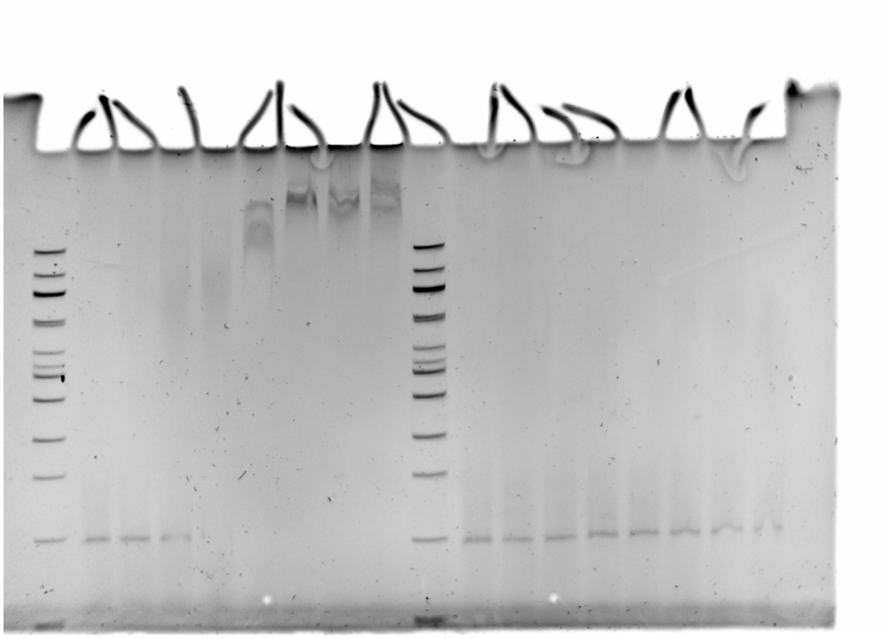

Figure S2 C:

Rep 3:

EtBr Stain

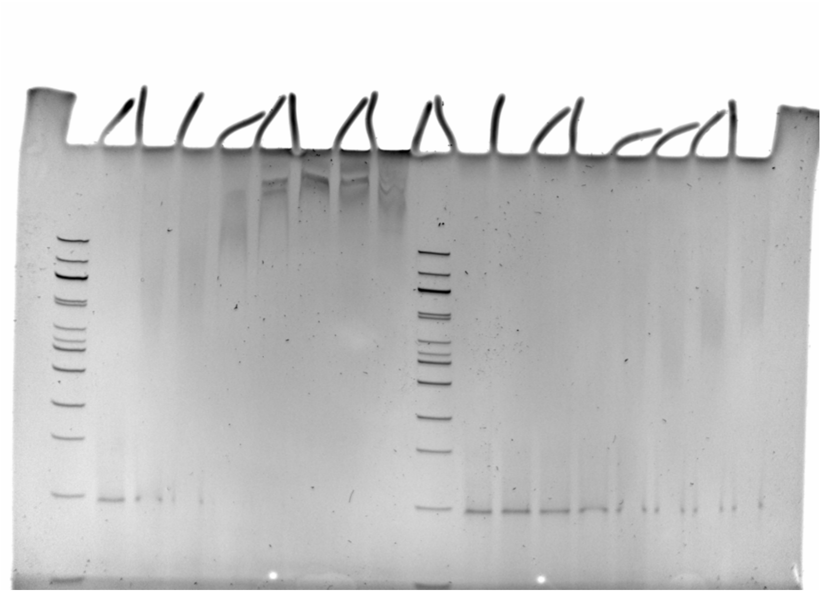

Figure S2 D:

Displayed Image:
Anti-ADPr

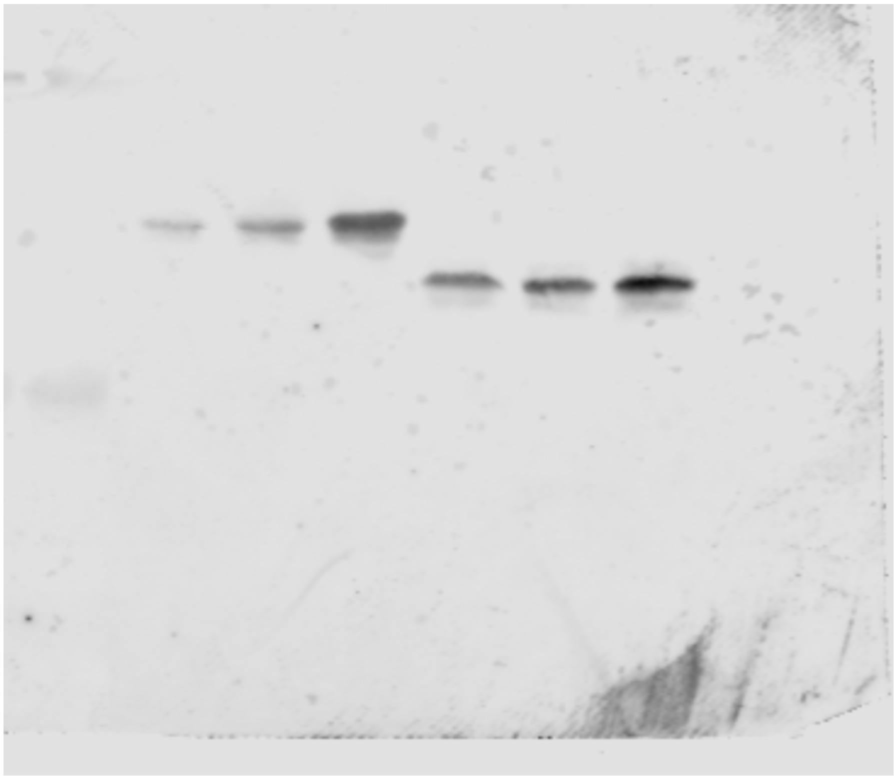

Total Protein

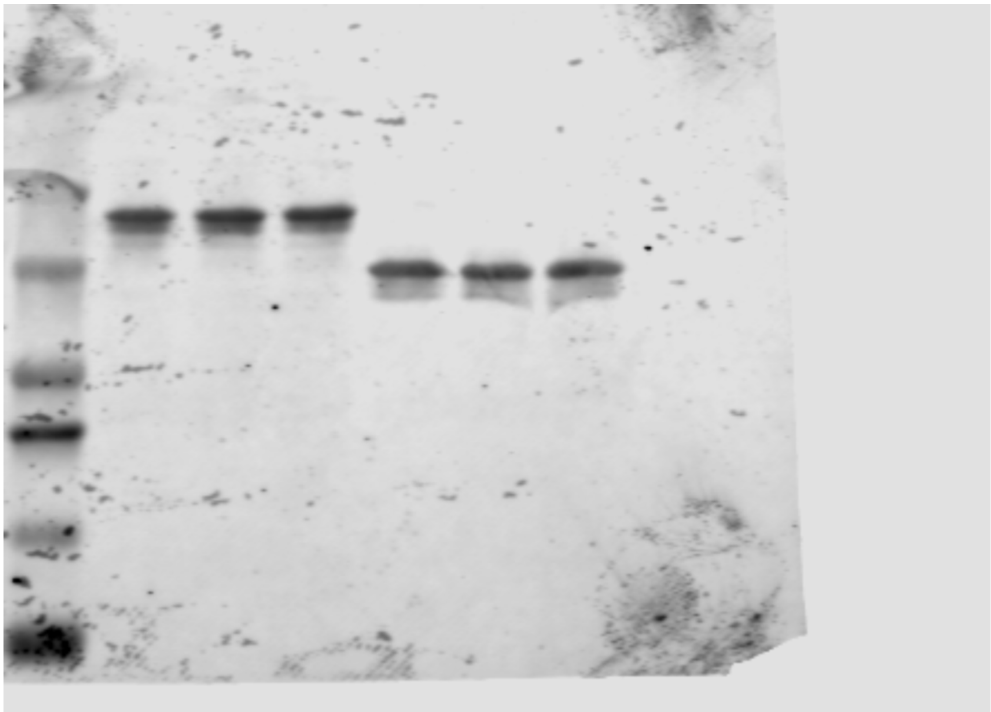

Figure S2 D:

Rep 2:
Anti-ADPr

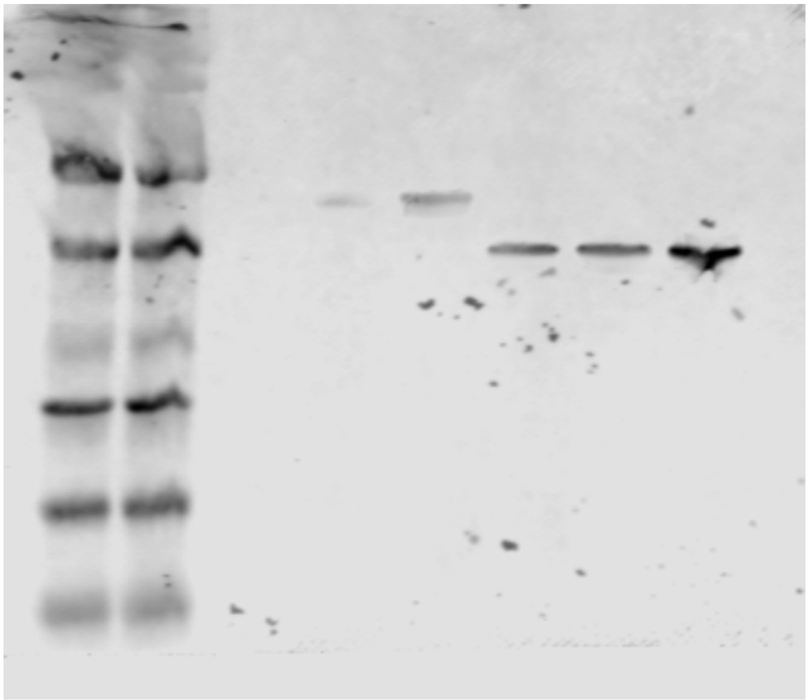

Total Protein

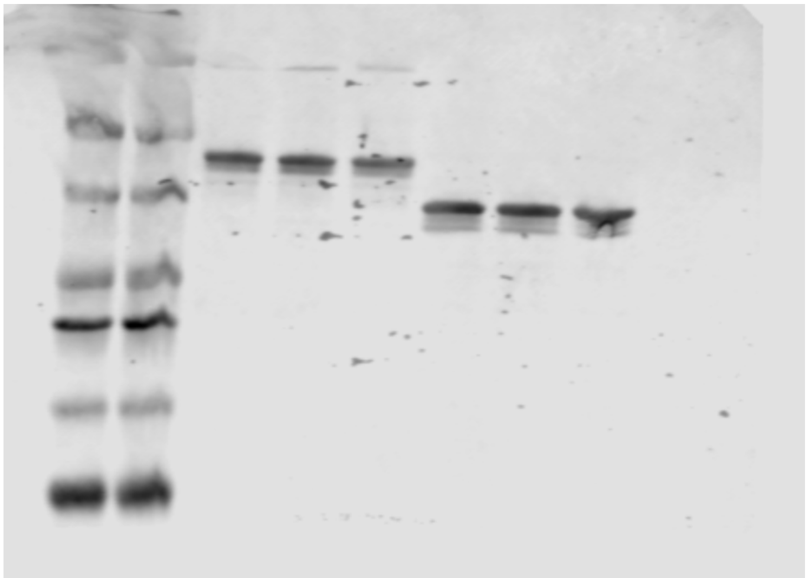

Figure S2 E:

Displayed Image:

Anti-ADPr

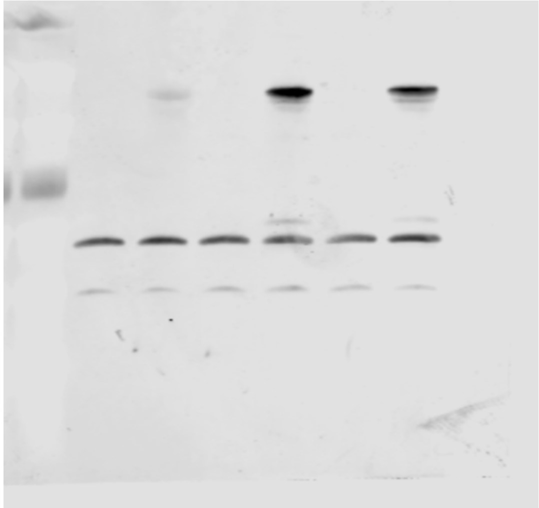

Anti-H3

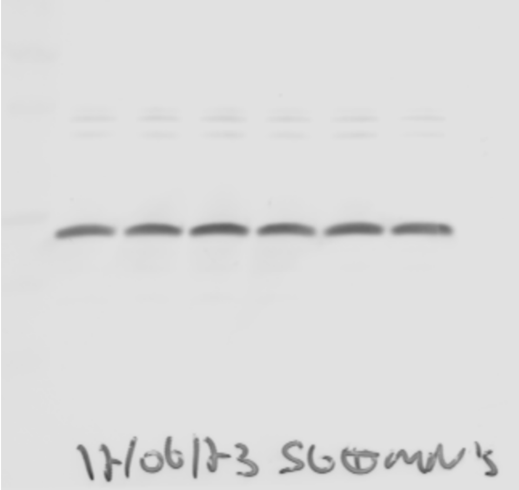

Total Protein

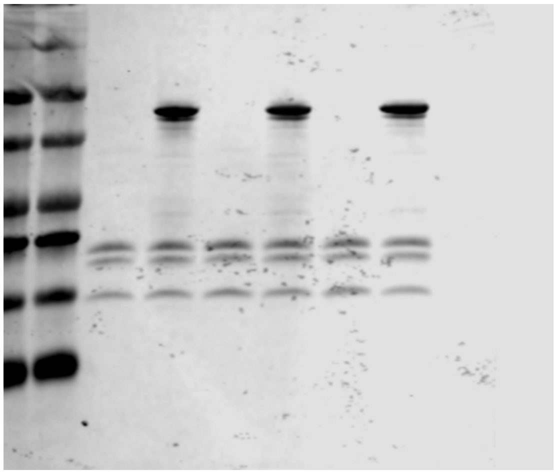

Figure S2 E:

Rep 2:

Anti-ADPr

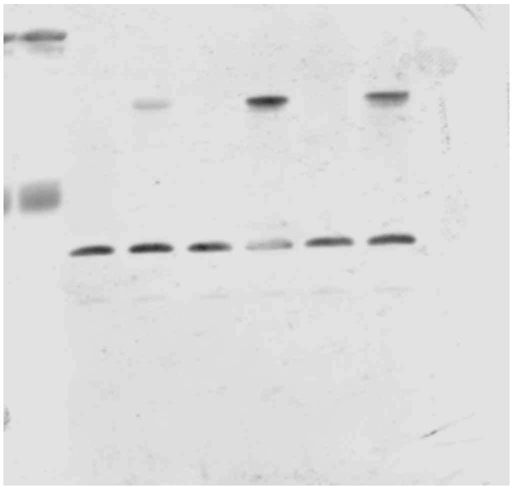

Anti-H3

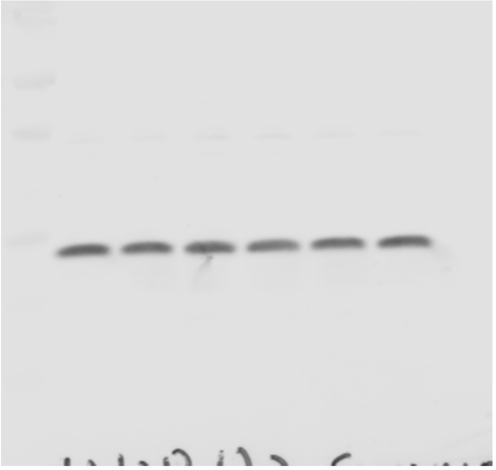

Total Protein

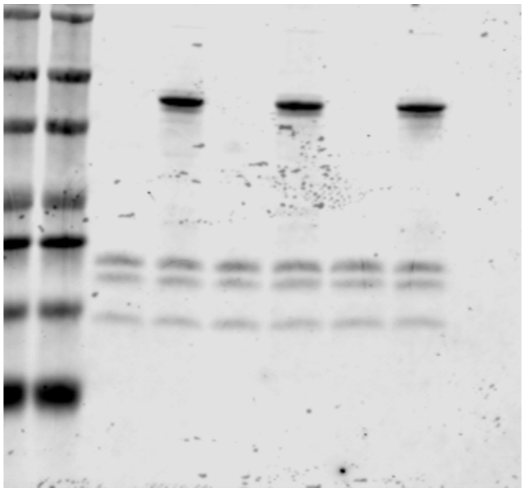

Figure 3 A:

Main Text:

Anti-ADPr

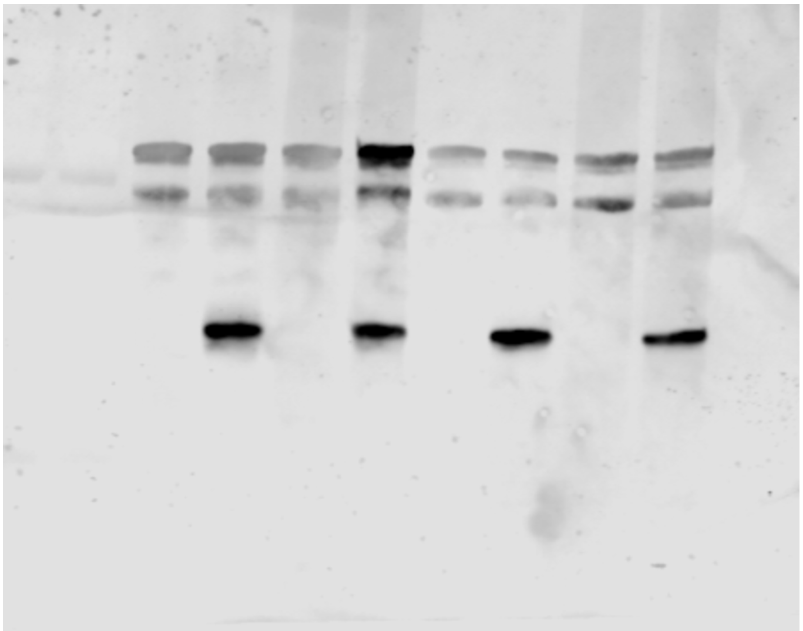

Total Protein

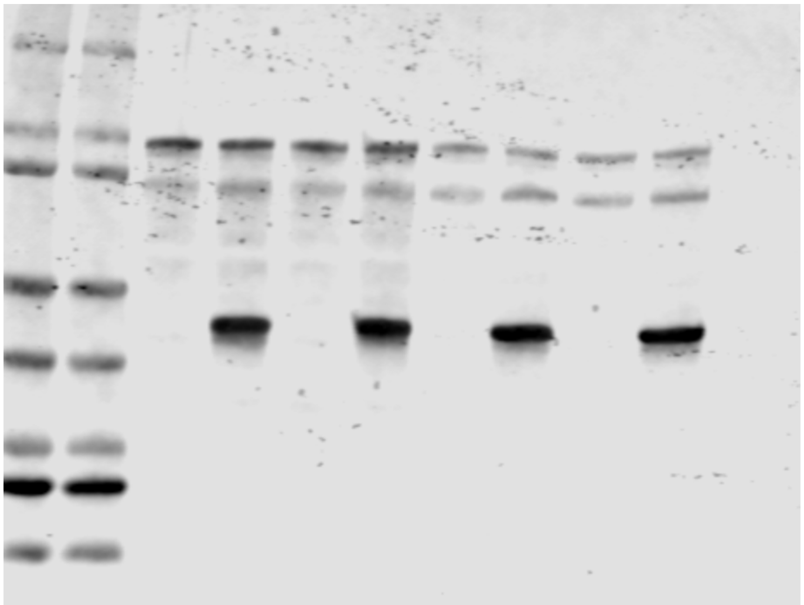

Figure 3 A:

Rep 2:

Anti-ADPr

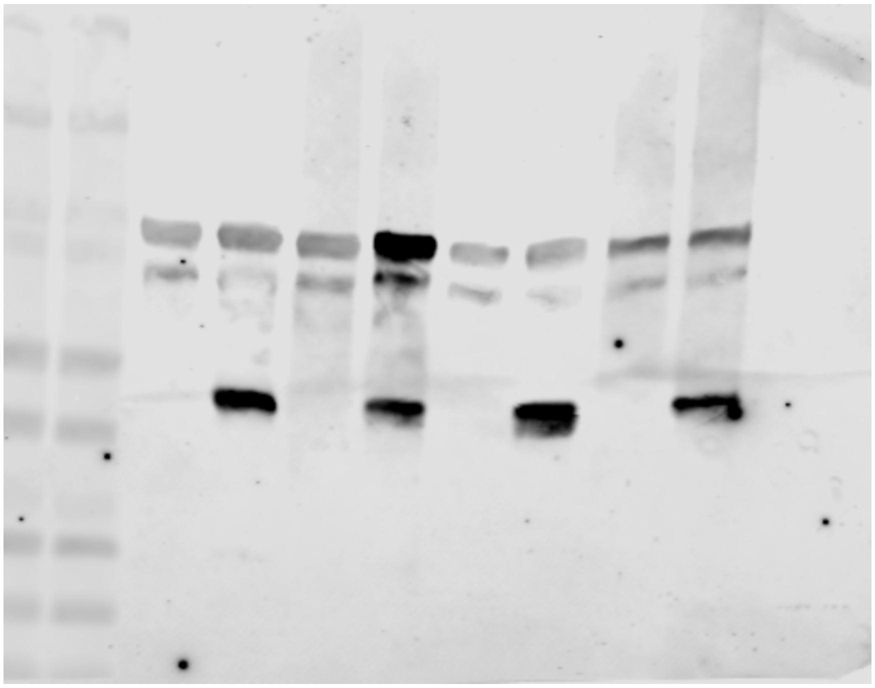

Total Protein

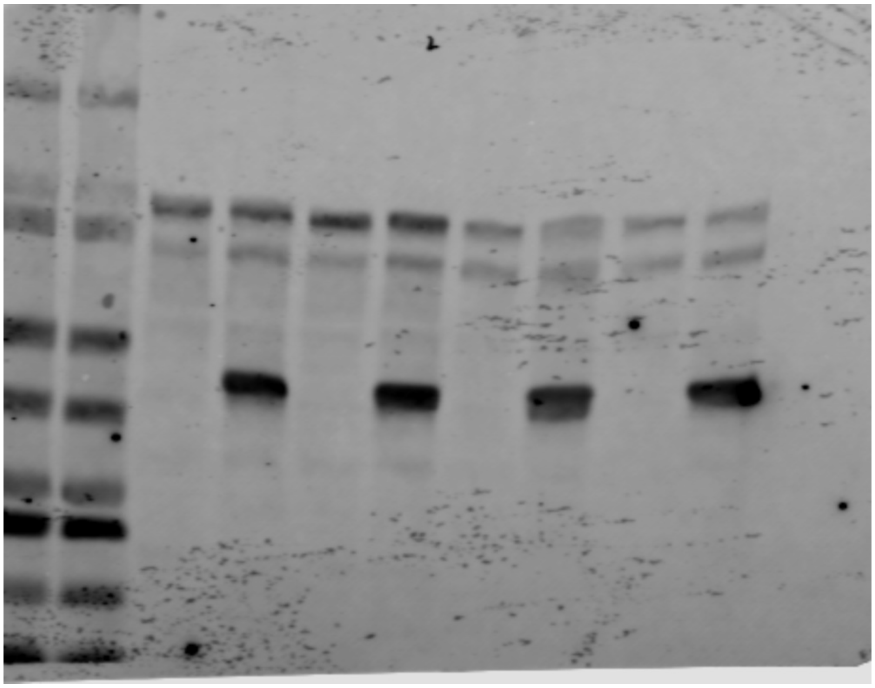

Figure 3 B:

Main Text:

Anti-ADPr

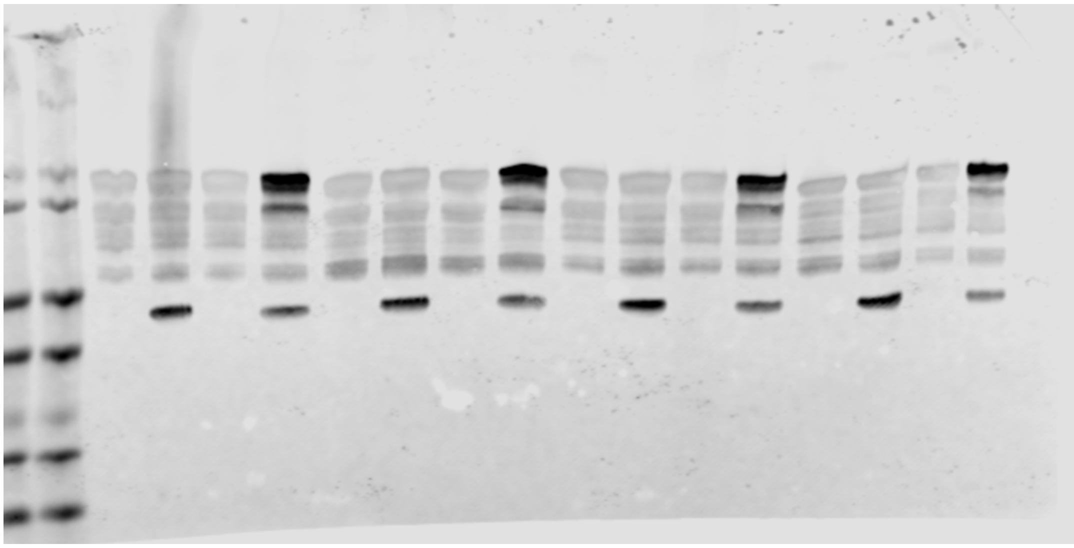

Total Protein

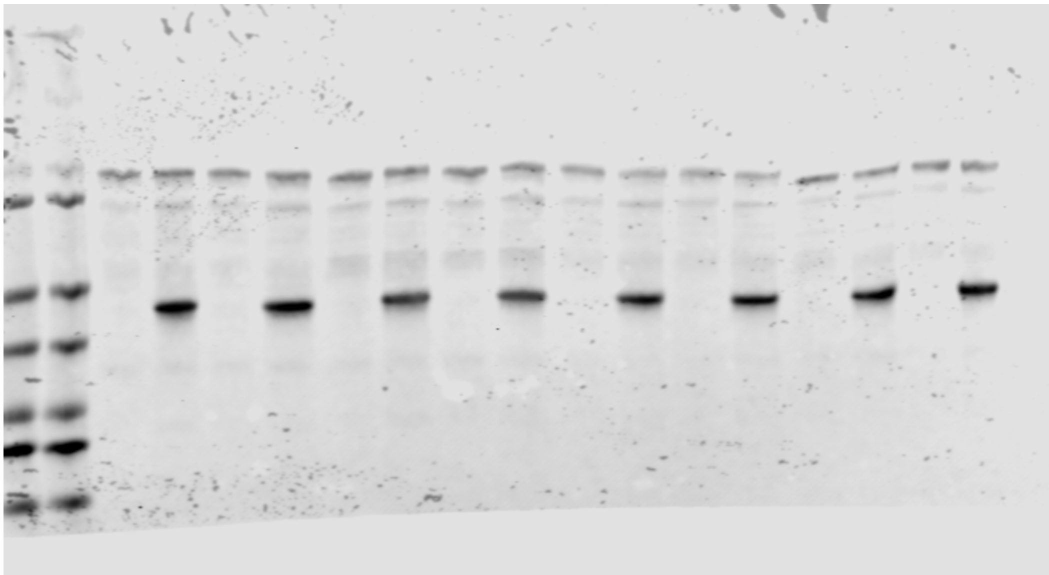

Figure 3 B:

Rep 2:

Anti-ADPr

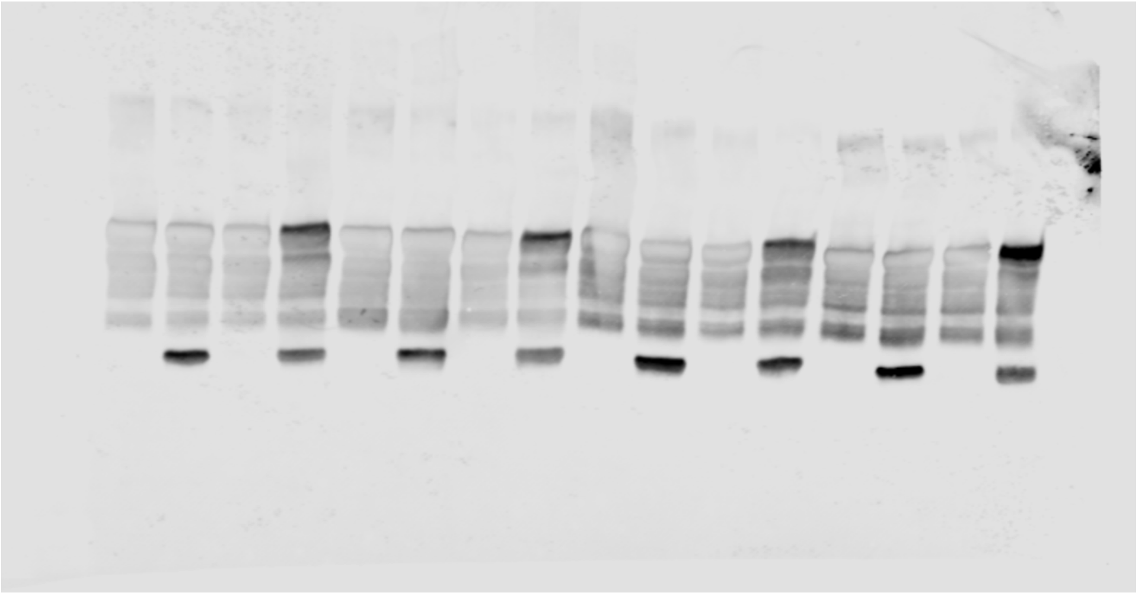

Total Protein

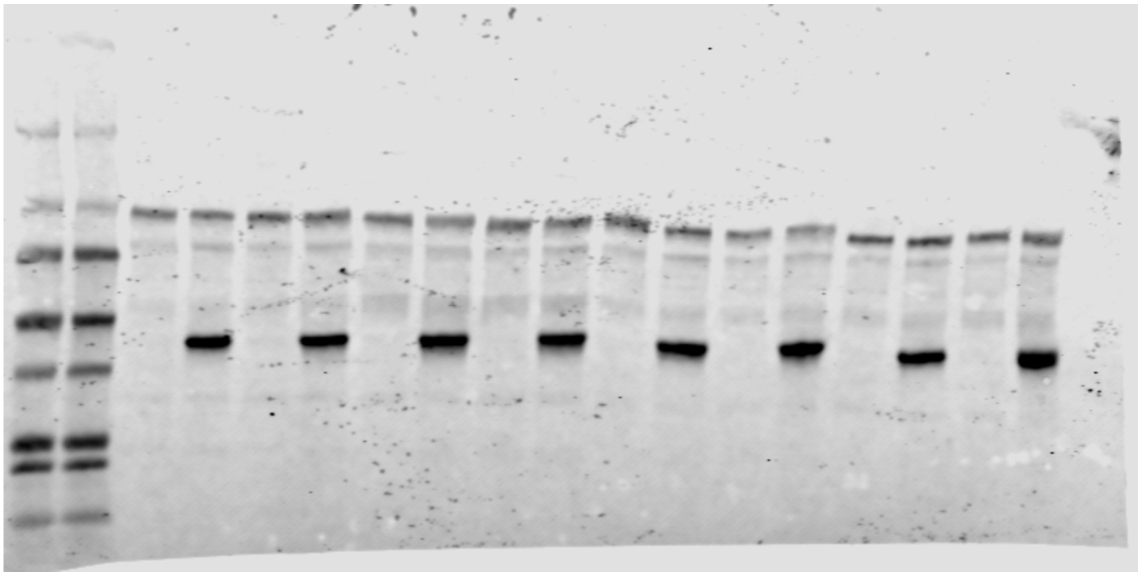

Figure 3 C:

Main Text:

Anti-ADPr

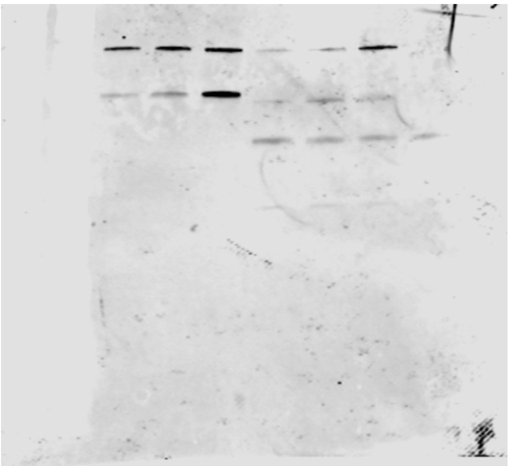

Anti-6XHis

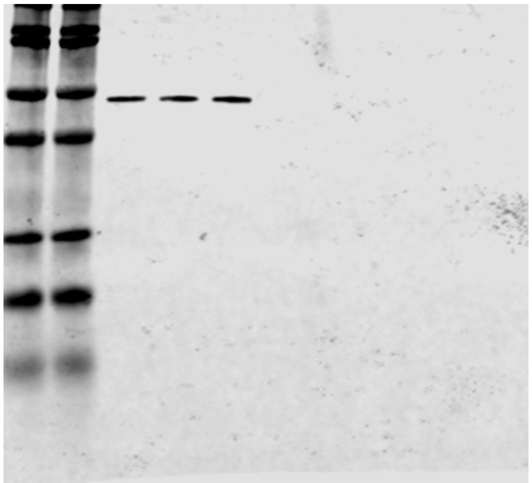

Anti-SIRT6

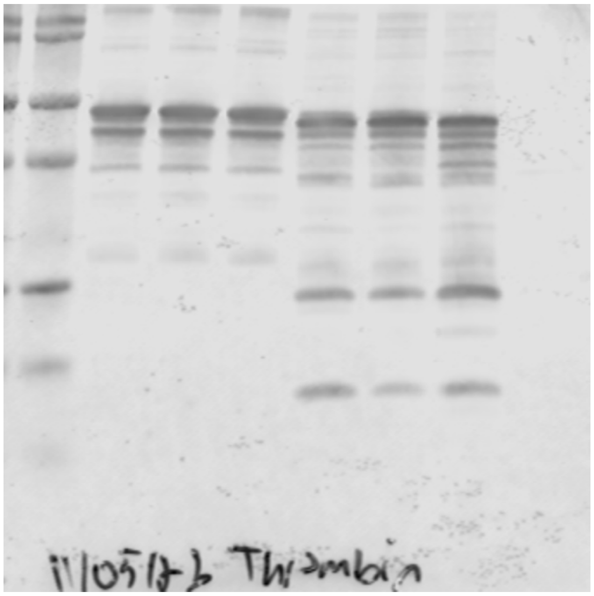

Figure 3 C:

Rep 2:

Anti-ADPr

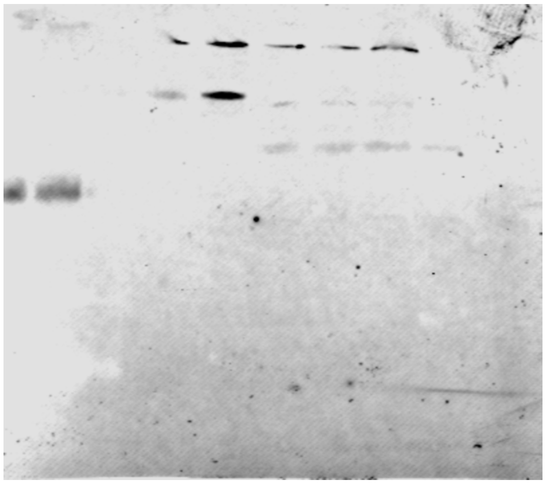

Anti-6XHis

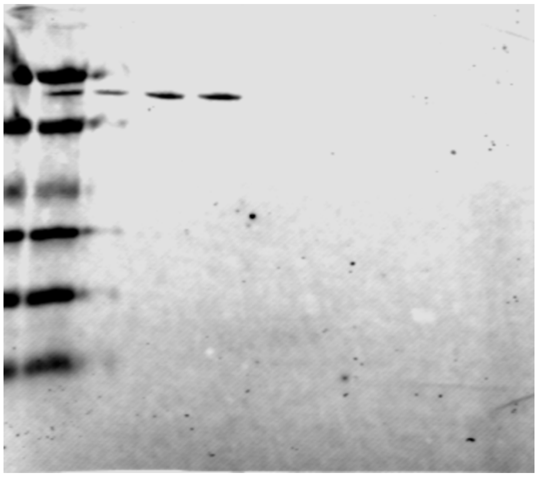

Anti-SIRT6

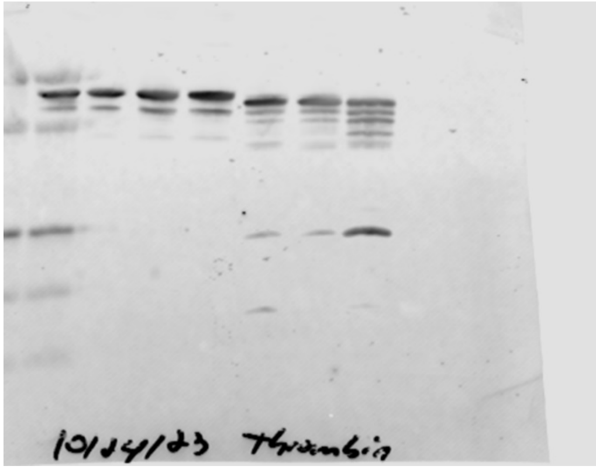


Figure S3 A:

Displayed Image:

Anti-ADPr

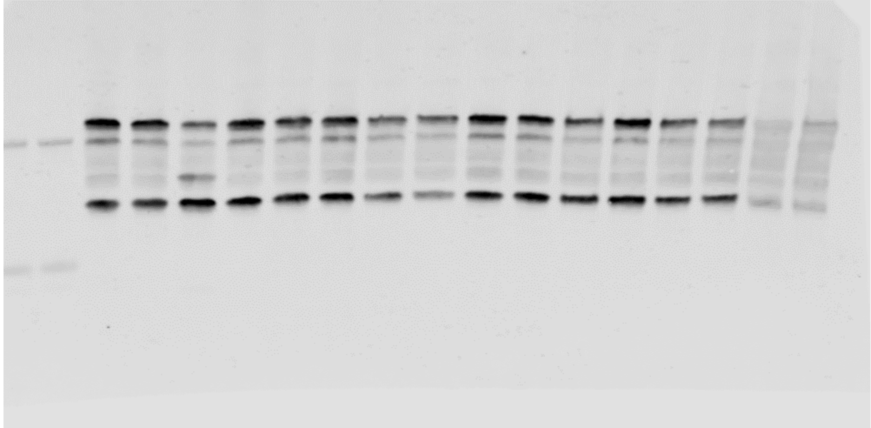

Total Protein

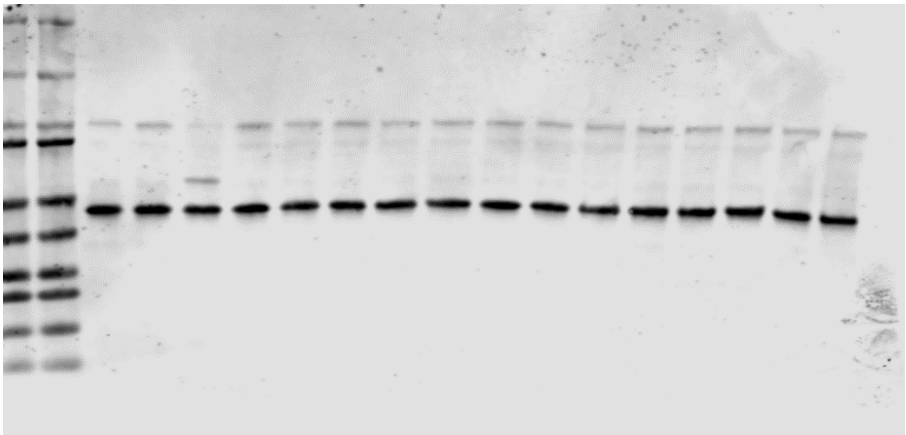

*For both blots, the right half (8 lanes) is the experiment referenced in the manuscript.

Figure S3 A:

Rep 2:

Anti-ADPr

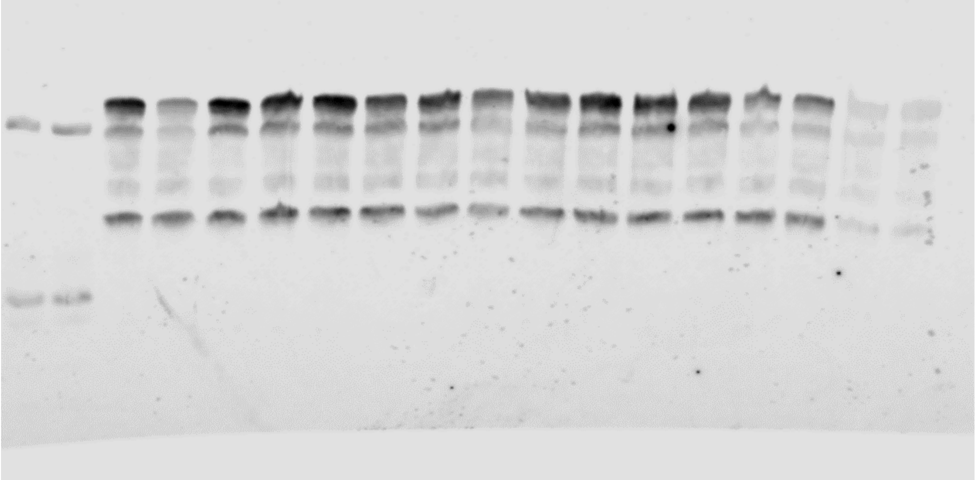

Total Protein

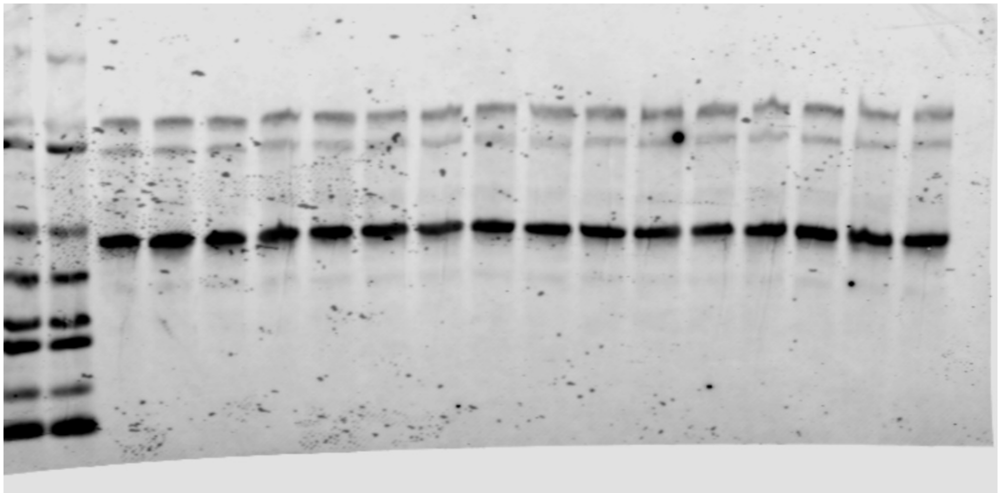

*For both images, the right half (8 lanes) is the experiment referenced in the manuscript.

Figure S3D

Main Text:

Anti-ADPr

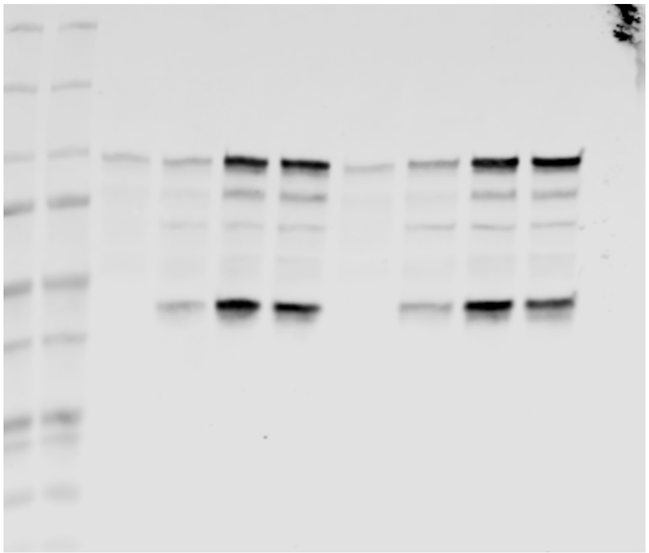

Total Protein

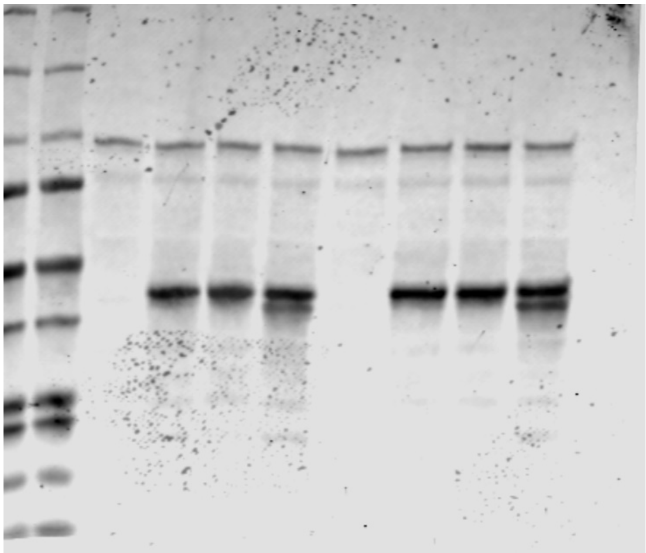

*Band below SIRT6 in reaction 4 and 8 is ARH3

Figure S3D:

Rep 2:

Anti-ADPr

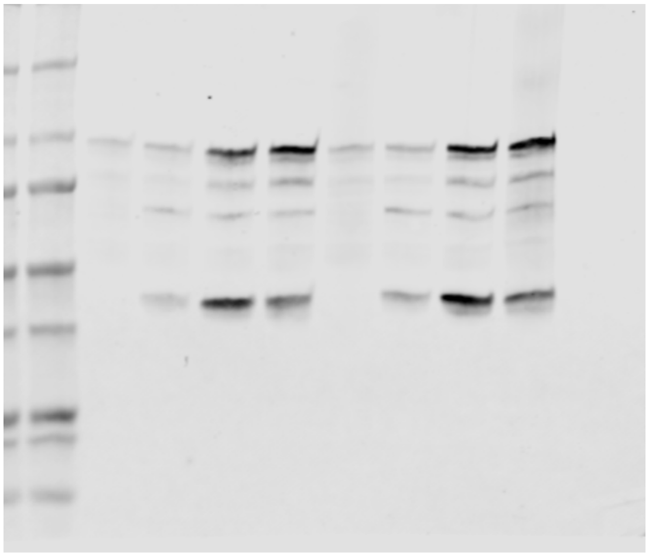

Total Protein

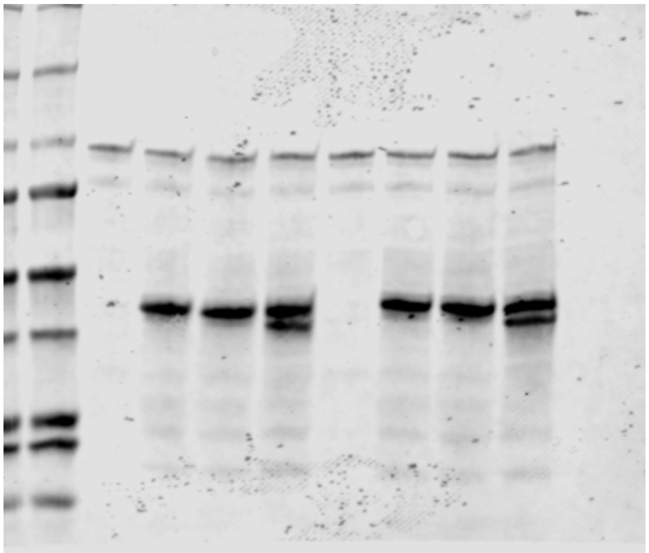

*Band below SIRT6 in reaction 4 and 8 is ARH3

Figure S4.

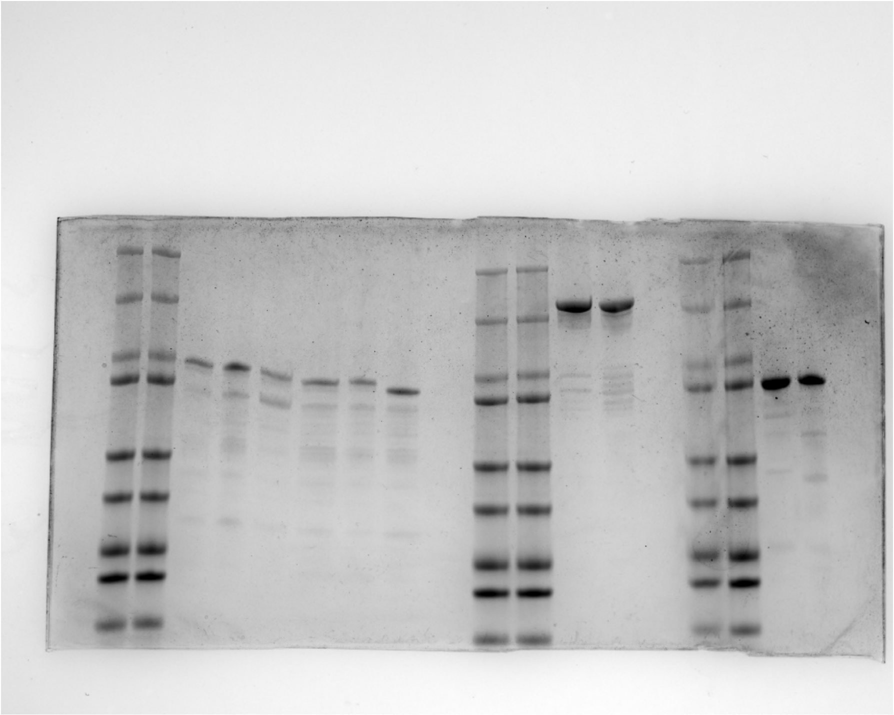

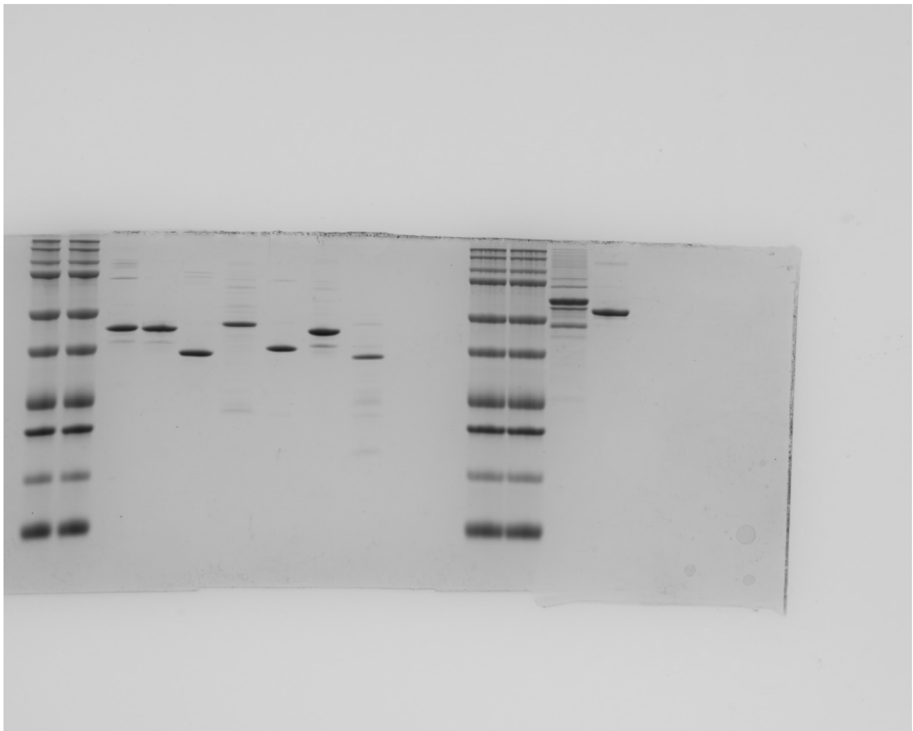

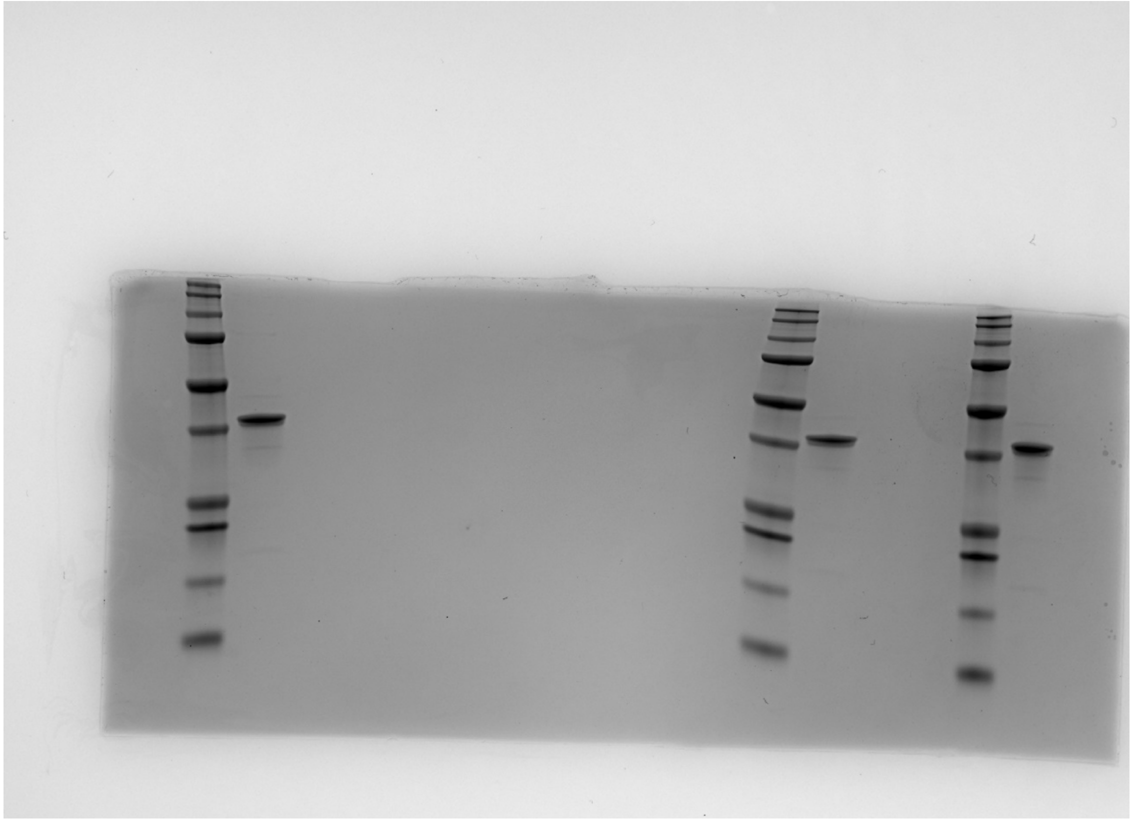

Figure S6B.

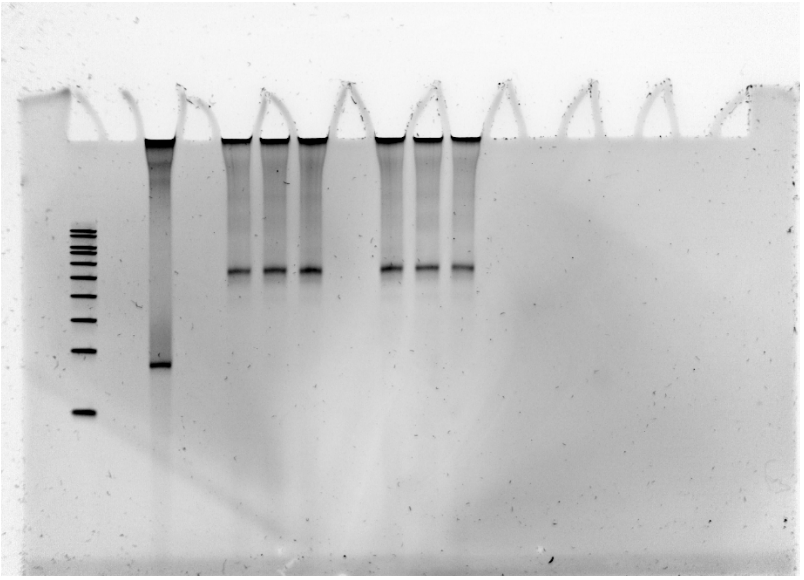

*The samples to the right are other batches of WT MNs

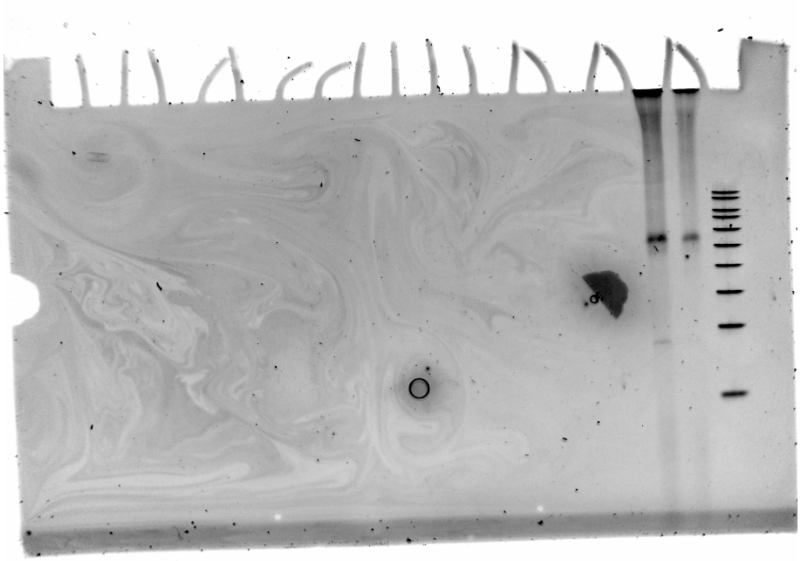

*The sample to the left is the same H3K9ac MN sample with a higher loading.
